# Enhancer-targeted CRISPR-A rescues haploinsufficiency and mutant phenotypes in organoid models of autism

**DOI:** 10.1101/2024.03.13.584921

**Authors:** George T. Chen, Gayatri Nair, Aubrey J. Osorio, Sandra M. Holley, Kimiya Ghassemzadeh, Yixuan Zhou, Jose Gonzalez, Jacqueline M. Martin, Congyi Lu, Neville E. Sanjana, Carlos Cepeda, Daniel H Geschwind

**Affiliations:** Program in Neurogenetics, Department of Neurology, David Geffen School of Medicine, UCLA, Los Angeles, CA; Center for Autism Research and Treatment, Semel Institute, UCLA, Los Angeles, CA; Intellectual and Developmental Disabilities Research Center (IDDRC), Department of Psychiatry and Behavioral Sciences; David Geffen School of Medicine, UCLA, Los Angeles, CA; Department of Human Genetics, David Geffen School of Medicine, UCLA, Los Angeles, CA; Institute of Precision Health, David Geffen School of Medicine, UCLA, Los Angeles, CA; New York Genome Center, New York, NY; Department of Biology, New York University, New York, NY

## Abstract

Autism Spectrum Disorder (ASD) is a highly heritable condition with diverse clinical presentations. Approximately 20% of ASD’s genetic susceptibility is imparted by *de novo* mutations of major effect, most of which cause haploinsufficiency. We mapped enhancers of two high confidence autism genes – *CHD8* and *SCN2A* and used CRISPR-based gene activation (CRISPR-A) in hPSC-derived excitatory neurons and cerebral forebrain organoids to correct the effects of haploinsufficiency, taking advantage of the presence of a wildtype allele of each gene and endogenous gene regulation. We found that CRISPR-A induced a sustained increase in *CHD8* and *SCN2A* expression in neurons and organoids, with rescue of gene expression levels and mutation-associated phenotypes, including gene expression and physiology. These data support gene activation via targeting enhancers of haploinsufficient genes as a therapeutic intervention in ASD and other neurodevelopmental disorders.

## Introduction

Autism Spectrum Disorders (ASD) are neurodevelopmental disorders characterized by impaired social behaviors and repetitive behaviors, often with severe functional impairment. ASD is highly prevalent, affecting approximately 1 in 54 individuals across all racial, ethnic, and socioeconomic groups, and thus presents a significant public health burden [Maenner et al., 2020]. Although it is highly heritable, a substantial fraction of genetic susceptibility, perhaps up to 20% [Gaugler et al., 2014], is imparted by *de novo* mutations with major effects (15-20%)[Geschwind, and State, 2015; Vorstman, and Scherer, 2023].

Studies in large cohorts of patients and their families – such as the Simons Simplex Collection (SSC), and the Autism Genetic Resource Exchange (AGRE) have identified hundreds of genes increasing autism risk with various levels of statistical support, most via harboring *de novo* mutations [Cirnigliaro et al., 2023; Fu et al., 2022; Iossifov et al., 2014; Ronemus et al., 2014; Ruzzo et al., 2019; Sanders et al., 2012; Satterstrom et al., 2020]. These “high-confidence” ASD genes (hcASD) are particularly enriched in genes that regulate gene expression or synaptic development during the peak of cortical neurogenesis [Parikshak et al., 2013; Satterstrom et al., 2020; Schafer et al., 2019]. Studies in model systems also strongly implicate haploinsufficiency as the underlying mechanism of mutation [Collins et al., 2020; Gompers et al., 2017; Huang et al., 2014; Platt et al., 2017; Tatsukawa et al., 2019]. This model is further supported by the observation that virtually all susceptibility genes are strongly depleted for heterozygous protein truncating variants in humans [Fu et al., 2022; Lek et al., 2016; Morrill, and Amon, 2019; Veitia, and Potier, 2015], which indicates that hcASD genes are highly dosage sensitive and require both alleles for normal function.

Here, we focused on two well-characterized hcASD risk genes, Chromodomain Helicase DNA-binding protein 8 (*CHD8*) and Sodium channel protein type 2 subunit alpha (*SCN2A*) [de la Torre-Ubieta et al., 2016; Rolland et al., 2023]. *CHD8* is strongly associated with ASD, and it appears to regulate many other hcASD risk genes involved in neurodevelopment and synaptic function [Bernier et al., 2014; Durak et al., 2016; Derafshi et al., 2022; Platt et al., 2017; Villa et al., 2020; Weissberg, and Elliott, 2021]. The *SCN2A* gene encodes the voltage-gated sodium channel Na_V_1.2 which is essential for the generation and propagation of action potentials in neurons, and is mutated in Dravet Syndrome and other sodium channelopathies [Ben-Shalom et al., 2017; Wolff et al., 2017]. The distinct roles of these two genes are evident in their normal patterns of expression in cortical development; *CHD8* peaks during neurogenesis and decreases over time, while *SCN2A* is expressed later, but still during mid-fetal development. Thus, in studying these two genes, we aimed to interrogate the efficacy of applying CRISPR-A during development.

CRISPR-A has effectively been shown to up-regulate target genes when directed to promoters, even rescuing neurodevelopmentally significant phenotypes [Chardon et al., 2024; Colasante et al., 2019; Colasante et al., 2020; Tamura et al., 2022]. One key issue in using CRISPR-A to compensate for haploinsufficiency is the potential deleterious effects on cell fitness caused by overexpression, which is a frequent consequence of targeting gene promoters [Kawamura et al., 2025; Matharu et al., 2018]. This is especially a concern with haploinsufficient genes, which are notoriously dosage sensitive [Fu et al., 2022; Huang et al., 2010] [Collins et al., 2022; Kawamura et al., 2025; Mao et al., 2023]. We therefore chose to target gene enhancers, with the rationale that endogenous gene expression regulation would be matched more closely across developmental time and cell type specific expression would be preserved [de la Torre-Ubieta et al., 2018; Won et al., 2019]. To date, no published studies targeting hcASD genes have used enhancer-targeted CRISPR-A, as few putative enhancers have been validated. We identified potential enhancers for *CHD8* and *SCN2A* predicted to be active during fetal brain development by leveraging our previous work identifying gene expression regulatory elements in human brain *in vivo* and *in vitro* [de la Torre-Ubieta et al., 2018; Won et al., 2019; Won et al., 2016]. We demonstrate that targeting these enhancer sequences significantly increased target gene expression, while maintaining the trajectory of expression across development in a cerebral organoid model system and in differentiating excitatory neurons. Core phenotypes, including over-proliferation in *CHD8*^+/-^ and electrophysiological abnormalities in *SCN2A*^+/-^ are also rescued by CRISPR-A for several months post-treatment. Taken together, our results indicate that enhancer-targeted CRISPR-A can rescue gene expression levels and mutant phenotypes, supporting the potential for using enhancer-targeted gene activation as a therapeutic approach in ASD.

## Results

### CHD8 haploinsufficiency leads to increased organoid size and over-proliferation of progenitor cell populations

We edited an embryonic stem cell line (HUES66, [Shi et al., 2023]) and an induced pluripotent stem cell line (KOLF 2.2J) using CRISPR to produce heterozygous knockouts (Methods). These lines were first characterized by establishing 2-dimensional excitatory neuron cultures using NGN2 induction (iNGN2) and 3-dimensional cerebral forebrain organoids (hCO; [Pașca et al., 2022]) to identify aberrant phenotypes to target for rescue (Figure 1A; Methods).

**Figure 1.**
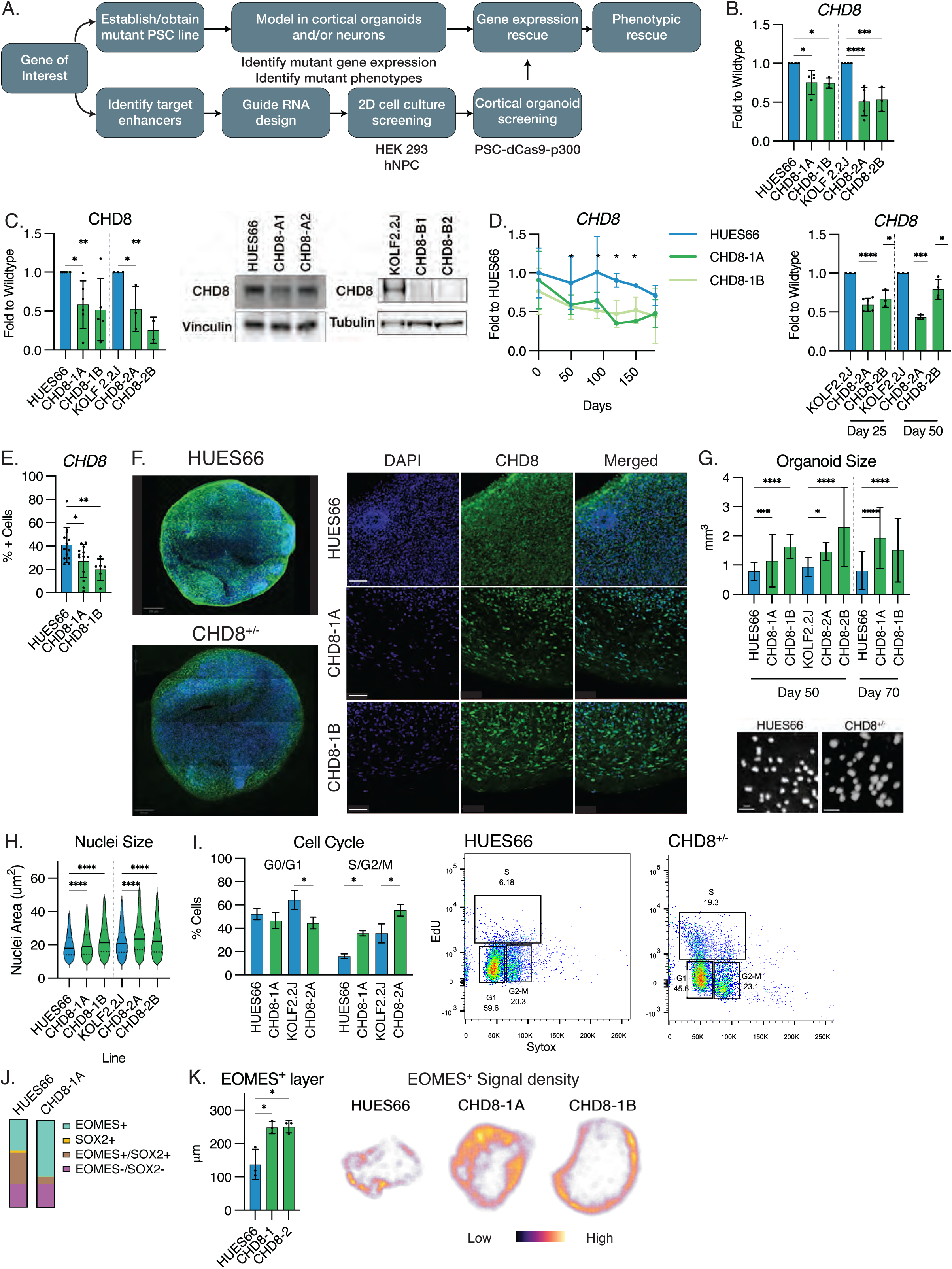
Characterization of CHD8 mutant stem cells and cortical organoids. (A) Overview of workflow for each gene of interest. (B) RNA expression of *CHD8* as measured by quantitative PCR shows expected reduction in mutant lines compared to wildtype control. N = 4 biological replicates, SD shown, * p < 0.05. (C) Protein expression of *CHD8* shows a 50% reduction in mutant lines compared to wildtype control. N = 6 biological replicates, SD shown, * p < 0.05. (D) Expression of *CHD8* mRNA by quantitative PCR in cortical organoids (hCO) from 0 to 180 days old shows a consistently lower level of *CHD8* across all timepoints measured. N = 6 biological replicates, SEM shown. Per replicate, 3 hCO are pooled and processed together. (E-F) immunofluorescence of CHD8 in 90-day old hCO sections shows lower levels of CHD8 in mutant hCO, quantified in (E). N = 7 biological replicates, SD shown, * p < 0.05. (G) hCO size measurements in 50- and 70-day old hCO show CHD8-mutant lines (CHD8-1A, CHD8-1B, CHD-2A, CHD8-2B) are significantly larger than wildtype (HUES66, KOLF 2.2J). N = 120 hCO, SEM shown. (H) Nuclei size of cells in hCO sections shows significantly larger nuclei in *CHD8*^+/-^ hCO. N = 120 hCO, * p < 0.05. (I) Flow cytometry of EdU- and Sytox-stained hCO shows an increase in cell proliferation in *CHD8* hCO. Quantitation of cell cycle state shows a significant difference in proportion of CHD8^+/-^ cells in active division compared to wildtype hCO at matching timepoints. N = 9 biological replicates, SD shown, * p < 0.05. Per replicate, 5 hCO per cell line were dissociated together for flow cytometry. (J) Expression of cell type markers EOMES and SOX2 in CHD8^+/-^ and wildtype hCO shows a larger proportion of EOMES+ cells in mutant hCO. N=6 (K) EOMES+ area in hCO sections shows an expanded EOMES+ layer thickness in CHD8-mutant hCO. N = 3, * p < 0.05. Sample density maps of hCO sections; brighter colors indicate higher Tbr2+ signal. All significance values calculated using Student’s two-tailed T-test.

Starting with the *CHD8*-mutant lines, we first confirmed the presence of the edit in the genomic DNA (Supplemental Figure 1A & B). We expect that the mutation of a single allele would lead to an approximately 50% reduction in mRNA and protein levels compared to the parental wildtype (HUES66 or KOLF2.2J), which we observed in both *CHD8*-mutated lines (CHD8-1A and CHD8-1B from HUES66, CHD8-2A and CHD8-2B from KOLF2.2J; Figure 1B, 1C). We next established cortical organoids (hCO) from all of the lines, using a protocol that was previously shown to accurately recapitulate many features of *in vivo* cortical development [Gordon et al., 2021; Pasca et al., 2015]. *CHD8* mRNA expression was consistently decreased in mutant hCO compared to the parental line in both backgrounds (Methods; Figure 1D) and was also reflected in a decrease in protein level as measured by immunofluorescence (Figure 1E&F). The expression trajectory of *CHD8* reflects *in vivo* patterns observed in human cortical development, peaking in the first and early second trimesters and decreasing gradually past late childhood (Supplemental Figure 2).

One of the most well-characterized phenotypes resulting from *CHD8* haploinsufficiency in affected individuals is macrocephaly [Bernier et al., 2014]. We therefore measured hCO sizes at multiple timepoints and found that at both 50- and 70-days, hCO produced from *CHD8* mutant lines were significantly larger than wildtype (Figure 1G). To determine if this size difference was the result of increased proliferation, increased cell size, or a combination of the two, we first measured the size of cell nuclei as a proxy for cell size and found a small, but significant increase in nuclear size in the HUES66-derived CHD8 mutant lines compared to the parental line (Figure 1H). To measure changes in proliferation, we performed cell cycle flow cytometry on dissociated hCO (Methods). We observed an increase in the proportion of cells in the S/G2/M phases (Figure 1I) in mutant organoids. Additionally, we labeled the dissociated cells with antibodies for SOX2, EOMES (TBR2), and RBFOX3 (NeuN) as markers for neural stem cells, neural progenitors, and neurons, respectively. Of the cells in the S/G2/M phases as identified by EdU, we found a larger proportion of EOMES^+^ cells in mutant hCO (Figure 1J), suggesting that an over-proliferation of neural progenitor cells also contributes to the macrocephaly phenotype. We confirmed this by immunofluorescence, observing a broadening of the EOMES^+^ cell layer in CHD8-mutant hCO, indicating an expansion of the neural progenitor population (Figure 1K). This agrees with published work from other groups [Paulsen et al., 2020] which used the same HUES66 CHD8^+/-^ lines, but a different differentiation protocol, as well as data in animal models [Nord et al., 2025]. Specifically, scRNAseq of three-month old hCO showed a decrease in excitatory neurons in CHD8^+/-^ organoids compared to wildtype and an increase in multiple progenitor cell populations, including cells from glial lineages (Supplemental Figure 3).

To further characterize the effects of CHD8 deficiency on development of hCO, we measured the expression of additional cell type markers and found no significant differences in stem cell, radial glia, and mature neuron marker expression between CHD8^+/-^ and wildtype hCO (Supplemental Figure 4A). We measured co-expression of CHD8 with SOX2 and TBR2 as markers for stem cells and intermediate progenitors, respectively and found significantly more co-expression of CHD8 and TBR2 in CHD8^+/-^ hCO sections than wildtype (Supplemental Figure 4B), reinforcing our finding that intermediate progenitors are an affected cell population in CHD8^+/-^ hCO. Lastly, we performed bulk RNAseq of HUES66 and CHD8^+/-^ hCO and found gene ontology terms for cell cycle and progenitor cells enriched in CHD8^+/-^ hCO, while oxidative phosphorylation and neuron projection terms were enriched in wildtype, consistent with our observations of a greater proportion of immature progenitor cells in the mutant hCO. Collectively, these results demonstrate that the hCO system recapitulates the patient macrocephaly phenotype, and that this enlargement is caused in part by an increase in cell size, but largely due to over-proliferation of progenitor cells early in development.

### SCN2A haploinsufficiency leads to aberrant excitatory neuron phenotypes

The HUES66 ESC line was originally edited to induce a heterozygous SCN2A mutation; SCN2A-1 [Lu et al., 2019]. We additionally received two iPSC lines established from patients with SCN2A mutations (SCN2A-2, SCN2A-3; Methods). The *SCN2A* mutation was first confirmed in the mutant stem cell lines by Sanger sequencing (Supplemental Figure 1C-D). We validated the effect of this mutation on expression levels of *SCN2A* mRNA and protein (Figure 2A, 2B). Unlike the *CHD8*^+/-^ lines, in the hCO from *SCN2A*^+/-^ cells, we observed that *SCN2A* expression did not significantly diverge in expression levels compared to the wildtype hCO until after 120 days *in vitro* (Figure 2C). This observation is consistent with the expression of *SCN2A* in human cortical development, where *SCN2A* increases monotonically during fetal cortical development, reaching an asymptote around birth (Supplemental Figure 2;[Brunklaus, and Lal, 2020]). Using immunofluorescence, we analyzed SCN2A protein expression levels in 90-, 150-, and 180-day old hCO. The percentage of SCN2A-positive cells was significantly decreased in mutant organoids at each measured timepoint and the largest difference in expression was observed between 150-180 days (Figure 2D, 2E).

**Figure 2.**
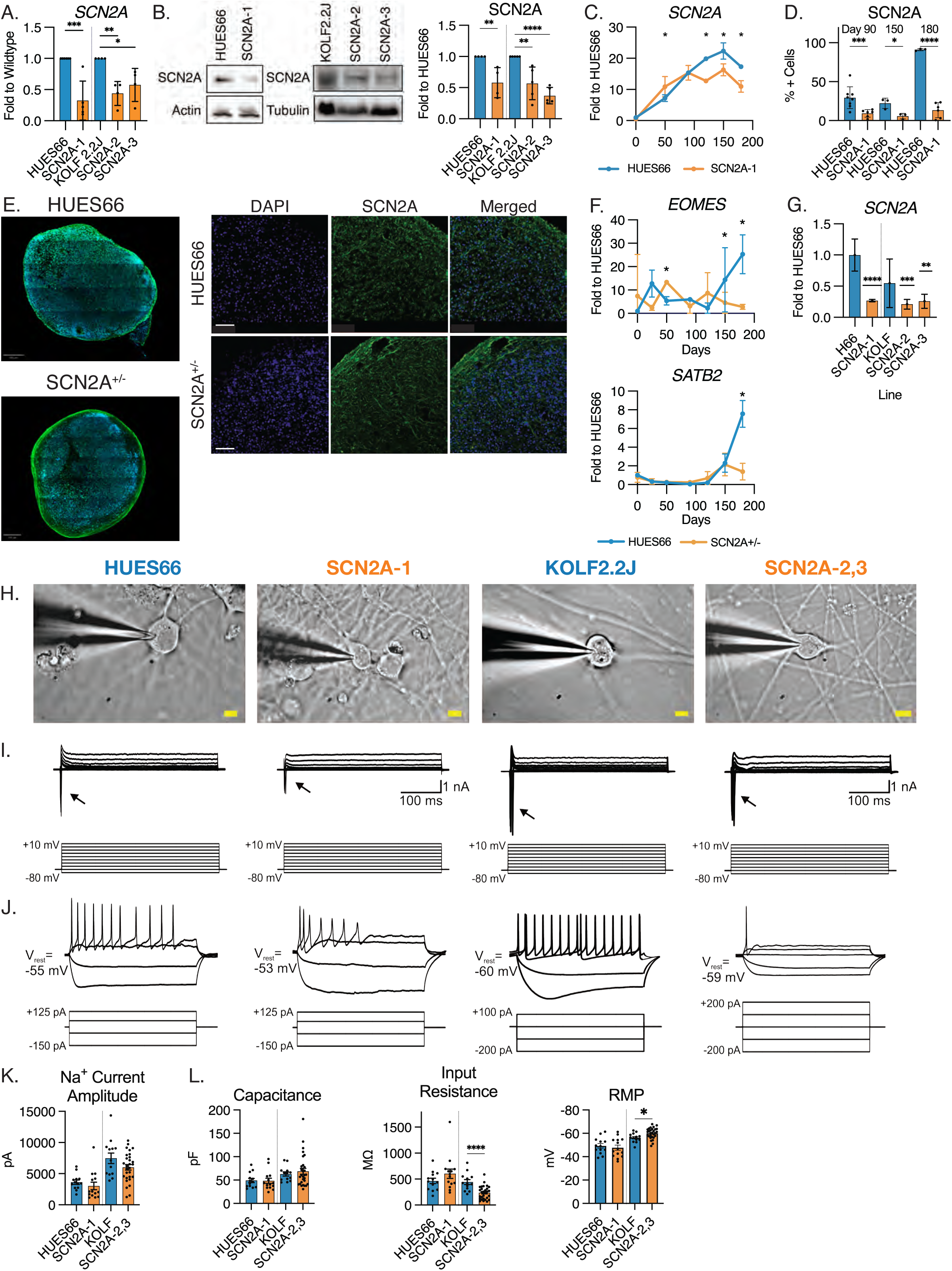
Characterization of SCN2A mutant stem cells and cortical organoids. (A) RNA expression of *SCN2A* shows expected 50% reduction in mutant lines (SCN2A-1, SCN2A-2, SCN2A-3) compared to wildtypes (HUES66, KOLF2.2J). N = 6, SD shown, * p < 0.05. (B) Protein expression of SCN2A shows a 50% reduction in mutant lines compared to controls. N = 6, SD shown, * p < 0.05. Full blot images in Supplement. (C) Expression of SCN2A in hCO from 0 to 180 days old shows SCN2A expression only decreased in mutant hCO after day 100. N = 6, SEM shown. (D-E) Immunofluorescence of SCN2A in 180-day old hCO sections shows lower levels of SCN2A in mutant hCO, quantified in (D). N= 6, SD shown, * < 0.05. (F) Expression of EOMES and SATB2, as markers for intermediate progenitor cells and upper layer neuron cells, respectively, are decreased in SCN2A-mutant hCO compared to wildtype. N = 6, SD shown. (G) Expression of *SCN2A* in iNGN2 neurons from wildtype and mutant lines. N=6, SEM shown, * p < 0.05. (H) Infrared differential interference contrast (IR-DIC) images of wildtype (HUES66, KOLF2.2J) and mutant lines (SCN2A-1^+/-^ iNGN2 and pooled SCN2A-2^+/-^ and SCN2A-3^+/-^ neurons during patch-clamp recordings. Scale bar = 10 µm. (I) Sample recorded traces in voltage-clamp mode from HUES66 and SCN2A-mutant expressing cells in response to depolarizing voltage steps (10 mV steps from -80 mV to +10 mV). Arrow indicates the presence of transient Na+ currents that are visible starting at ∼-40 mV. (J) Sample recorded traces of wildtype control and mutant cell current responses in current clamp mode following hyperpolarizing and depolarizing current injections. (K) Summary graphs of Na^+^ current amplitudes. (L) Summary graphs of passive cell membrane properties. Plots show mean values ±SEM. N=13 (HUES66), N=14 (SCN2A-1^+/-^), N=13 (KOLF2.2J), and N=29 (pooled SCN2A-2^+/-^ and SCN2A-3^+/-^). All significance values calculated using Student’s two-tailed T-test.

We next analyzed the expression of *EOMES* and *SATB2* as markers of intermediate progenitor cells and upper layer neurons, respectively, cells that are well-represented at this time. Mirroring the trajectory of *SCN2A* expression, both *EOMES* and *SATB2* showed significant decreases in expression only at later timepoints (Figure 2F). By qPCR and immunofluorescence, we performed additional characterization of hCO and found no significant differences in stem cell, radial glia, or neuron marker expression (Supplemental figure 4A). However, comparing SCN2A^+/-^ hCO to wildtype by bulk RNAseq showed significant enrichment in ontology terms for chromatin remodeling, neural stem cells, and neural progenitor cells in SCN2A^+/-^ hCO and enrichment in calcium signaling pathway, ion transport, and axoneme assembly in wildtype hCO (Supplemental Figure 4D), suggesting that the SCN2A-mutant hCO were less mature than wildtype.

We additionally established excitatory neuron cultures using *NGN2* overexpression [Zhang et al., 2013] and measured *SCN2A* expression after 35 days of differentiation, finding significant decreases in *SCN2A* expression in the mutant lines (Figure 2G). We then performed whole-cell patch-clamp electrophysiology on individual neurons (Figure 2H-L). Cells from all lines formed synaptic connections and exhibited spontaneous postsynaptic current (PSC) events of varying frequencies; however, PSC frequencies were highly variable across cells within each line and did not differ significantly between genotypes (HUES66: mean = 3.07±0.87 Hz, range = 0.60-9.80 Hz; KOLF2.2J: 1.43±0.66 Hz, range = 0.02-7.88 Hz; *SCN2A-1^+/-^*: mean = 3.91±2.68 Hz, range = 0.04-33.07 Hz; pooled SCN2A-2^+/-^ and SCN2A-3^+/-^: 2.25±0.40 Hz; pooled). This variability may reflect, in part, differences in local synaptic network formation that can arise with modest variations in cell confluence in NGN2-derived cultures. While peak Na+ current amplitudes were lower in SCN2A-mutant neurons across all lines as expected (Figure 2I), these differences did not reach statistical significance (Figure 2K). Despite this, we observed a consistent reduced excitability in response to a sustained depolarizing current injection across all mutant lines compared with controls (Figure 2J, and Figure 6D, 6E). We additionally measured passive membrane properties and intrinsic electrophysiological measures, which exhibited line-dependent effects, with significant differences in input resistance (passive) and resting membrane potential (RMP) observed selectively in KOLF2.2J compared to the patient-derived SCN2A-2,3 lines (Figure 2L). These results show that *SCN2A* haploinsufficiency disrupts the development and function of excitatory neurons, consistent with what has been previously observed in both mouse models [Eaton et al., 2021; Tamura et al., 2022; Yang et al., 2023] and cultured neurons from affected individuals [Brown et al., 2024; Mao et al., 2023; Que et al., 2021].

### Enhancer-targeted guideRNA increase expression of CHD8 and SCN2A using CRISPR-A

We hypothesized that using CRISPR-A to increase expression of haploinsufficient genes would effectively rescue gene expression and subsequently, mutant phenotypes. We first identified enhancer regions to target (Methods) by combining previously published maps of fetal human brain genome using ATAC-seq [de la Torre-Ubieta et al., 2018] and Hi-C [Won et al., 2016] with a broader list of adult human brain enhancers generated by the PsychENCODE Consortium [Wang et al., 2018]. We focused primarily on fetal brain enhancers, as they were more likely to be relevant during the time points that we examined. Using these maps (Table 1, Supplemental Figure 5), we identified four putative human fetal brain enhancers for *CHD8* and three for *SCN2A*, which we subsequently filtered (Methods) down to two enhancers for CHD8 and one for SCN2A. Per enhancer, we synthesized three guide RNA, for a total of six *CHD8* gRNA candidates and three *SCN2A* gRNA candidates (Figure 3A).

**Figure 3.**
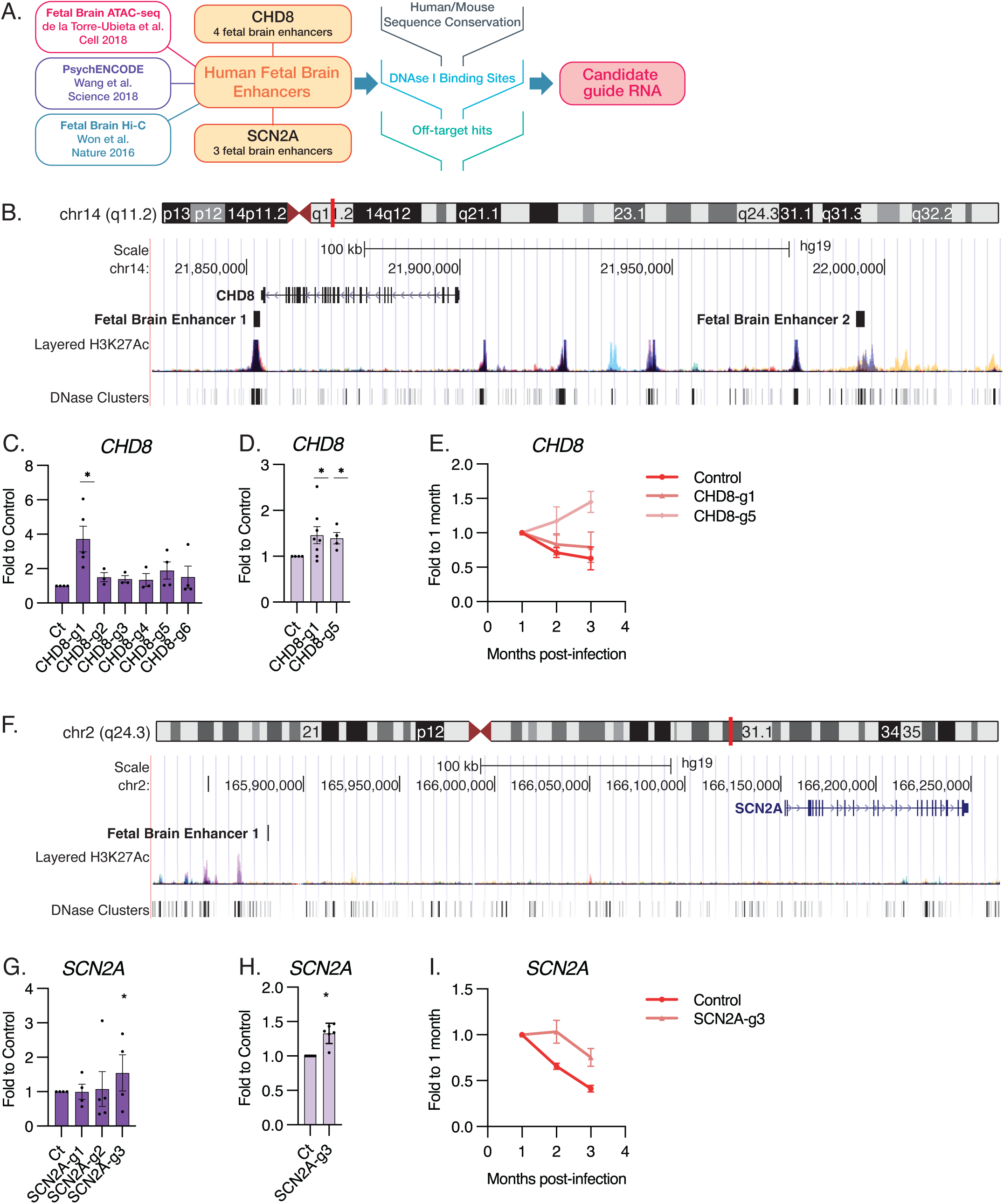
A stepwise approach to identifying and screening human fetal brain enhancer activity. (A) Workflow for identifying human fetal brain enhancers from previously published datasets and subsequent design and refinement of guide RNA sequences based upon sequence conservation, DNAse I binding, and off-target effects. (B) UCSC Genome browser window highlighting CHD8 in the genome and the two human fetal brain enhancers targeted in this study. (C) Screening *CHD8* guide RNA in HEK293T cells shows an increase in the mRNA expression of a gene of interest for at least one guide per identified enhancer. Expression is measured as fold over empty vector control. N = 4, SD shown, * p < 0.05. (D) Successful guide RNA from HEK293T assays were packaged into lentivirus and tested in primary human neural progenitor cells. All tested guide RNA viruses yielded a significant increase in expression of the gene of interest. Expression is measured as fold over empty vector control. N = 5, SD shown, * p < 0.05. (E) RNA expression of CHD8 in dCas9-p300 hCO is increased by guide RNA expression over empty vector control up to 3 months post-infection. Measurement of gene expression is relative to sample collected at 1-month post-infection. N = 3, SD shown. (F) UCSC Genome browser window highlighting SCN2A in the genome and the human fetal brain enhancer targeted in this study. (G) Screening *SCN2A* guide RNA in HEK293T cells shows an increase in the mRNA expression of a gene of interest. Expression is measured as fold over empty vector control. N = 4, SD shown, * p < 0.05. (H) SCN2A guideRNA significantly increased gene expression in human neural progenitor cells. N = 6, SD shown, * p < 0.05. (I) RNA expression of SCN2A in dCas9-p300 hCO is increased by guide RNA expression over empty vector control up to 3 months post-infection. Measurement of gene expression is relative to sample collected at 1-month post-infection. N = 3, SD shown. All significance values calculated using Student’s two-tailed T-test.

For *CHD8*, the two targeted enhancers flanked opposite ends of the gene (Figure 3B, full map in Supplemental Figure 5). We first screened the candidate gRNA in HEK293T cells, co-transfecting the guideRNA with a dCas9-p300 plasmid. We chose to use dCas9-p300 because it is an endogenous regulator of gene expression, as opposed to other synthetic activation constructs [Hilton et al., 2015]. Of the six *CHD8* guides tested, only two, CHD8-g1 and CHD8-g5, significantly increased *CHD8* expression in HEK293T cells, one for each of the enhancers (Figure 3C). As an additional validation step, we transduced the guides into primary human neural progenitor cells (phNPCs) to determine if the identified enhancers were active in a more physiologically relevant model system (Figure 3D). Both guides significantly increased *CHD8* expression in phNPCs. Finally, we screened the guides in hCO, using a stem cell line constitutively expressing dCas9-p300. We established dCas9-p300 hCO and cultured them for fifty days prior to infection with the guide RNA. Both guides increased *CHD8* expression over control-infected organoids (Figure 3E), indicating that a single treatment was durable across several months of development.

After filtering (Methods), only one human fetal brain enhancer was targeted for *SCN2A* activation (Figure 3F, full map in Supplemental Figure 5). Of the three guides screened in HEK293 cells, one, SCN2A-g3, produced a significant increase in SCN2A expression level (Figure 3G), which also significantly increased expression in phNPCs (Figure 3H). As a positive control, we also designed guide RNAs to target the promoter of SCN2A and found that SCN2A expression was significantly increased as well (Supplemental figure 6A). In dCas9-p300 organoids, we transduced the gRNA after fifty days in culture and collected organoids for 3 months post-infection. We observed an increase in *SCN2A* expression across all measured timepoints, validating the durability of the CRISPR-A treatment (Figure 3I).

### CRISPR-A treatment increases *CHD8* and *SCN2A* expression in haploinsufficient organoids and neurons

We next applied the CRISPR-A system to the haploinsufficient hCO models. For each gene, we first established mutant lines constitutively expressing the validated guide RNA by flow cytometry prior to creating hCO (Methods). We then applied CRISPR-A at the stages of developmental timepoints overlapping with robust expression of *CHD8* or SCN2A (Figure 1D, 2C). Additionally, as a complementary approach, we established 2D excitatory neuron cultures derived by *NGN2* overexpression [Lu et al., 2019; Zhang et al., 2013] and treated the cells with CRISPR-A at the neural progenitor stage (Figure 4A).

**Figure 4.**
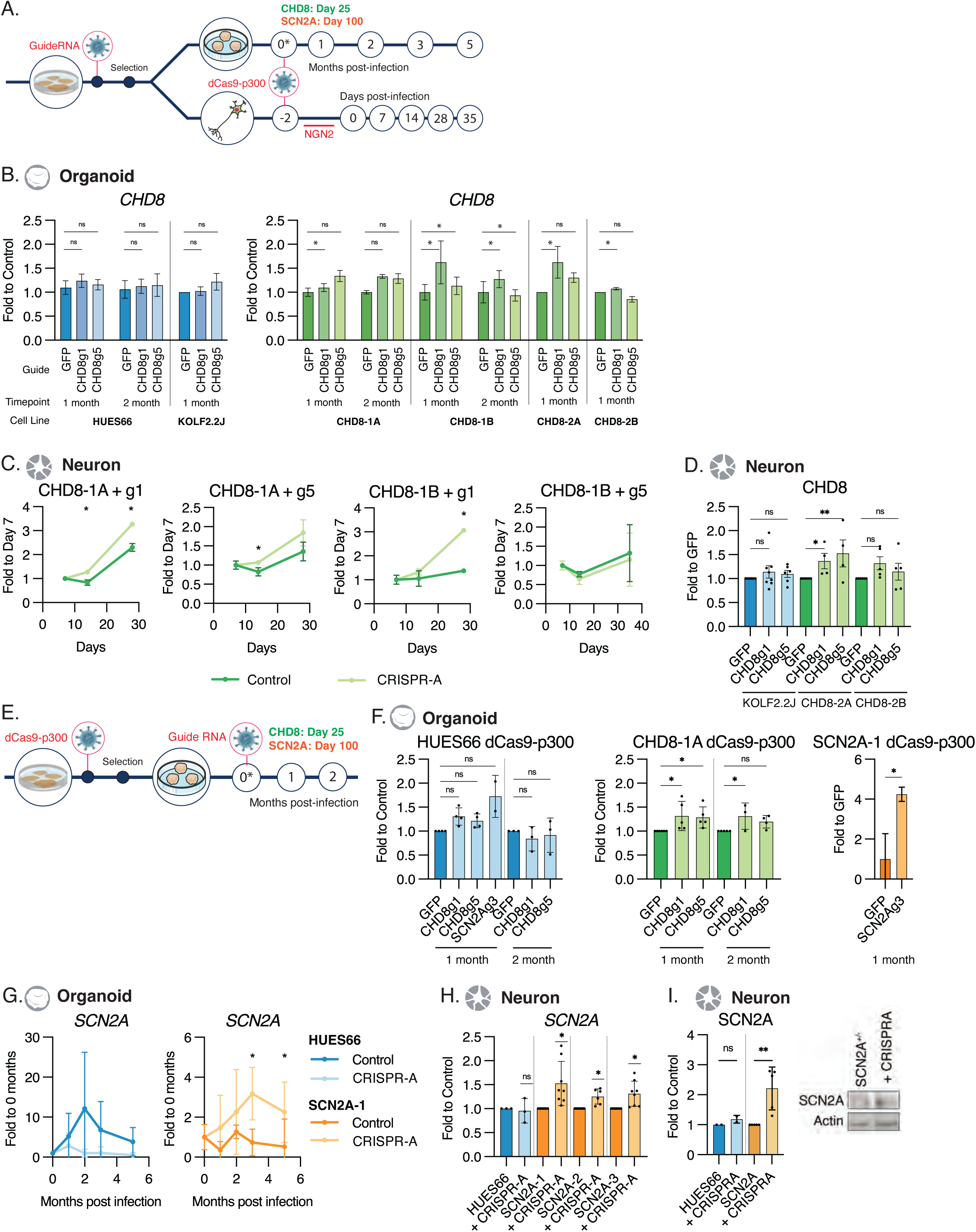
CRISPR-A treatment of mutant cortical organoids and neurons leads to increased expression of CHD8 and SCN2A. (A) Workflow for infection of hCO or iNGN2 neurons. Guide RNA are first transduced into cell lines and sorted by GFP. dCas9-p300 is transduced into the cells during differentiation; for iNGN2 neurons, at day -2 of the protocol and day 25 for CHD8 hCO or day 100 for SCN2A hCO. (B) CHD8 expression in hCO treated with CRISPR-A shows no increases in wildtype. Modest but significant increases are observed in the all CHD8-mutant lines for at least one guide RNA and timepoint. N = 5-9, SEM shown, * p < 0.05. (C) CHD8 mRNA expression is increased in CHD8-mutant iNGN2 neurons transduced with CRISPR-A. Neurons were collected at day 7, 14, 28, and 35 post-initiation. (D) CHD8 mRNA expression is increased in CHD8-mutant KOLF2.2J iNGN2 neurons but not in the parental line. Neurons were collected 35 days post initiation. N = 5, SEM shown, * p < 0.05. (E) Modified workflow for infection of hCO or iNGN2 neurons in which dCas9-p300 is first transduced into cell lines and selected for by puromycin. Guide RNA were transduced into the cells during differentiation at the same timepoints in (A). (F) hCO established from constitutively expressing dCas9-p300 lines show significant increases in CHD8 or SCN2A expression in mutant lines but not wildtype. N=3-6, SEM shown, * p < 0.05. All significance values calculated using Student’s two-tailed T-test. (G) SCN2A expression is rescued in mutant hCO but not wildtype. hCO were infected at 100 days old and collected 1-5 months post-infection. Expression shown as fold to 1-month post-infection control. (H) Expression of *SCN2A* is rescued by CRISPR-A in SCN2A-mutant neurons but not wildtype. Expression in wildtype (HUES66) is unchanged by CRISPR-A, whereas expression is significantly increased in the mutant lines. N = 7, SD shown, * p < 0.05. (I) Expression of SCN2A protein is significantly increased by CRISPR-A in SCN2A-1 neurons but not in wildtype neurons. N = 5, SD shown, * p < 0.05.

For *CHD8*, we infected mutant and wildtype hCO at 25 days old (Figure 4A) and collected samples monthly for three months for RNA. In the wildtype hCO, we observed no increase in *CHD8* expression (Figure 4B), consistent with observations from other groups using CRISPR-A on haploinsufficient genes [Colasante et al., 2019]. In CHD8-mutant hCO, we observed a general increase in CHD8 expression by both guide RNAs, though the level of activation varied by line and only CHD8-g1 consistently reached statistical significance (Figure 4B, Supplemental Figure 7A-E). In excitatory neuron cultures, both guides generated robust increases in *CHD8* expression in both *CHD8*^+/-^ lines during differentiation, with the most significant increases at 28-35 days old (Figure 4C). We found that this difference in effect of CRISPR-A was largely due to a difference in dCas9-p300 expression (Supplemental Figure 8A) between 2D excitatory neuron cultures and 3D hCO cultures. We observed more robust dCas9-p300 expression in excitatory neurons than in the hCO, while GFP levels, as a proxy for guide RNA expression, remained unchanged (Supplemental Figure 8B). By immunofluorescence, we observed that approximately 25% of cells in treated hCO expressed dCas9-p300 (Supplemental Figure 8C).

These results indicate that *CHD8* expression can be effectively increased by enhancer-targeted CRISPR-A, but are highly dependent on Cas9 levels, as has been shown [Maddineni et al., 2024; Uenaka et al., 2025]. To improve expression of dCas9-p300, we reversed the order of vector transduction in hCO; first establishing *CHD8*-mutant lines constitutively expressing dCas9-p300, then introducing guide RNA in 25-day-old hCO (Figure 4D). This approach led to significant increases in *CHD8* expression in *CHD8*-mutant hCO for both CHD8-targeting guides (Figure 4E, Supplemental Figure 7F & G).

As *SCN2A* expression peaks later in development, we infected wildtype and mutant cortical organoids at 100 days and collected samples monthly for up to five months post-infection. We observed no significant increase in *SCN2A* expression in wildtype hCO, matching the observation of the same organoids treated with *CHD8* gRNA. However, we found a significant increase in *SCN2A* expression in *SCN2A*^+/-^ hCO across all timepoints measured, though the degree of increase decreased over time (Figure 4F, Supplemental Figure 9A). We also established iNGN2 excitatory neuron cultures from the SCN2A-mutant lines and observed the same result; no significant change in *SCN2A* levels in wildtype neurons, but a significant increase in *SCN2A* mRNA and protein levels in the mutant lines treated with CRISPR-A (Figure 4G & H). We also established *SCN2A*-mutant organoids constitutively expressing dCas9-p300 and introduced the guide RNA at day 120; this approach also led to a significant increase in *SCN2A* expression (Figure 4H). These results demonstrate that *SCN2A* expression can be reliably increased by enhancer-targeted CRISPR-A.

We hypothesized that an enhancer-targeted activation approach would be preferential in some cases to promoter targeting, in that targeting enhancers is more likely to lead to physiological levels of gene expression and thus avoid concerns of ectopic spatial or temporal expression or overexpression of dosage sensitive genes. Indeed, introduction of gRNA for *CHD8* and *SCN2A* into hCO and neuron cultures led to a significant increase in target gene expression in *both* wildtype and mutant cells (Supplemental Figure 6B, 6C). These results suggest that while promoter and enhancer-targeted CRISPR-A both reliably increase target gene expression, for dosage sensitive genes, enhancer-targeting may be more physiological.

Additionally, while we chose to introduce CRISPR-A at timepoints related to the expected peaks in expression for *CHD8* and *SCN2A*, we also considered whether the addition of CRISPR-A at earlier or later timepoints would also yield increases in gene expression (Supplemental Figure 10). Using the HUES66-derived lines, we infected wildtype and mutant hCO at multiple timepoints (CHD8: 50, 100, 300 days old, SCN2A: 50 days old). No timepoints showed significant increases in gene expression in wildtype hCO. For CHD8, infection of 50- and 100-day old hCO lead to a modest increase in CHD8 expression that did not reach statistical significance, while the 300-day old hCO showed no increase in expression. For SCN2A, early introduction of CRISPR-A at day 50 did not lead to an increase in SCN2A expression (Supplemental Figure 10) in contrast to the day 100 infection (Figure 4G). These results support that the enhancers that we identified are sensitive to developmental time, relatively matching the trajectories of endogenous gene expression.

### CRISPR-A treatment rescues mutant phenotypes in *CHD8* and *SCN2A* cortical organoids and neurons

To further study the rescue of over-proliferation of progenitor cells caused by CHD8 haploinsufficiency, we quantified the expression of EOMES and DCX protein in two-dimensional (2-D) excitatory neuron cultures (Figure 5A). Compared to wildtype neurons, the CHD8-mutant neurons had significantly higher numbers of EOMES+ cells and significantly lower numbers of DCX+ cells, suggesting a larger population of intermediate progenitor cells. In contrast, mutant neurons infected with dCas9-p300 showed a significant decrease in EOMES expression and significant increase in DCX expression to levels near wildtype across both guide RNAs and both mutant lines (Figure 5B). Additionally, significantly more DCX+ cells were found to be present along cell processes (Figure 5C), suggesting an increase in neuron maturity with CRISPR-A. We quantified mRNA expression of *EOMES*, *DCX*, *TUBB3*, and *MAP2*, as markers for intermediate progenitors, immature neurons, and mature neurons, respectively, and observed a decrease in *EOMES* expression and increase in neuronal marker expression in treated excitatory neuron cultures (Figure 5D). Collectively, these results show that the overabundant progenitor phenotype caused by CHD8 haploinsufficiency is rescued by our enhancer-targeted CRISPR-A.

**Figure 5.**
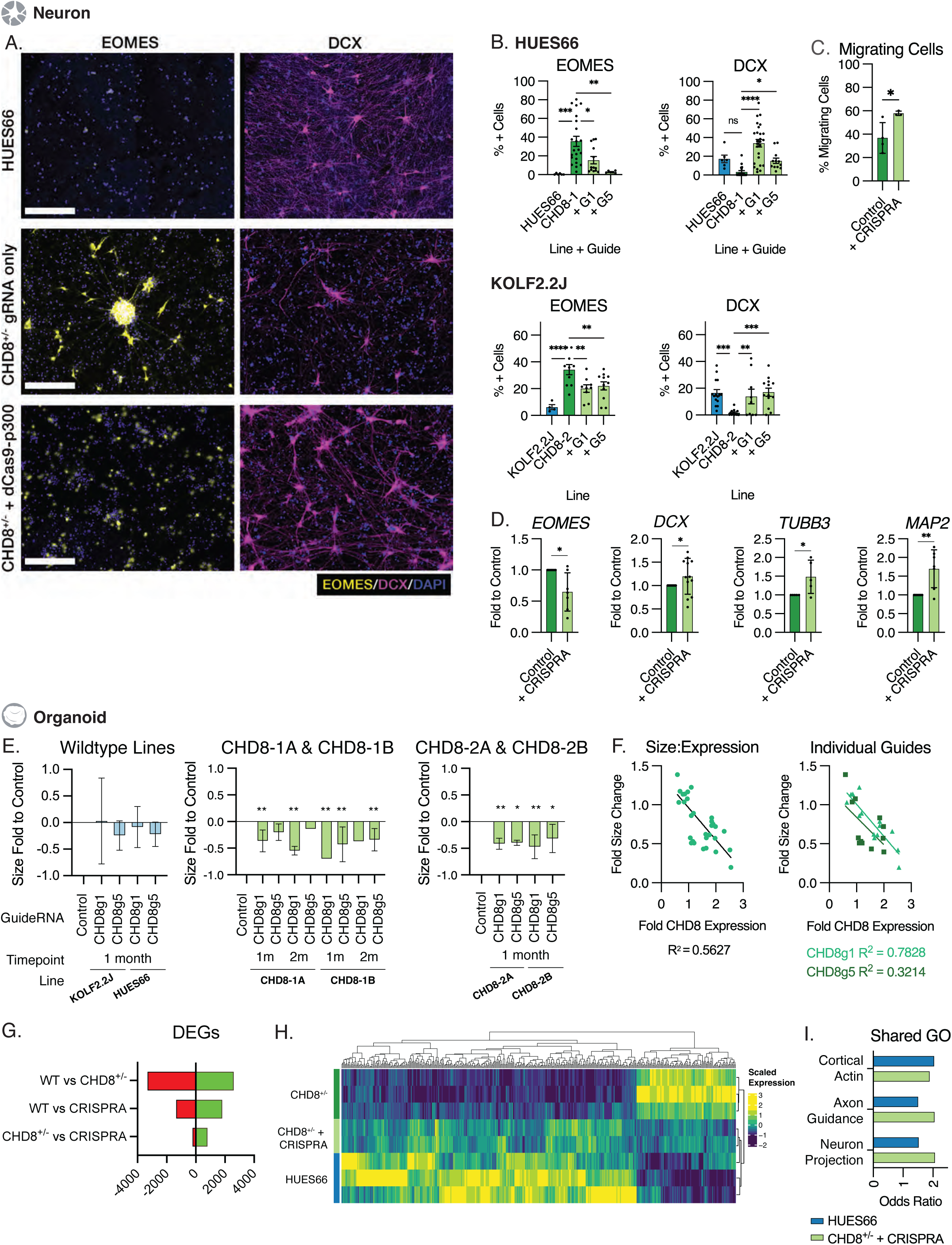
Aberrant phenotypes are reversed with CHD8 CRISPR-A treatment. (A) Expression of EOMES and DCX in CHD8-mutant iNGN2 neurons with and without dCas9-p300. (B) Quantification of immunofluorescence images show EOMES expression is significantly decreased by CRISPR-A treatment while DCX is significantly increased, suggesting increased maturity of the neurons. N = 9, SEM shown, * p < 0.05 (C) Increase in DCX+ cells on cell processes shows a significant increase in migrating neurons in CRISPR-A treated iNGN2 cultures. N = 5, SD shown, * p < 0.05. (D) Expression of cell type markers *EOMES*, *DCX*, *TUBB3*, and *MAP2* in iNGN2 neurons shows an increase in neuron maturity with CRISPR-A treatment. N = 5, SD shown, * p < 0.05. (E) hCO size is significantly reduced by CRISPR-A treatment. Size difference is represented as fold change relative to average control hCO size minus one. No significant size changes are observed in wildtype hCO. N = 50, SEM shown, * p < 0.05. (F) Correlation of CHD8 expression to changes in hCO size shows a positive correlation between increasing CHD8 expression and decreasing hCO size, stronger in CHD8g5 compared to CHD8g1. (G) Significant (padj < 0.05) differentially expressed genes (DEGs) from bulk RNAseq in hCO 1-month post-infection with CRISPR-A. Compared to CHD8^+/-^ hCO, HUES66 hCO show a high number of both upregulated and downregulated DEGs. Treatment of CHD8^+/-^ with CRISPR-A decreased the number of DEGs compared to HUES66. CHD8^+/-^ hCO treated with CRISPR-A show a larger proportion of upregulated than downregulated DEGs. (H) Heatmap of DEG expression shows CRISPR-A treatment induces a change in gene expression pattern towards wildtype organoids (HUES66). (I) Shared Gene Ontology terms related to neuron development (Cortical Actin GO:0030864, Axon Guidance GO:0007411, Neuron Projection GO:0097485) are significantly upregulated in both wildtype and CRISPR-A treated mutant hCO relative to control-treated hCO. All significance values calculated using Student’s two-tailed T-test.

As we observed in our initial characterization, *CHD8*^+/-^ hCO were significantly larger than wildtype at matched timepoints, a phenotype caused by an increase in progenitor cell proliferation, as previously described [Paulsen et al., 2022; Villa et al., 2020]. We therefore measured hCO size monthly after infection with dCas9-p300 to determine if an increase in CHD8 expression decreased hCO size. For CHD8-1A & B, both guides led to decreased organoid size compared to the vehicle-treated control, though only CHD8g1 lead to a statistically significant size reduction in CHD8-1A. In the KOLF2.2J-derived CHD8-2A & B hCO, both guides lead to statistically significant reductions in size after 1-month post-infection (Figure 5E). By comparison, no significant size changes were observed in wildtype HUES66 or KOLF 2.2J hCO treated with either CHD8-targeting guide. Given the variable increases to CHD8 gene expression that we observed with different guides and lines (Figure 4B & E), we tested to see if changes in hCO size change correlated with changes in levels of CHD8 expression. We observed a linear correlation, where increasing *CHD8* expression correlates with decreasing hCO size (Figure 5F); the correlation was higher for CHD8g5 than CHD8g1. Additionally, we found that promoter-targeted CRISPR-A significantly reduced hCO size in both wildtype and mutant lines (Supplemental Figure 6D, 6E); this again highlights a potential concern for dosage-sensitive genes.

To further characterize the effect of CRISPR-A on CHD8-mutant hCO, we performed bulk RNA sequencing on 50-day old hCO from wildtype, CHD8-mutant, and CHD8-mutant infected with CRISPR-A machinery. Comparing wildtype to mutant, we identified 5,933 significant differentially expressed genes (Methods; Figure 5G; p < 0.05, Wald test); 2,608 upregulated and 3,325 down-regulated in CHD8^+/-^ hCO. This pattern was partially reversed in CRISPR-A-treated hCO and unbiased clustering grouped the CRISPR-A-treated samples with wildtype and not the mutant samples, indicating that CRISPR-A infection led to a shift in expression towards wildtype expression (Figure 5H). Furthermore, Gene Ontology (GO) analysis of significantly upregulated genes in both HUES66 and CHD8^+/-^ + CRISPR-A hCO showed significant enrichment for GO terms associated with neuron maturation; including neuron projection, axon guidance, and cortical actin cytoskeleton, further evidence that the enhancer-targeted CRISPR-A rescues CHD8^+/-^ mutant phenotypes associated with neurogenesis and maturation (Figure 5I).

To determine the efficacy of using CRISPR-A to increase *SCN2A* expression, we focused on phenotypes related to neuron maturation and function that have been previously reported [Brown et al., 2024; Que et al., 2021]. Infecting *SCN2A*^+/-^ hCO with CRISPR-A lead to reversal of changes in both *EOMES* and *SATB2* expression that was sustained over the timepoints measured, though sample-to-sample variation between infected organoids was high, suggesting an inconsistent level of phenotype rescue (Figure 6A). We therefore measured expression of cell type markers in 2D differentiated excitatory neuron cultures and found a more consistent, significant increase in both *EOMES* and *SATB2* mRNA levels in neurons infected with CRISPR-A (Figure 6B). Additionally, we observed an increase in TUJ1 and MAP2 protein levels by immunoblot in neurons infected with CRISPR-A (Figure 6C, Supplemental Figure 9B) and a significant increase in SCN2A, DCX, and MAP2 protein expression by immunofluorescence in patient-derived neurons (Figure 6D). These results support the conclusion that CRISPR-A treatment leads to a more mature phenotype in excitatory neurons.

**Figure 6.**
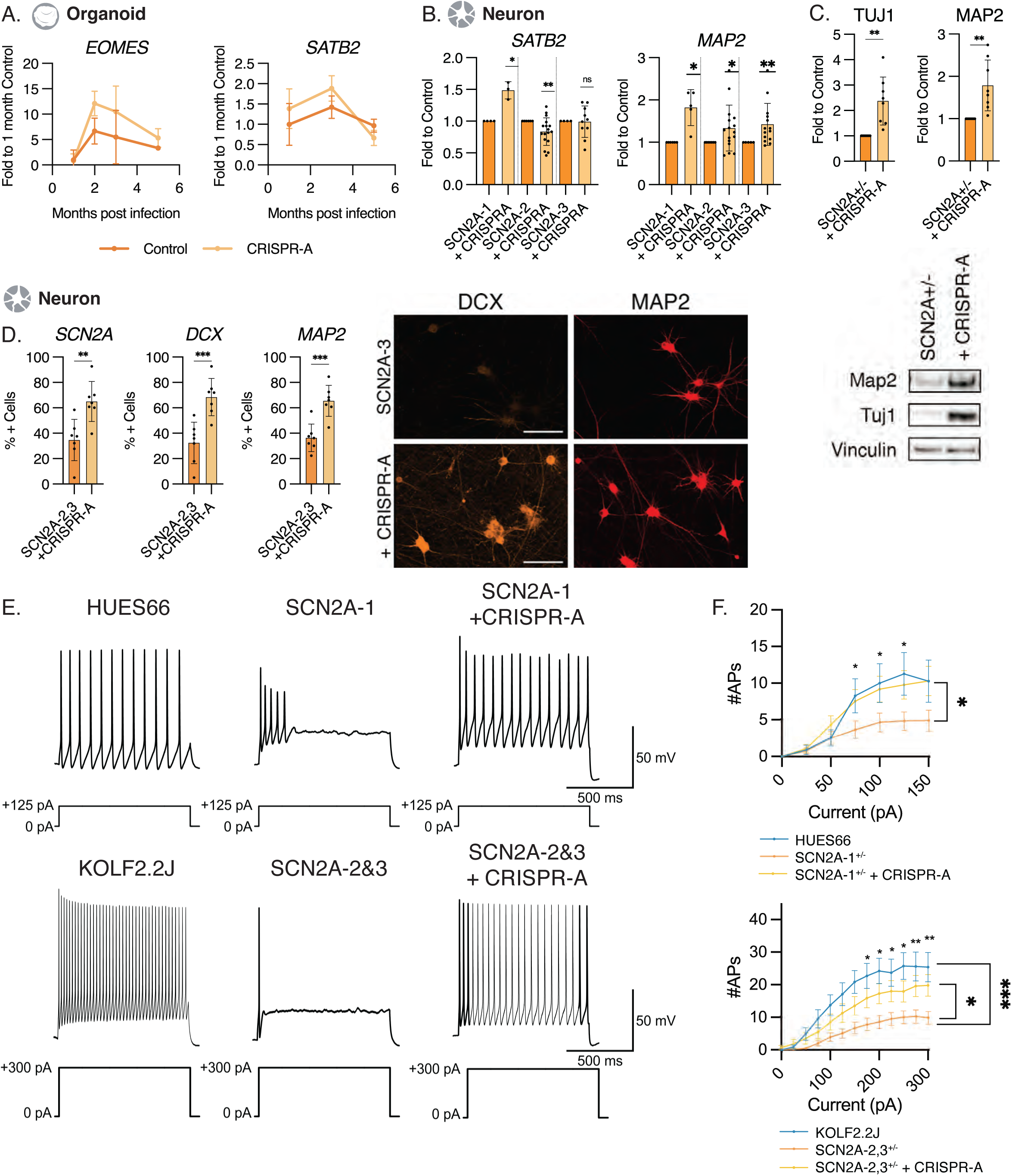
Aberrant phenotypes are reversed with SCN2A CRISPR 96-A treatment. (A) Expression of *EOMES* and *SATB2*, as markers for intermediate progenitor cells and upper layer neuron cells, respectively, are increased with CRISPR-A treatment of SCN2A-mutant hCO. N = 3-8, SEM shown. (B) Expression of *EOMES* and *SATB2* are increased with CRISPR-A treatment of SCN2A-mutant neurons. N = 5, SD shown, * p < 0.05. (C) Expression of TUJ1 and MAP2 protein are significantly increased by CRISPR-A treatment of SCN2A-mutant neurons. N = 8, SD shown, * p < 0.05, sample blot shown. Full blot images in Supplement. (D) Sample excitability traces from wildtype, SCN2A^+/-^, and SCN2A^+/-^ + CRISPR-A cells show partial rescue of sustained neuronal firing activity in response to a depolarizing current pulse (+125 and +300 pA, 1 sec). Cells were held at a membrane potential of -60 mV prior to current injection. (E) Graph comparing the number of action potentials (APs) fired in response to increasing current injections (1 sec). CRISPR-A treatment of mutant neurons significantly rescued the number of action potentials fired per injected current (N=11, N=14, and N=17 for HUES66, SCN2A^+/-^, and SCN2A^+/-^ +CRISPR-A, respectively). Plots depict mean number of action potentials ±SEM. Two-way repeated ANOVA revealed a significant difference between SCN2A^+/-^ and SCN2A^+/-^ +CRISPR-A distributions, indicated by * p < 0.05. Bonferroni *posthoc* analysis further showed significant differences at each level of current injection, where * p < 0.05, ** p < 0.01 and *** p < 0.001. All significance values calculated using Student’s two-tailed T-test or two-way repeated measures ANOVA followed by Holm-Sidak post hoc tests.

We next performed whole-cell patch-clamp recording on *SCN2A*^+/-^ neurons with or without CRISPR-A to determine whether there was physiological rescue. We were able to completely restore excitability to wildtype levels in HUES66-derived *SCN2A*^+/-^ + CRISPR-A neurons (Figure 6E&F). The addition of CRISPR-A significantly increased the number of action potentials fired in response to increasing current injections, reaching levels comparable to wildtype (Figure 6F). This occurred in the absence of detectable differences in rheobase measurements, AP threshold and RMP, indicating a nearly complete restoration of neuron function and excitability (Supplemental Figure 9). Similarly, in the patient-derived SCN2A-2 and SCN2A-3 iPSC neurons, CRISPR-A significantly increased the number of action potentials fired in response to increasing depolarizing current pulses compared to control (Figure 6F). The number evoked of action potentials fired in CRISPR-A-treated patient-derived neurons approached wildtype levels and did not differ significantly from neurotypical KOLF2.2J controls across most current pulses, with differences emerging at the highest current injections that elicit sustained firing. Consistent with the recovery of AP firing behavior, CRISPR-A normalized intrinsic excitability parameters in patient-derived iPSC mutant neurons, including rheobase and action potential threshold, whereas RMP remained similarly shifted in both mutant and CRISPR-A–treated neurons relative to wildtype controls (Supplementary Figure 9). Notably, patient-derived iPSC neurons required higher current injections to achieve sustained firing compared to HUES66-derived neurons across conditions, indicating a line-dependent difference in baseline intrinsic excitability, rather than incomplete functional rescue.

We hypothesized that this shift could be the result of compensatory gene expression of other Na^+^ channel subunits that are expressed at higher levels in patient-derived SCN2A-mutant lines (SCN2A-2,3) than the CRISPR-edited HUES66 (SCN2A-1) line. This genetic compensation has been has been observed in other disease and model contexts previously [El-Brolosy et al., 2019; Veitia et al., 2013], and there has been some evidence that suggests Na^+^ channel subunits behave similarly [Vega et al., 2008]. We therefore compared the expression of *SCN1A*, *SCN3A*, *SCN5A*, and *SCN8A* across all wildtype and SCN2A-mutant lines (Supplemental Figure 9). Expression of all subunits was lowest in the HUES66-derived SCN2A-1 line, while the patient-derived iPSC lines SCN2A-2 and SCN2A-3 showed subunit expression comparable to the wildtype KOLF2.2J. These findings suggest that elevated expression of alternative Na^+^ channel subunits in patient-derived iPSC neurons reshapes excitability parameters, including rheobase and action potential threshold, thereby shifting the input-output relationship and the current requirements for sustained firing without preventing robust CRISPR-A-mediated rescue of action potential firing. Taken together, we demonstrate that the enhancer-targeted gRNA for *CHD8* and *SCN2A* rescued mutant phenotypes in hCO and induced neurons in 2D cultures.

## Discussion

There are few currently available therapies other than symptomatic treatments for ASD, whose efficacy are limited by clinical and genetic heterogeneity. Here we leveraged the growth of nucleic acid therapeutics, which is a promising frontier, especially for the treatment of rare brain diseases [Chen, and Geschwind, 2022; Ricci, and Colasante, 2021; Sahin, and Sur, 2015; Weuring et al., 2021; Whiteley et al., 2022]. We hypothesized that by using CRISPR-A to target a haploinsufficient gene, one could increase the expression of the wildtype allele to restore gene expression to normal levels. CRISPR-A is more generalizable than other precision medicine technologies, as it may be more broadly applied to any individual with a haploinsufficient mutation in the given gene, as demonstrated in other disease models. This includes *Sim1* in obesity [Matharu et al., 2018], *Scn1a* in Dravet Syndrome [Colasante et al., 2019], and others [Colasante et al., 2020; Jaudon et al., 2022; Qian et al., 2023; Tamura et al., 2022]. But it is not yet widely appreciated as a therapeutic approach. It is worth noting that haploinsufficient genes have been shown in animal models to be embryonic lethal when knocked out, and their overexpression is associated with decreased fitness [Collins et al., 2020; Huang et al., 2010; Kawamura et al., 2025; Morrill, and Amon, 2019], emphasizing the importance of moderating any induced increase in expression. With this in mind, we targeted enhancer regions. For both *CHD8* and *SCN2A*, we were able to moderately increase the expression of each gene in mutant hCO using a single enhancer-targeted gRNA paired with dCas9-p300. This significant increase in expression resulted in partial or complete rescue of mutant phenotypes; increase of *CHD8* led to near normalization of organoid size, decreased progenitors, and increased neurons, as well as partial normalization of changes in gene expression observed with the mutation. Increase of *SCN2A* expression increased the expression of upper layer neuron markers and rescued neuronal firing output in excitatory neuron cultures, with line-dependent normalization of intrinsic excitability parameters.

We replicate previous findings by showing the phenotype of over-proliferation in *CHD8* haploinsufficiency and electrophysiological abnormalities in *SCN2A* haploinsufficiency [Deneault et al., 2018; Paulsen et al., 2022; Spratt et al., 2019], two monogenic causes of ASD that have very high penetrance for ASD (OR >100; [Rolland et al., 2023]). We observed a significant increase in size in *CHD8*^+/-^ hCO compared to wildtype at matched timepoints, a phenotype that appears analogous to macrocephaly observed in patients [Bernier et al., 2014; Villa et al., 2020]. The increase in size in *CHD8*^+/-^ hCO resulted from an increase in progenitor cell proliferation, consistent with previously published studies using different brain organoid [Paulsen et al., 2022; Villa et al., 2020; Wang et al., 2017] and animal [Dong et al., 2022; Gompers et al., 2017; Ji et al., 2025; Nord et al., 2025] models. We also found *CHD8*^+/-^ neurons to be less mature than wildtype, consistent with the role that *CHD8* and its downstream targets play in neurogenesis [Villa et al., 2020]. In *CHD8*-deficient hCO and neurons, we were able to increase *CHD8* expression with CRISPR-A, resulting in a decrease in organoid size and increase in neuronal differentiation. These results point to a rescue of neurogenesis, which has been highlighted in other studies [Dong et al., 2022; Gompers et al., 2017; Nord et al., 2025; Paulsen et al., 2022; Wang et al., 2017] as a key mutant phenotype.

The two HUES66-derived *CHD8*^+/-^ stem cell lines used in this study differ by only an additional four base pair deletion in CHD8-1B compared to CHD8-1A (Supplemental Figure 1, [Shi et al., 2023]). While the two lines both demonstrate decreased *CHD8* expression as stem cells, neurons, and organoids, we observed moderate differences in hCO composition (Supplemental Figure 4), as well as response to CRISPR-A (Figures 4, 5). However, none of these differences reached statistical significance and differential gene expression analysis from bulk RNAseq data comparing CHD8-1A and CHD8-1B organoids yielded no significant DGE. To validate the findings in these lines, we repeated a subset of key experiments using CHD8^+/-^ lines derived from the well-characterized KOLF2.2J iPSC line (CHD8-2A & CHD8-2B). Both KOLF2.2J-derived lines showed increased CHD8 expression in response to CRISPR-A as hCO or excitatory neuron cultures, increased neuron maturation, and decreased hCO size, indicating that our findings are not limited to the HUES66 background.

In *SCN2A*-deficient hCO, we observed a decrease in upper layer neurons compared to wildtype, specifically at later points in development, epochs at which *SCN2A* expression is expected to peak. These results are consistent with animal models that show impaired excitability in slices of prefrontal cortex [Eaton et al., 2021; Spratt et al., 2019; Yang et al., 2023]. Neurons from this mutant line showed impaired excitability and reduced network activity, mirroring other studies that sought to model ASD-associated variants of *SCN2A* [Ben-Shalom et al., 2017; Brown et al., 2024; Que et al., 2021; Tatsukawa et al., 2019]. In *SCN2A*-deficient hCO and iNGN2 neurons, we were able to rescue gene expression to near wildtype levels and subsequently observe a rescue of signaling activity in excitatory neurons. To further validate our findings, we established neuron cultures from two *SCN2A*^+/-^ patient-derived iPSC lines, which showed similar impaired excitability and rescue of SCN2A expression and neuron excitability with CRISPR-A. Similar results were also recently demonstrated using a *SCN2A* promoter-targeting CRISPR-A system, where the investigators paid careful attention to avoid over-expression by using a less powerful activating construct [Tamura et al., 2022].

Despite our concerns of overexpression in using promoter-targeted gRNA constructs, work by other groups have targeted promoter regions of their genes of interest with no apparent impact on cell fitness [Colasante et al., 2019; Colasante et al., 2020; Matharu et al., 2018; Tamura et al., 2022; Tian et al., 2021]. This may be due to several factors including, endogenous regulatory mechanisms to limit gene overexpression in an organ-specific manner, the culture system in which the assay is conducted [Ho et al., 2017; Tian et al., 2021; Wu et al., 2023], differing tolerances for gene overexpression based upon normal gene function, or limitations on the effect of the dCas9-bound activator complex. The question of organ specificity was briefly explored in *Sim1* [Matharu et al., 2018], in which promoter and enhancer targeting guides were tested in multiple organ systems, with varying degrees of gene activation between organs. We note that in our *CHD8* experiments, we utilized two guide RNA constructs which exhibited differing levels of *CHD8* activation. As these guides targeted different enhancers, we hypothesize that differences between these enhancers accounts for differences in activation levels, which is in line with recent screening data that suggest restrictions on enhancer activity dependent upon cell type [Chardon et al., 2024]. Additionally, while the focus of our study was to address the feasibility of targeting enhancer regions for gene activation, we designed promoter-targeting guide RNA for both *CHD8* and *SCN2A*, finding that these constructs lead to robust increases in target gene expression in wildtype and mutant hCO and neuron cultures (Supplemental Figure 6). We observed the same phenotypic changes in both wildtype and mutant cultures; a decrease in hCO size when treated with CHD8 CRISPR-A and an increase in MAP2 expression from both CHD8 and SCN2A activation. Given that our enhancer-targeted constructs did not lead to any activation in wildtype lines, these observations suggest that enhancer-targeted CRISPR-A may allow for the use of more powerful activation constructs without the risk of overexpression. Moreover, we show a degree of temporal specificity with enhancer targeting that has not previously been demonstrated with promoter targeting constructs [Colasante et al., 2019; Tamura et al., 2022; Matharu et al., 2018].

We view these results as an important proof of principle and recognize limitations that future work should address. First, the combined expression of both guide RNA and dCas9-p300 in hCO was limited by the ability of the dCas9-p300 lentivirus to infect only a subset of cells in the hCO (Supplemental Figure 8). To address this, we repeated key experiments using hCO constitutively expressing dCas9-p300 and introduced the guide RNA at the developmentally relevant timepoint for each gene. This approach resulted in a more reliable increase in target gene expression, but one that was not substantially greater than the previous transduction approach. This indicates that further optimization of viral infection, constructs, and use of alternative Cas enzymes [Knott, and Doudna, 2018; Mahata et al., 2023; Maddineni et al., 2024; Tycko et al., 2025; Li et al., 2020] would increase the clinical utility of this approach.

Second, we noted the effect of CRISPR-A decreased over time, with a decrease in expression of *CHD8* or *SCN2A* between two- and five-months post-infection. From the perspective of paralleling endogenous expression during development this may be optimal, although it suggests that rescuing adult expression may require targeting different enhancers or the gene promoters. Third, there is evidence that multiple gRNA may exhibit synergistic effects when used to target the same gene of interest [Hilton et al., 2015], though the same group noted that dCas9-p300 appeared to have a moderately additive, rather than synergistic level of increase in activation. Given the limited potential benefit of combining gRNA, we kept a narrow focus on testing individual gRNAs alone, though future optimization may include combining guides to increase gene expression. We also performed a post hoc analysis of predicted gRNA efficiency based on insights from CRISPR-editing [Konstantakos et al., 2022]. However, based upon the guides screened for this project, as well as additional hcASD target genes that we have studied, we do not find the guidelines for editing efficiency to translate to success in gene activation. Rather, cell type specific characteristics of a particular enhancer are more likely to predict gRNA efficacy (Supplemental Figure 11 and Supplemental Table 1). Nonetheless, the fact that mutant phenotypes were still partially, but significantly, rescued points to the potential for increased phenotypic rescue with increased efficiency.

Caveats to the successful clinical application of CRISPR-A include the potential for off-target effects, as is the case for most therapeutic approaches. To control for this in our study, we measured the expression of neighboring and closely associated genes to *CHD8* and *SCN2A* and found no increases in expression levels. Second, as both *CHD8* and *SCN2A* vary in their expression across developmental time, there is an underlying concern of increasing expression at the relevant period in development. To address this, we focused on enhancers predicted to be specifically active during fetal brain development. Additionally, we applied the CRISPR-A system to mutant organoids at the period of development where *CHD8* or *SCN2A* expression peaked in wildtype organoids. Endogenous gene expression regulation may render the timing of CRISPR-A treatment irrelevant, though an open question remains regarding the effect of introducing CRISPR-A after the expected peak in expression. It is likely that this would vary on a gene-by-gene basis; as *CHD8* is a key regulator of many downstream processes, including many other hcASD genes, it is likely that later treatment with CRISPR-A may have limited benefit. By contrast, *SCN2A* potentially could be targeted later, since neurons express the sodium channel subunit, and *SCN2A* expression is successively replaced by *SCN8A* as development progresses [Brunklaus, and Lal, 2020], which may also contribute to a milder phenotype. Lastly, for genes such as *CHD8*, where extra-CNS manifestations are observed, different enhancer guides may be necessary to target all involved organ systems.

In summary, our work demonstrates the practical application of iPSC-derived cells in modeling neurodevelopmental disorders and testing clinical interventions. We provide a workflow that can be adapted to target other hcASD genes and a proof-of-concept for the use of gene activation techniques as a therapeutic intervention for ASD.

## Materials and Methods

### Guide RNA Design

Putative enhancer regions were identified using data from previously published studies from our lab [de la Torre-Ubieta et al., 2018; Won et al., 2019; Walker et al., 2019]. We used the ECR browser (Ovcharenko et al., 2004; RRID:SCR_001052) to check identified sequences for conservation in the Mus Musculus genome, for the potential use of guides in animal studies. Using the UCSC Genome Browser (Frankish et al., 2018; RRID:SCR_005780), we focused on sequences within identified regions that corresponded with a DNAse I binding site. The resulting optimized sequences were run through CRISPOR (Concordet & Haeussler, 2018; RRID:SCR_015935) to design putative guide RNA. One to three top-scoring guide RNA sequences per enhancer were chosen for synthesis.

### Golden Gate Assembly

Per guide RNA, “CACCG” was appended to the 5’ end of the sense strand, “AAAC” was appended to the 5’ end of the antisense strand. Single strand DNA oligonucleotides were ligated using T4 Polynucleuotide Kinase (New England Biolabs M0201) using manufacturer instructions. Ligated oligonucleotides were then assembled with pLV-U6-sgRNA using BsmBI (New England Biolabs R0739S) and T7 ligase (ThermoFisher Scientific EP0081). pLV-U6-gRNA-UbC-eGFP-P2A-Bsr was a gift from Charles Gersbach (Addgene plasmid # 83925; http://n2t.net/addgene:83925; RRID:Addgene_83925). Plasmids were transformed into DH5a competent cells (New England Biolabs) on LB agar plates with ampicillin. After overnight culture, single colonies were picked and cultured overnight in Terrific Broth with Ampicillin. Plasmids were purified using a Zymo miniprep kit (Zymo Research D4210) and sequenced to verify correct insertion.

### CRISPR-edited line generation

HUES66-derived CHD8 and SCN2A mutant lines were established as previously described [Lu et al., 2019; Shi et al., 2023]. To generate clonal knockout lines for CHD8 in the KOLF2.2J iPSC line, a high-throughput multi-gRNA ribonucleoprotein (RNP) approach was employed. A pool of three gRNAs targeting an early exon of the gene (UGAAUCGAAACGCAUCACCC, GGACAUCGGCAUGUUGUGCU, CAGCUGGAUAGUUACUACCU) was designed and purchased from Edit Co’s GKO Revolution Kit.

For RNP assembly, 6 µl of 33 µM gRNA pool was mixed with 2 µl of 20 µM Alt-R™ S.p. Cas9-RFP V3 (IDT, 10008163) and incubated for 10 minutes at room temperature. The resulting RNPs were then added to 500,000 cells suspended in 20 µl of Lonza’s X solution (Lonza P3 Primary Cell 4D X Kit, V4XP-3032). Nucleofection was performed using the Lonza Nucleofector 4D, after which cells were resuspended in mTeSR™ Plus medium (Stem Cell Technologies, 100-0276) supplemented with a 1:10 dilution of CloneR™2 (Stem Cell Technologies, 100-0691).

Approximately 90% of the culture exhibited RFP fluorescence 24 hours post-nucleofection. Cells were then resuspended using Accutase (Invitrogen 00-4555-56) and plated at a density of 1 cell per well in 96-well plates via limiting dilution.

Colonies were imaged daily for 10 days. To ensure accurate selection of viable colonies, defined as those likely to be clonal, originating from a single cell, and lacking overt signs of differentiation, we applied a deep neural network–based segmentation model, Omnipose [Cutler et al., 2022], which we trained specifically for this task. After 10 days of culture, viable colonies were expanded and subjected to Sanger sequencing to confirm successful editing, determine zygosity, and verify maintenance of clonality. (Forward Primer: TTGTCCTGGGCTTTGTCCTG, Reverse Primer: TGCATCAAGACAGATAGGATGATG, Sanger Sequencing Primer: CTTTGTCCTGGTCCCATTATCTGAG).

For quality control, pluripotency was assessed via immunofluorescent staining for DAPI, (Sigma-Aldrich D95420), OCT4, and TRA-1-60 (Antibodies in Table 3). Marker expression was quantified using Omnipose-based segmentation, followed by analysis in CellProfiler [Stirling et al., 2021], confirming that >99% of DAPI⁺ nuclei co-expressed OCT4. Finally, low-pass whole-genome sequencing (0.3× coverage) was performed to assess genomic stability.

### Cell Culture

HEK293T (RRID:CVCL_0063) cells were cultured in DMEM + 10% FBS. For guide RNA screening, 4*10^5 cells per well were seeded into a 12-well tissue culture plate. The next day, cells were transfected with 300ng/well dCas9-p300 plasmid and 200ng gRNA plasmid using Lipofectamine 2000 (ThermoFisher 11668500) and incubated for 48 hours. Cells were then collected with Trizol (ThermoFisher) and RNA extracted using a DirectZol RNA miniprep kit (Zymo R2052). cDNA was synthesized from 500ng RNA using a Multiscribe High Capacity cDNA Synthesis kit (ThermoFisher 4368813) and qPCR was performed using Luna qPCR Master Mix (New England Biolabs M3003X) against primers listed in table 2.

Human Neural Progenitor cells (hNPs) were isolated from discarded tissues as previously published [de la Torre-Ubieta et al., 2018] and cultured on laminin-coated TC plates. For guide RNA screening, 2*10^5 cells per well were plated in 6-well plates. After 24 hours, cells were transduced with lentivirus with 6 ug/ml Polybrene overnight. Media was replaced after overnight incubation and cells were incubated for 72 hours prior to collecting the cells for RNA isolation, cDNA synthesis, and qPCR as described with HEK293T cells.

Stem cells were thawed and plated into 1 well of a 6-well plate coated with vitronectin using E8 media (ThermoFisher A1517001) or on Matrigel (Corning 354277) with mTESR Plus media (StemCell Technologies 100-0276). Media was changed daily and cells passaged weekly. The dCas9-p300 line was a generous gift from Dr. Charles Gersbach.

### Lentiviral Production

HEK293T cells were plated in 10 cm tissue culture dishes at a density of 3.8*10^5 cells per dish. The next day, cells were transfected with 9ug psPAX2 (RRID:Addgene_12260), 900ng pCAG-VSVG (RRID:Addgene_64084), and 9ug plasmid of interest using OptiMEM (ThermoFisher 31985062) and Lipofectamine 3000 (ThermoFisher STEM00001). After 18 hours, cell media was replaced by DMEM + 10% FBS + BSA and harvested every 24 hours for 48 hours. Collected crude virus was concentrated with Lenti-X (Takara Bio 631231) following manufacturer instructions. Titer was determined using Lenti-X qRT-PCR Titration Kit (Takara Bio 631235). pLV-dCas9-p300-P2A-PuroR was a gift from Charles Gersbach (Addgene plasmid # 83889; http://n2t.net/addgene:83889; RRID:Addgene_83889). SP-dCas9-VPR was a gift from George Church (Addgene plasmid # 63798; http://n2t.net/addgene:63798; RRID:Addgene_63798).

### Cortical organoids

Human cortical organoids were produced as previously published [Pasca et al., 2015; Yoon et al., 2019]. Briefly, stem cells were seeded into an Aggrewell 800 tissue culture plate (Stem Cell Technologies 34815) at 3*10^6 cells per well in 1.5mL E8 media (Thermo). Every 24 hours for 3 days, media was replaced without disrupting the embryoid bodies in the microwells with E6 media containing 5uM dorsomorphin (PubChem 11524144), 10uM SB431542 (PubChem 4521392), and 2.5uM XAV939 (PubChem 329830802). On the fourth day, EBs were collected from each well with a wide bore pipette tip and a 40um strainer (Pluriselect 43-50040-51) to be plated in 10cm ultralow attachment tissue culture dishes (Corning 4615, Sbio MS-90900Z). The culture was undisturbed for the fifth day. After an additional 2 days of culture in E6 media with dorsomorphin, SB431542, and XAV939 with daily changes, the media is changed to Neurobasal A (ThermoFisher 10888022) supplemented with B27 without vitamin A (ThermoFisher 12587010), GlutaMAX (ThermoFisher 35050061), 20ng/ml EGF, and 20ng/ml FGF and replaced daily until day 17 post-initiation. From day 18-25, media is replaced every other day. From day 26-43, the media is replaced every other day with Neurobasal A with B27 without vitamin A, GlutaMAX, 20ng/ml BDNF, and 20ng/ml NT3. After day 43, media is replaced approximately every four days with Neurobasal A with B27 without vitamin A and GlutaMAX.

For organoid infection, three organoids per condition were plated in an ultralow attachment 24-well plate (Corning 3473). Lentivirus was infected at MOI 7 for 24 hours. Organoids were collected post-infection for RNA and imaging. For each application, a minimum of three organoids were combined to create one biological replicate for bulk RNA isolation. Organoids were collected and flash frozen in Trizol. RNA isolation was performed using an Arcteus PicoPure RNA Isolation Kit (ThermoFisher KIT0204). RNA integrity and concentration was determined using an Agilent Bioanalyzer RNA Pico chip (Agilent 5067-1513). cDNA synthesis was performed using a Multiscribe High-Capacity cDNA synthesis kit (ThermoFisher) or SuperScript IV (ThermoFisher 11766050).

### Bulk RNA Sequencing

RNA sequencing libraries were prepared using the Ovation RNA Ultralow input kit (Tecan) and sequenced using an Illumina NovaSeq S4 flow cell (PE 2 x 100). Raw reads were trimmed with cutadapt (Martin, 2011; RRID:SCR_011841) and aligned to hg19 using hisat2 (RRID:SCR_015530). Differential expression analysis was conducted using DESeq2 (Love et al., 2014; RRID:SCR_015687). Gene Ontology analysis was conducted using clusterProfiler (Wu et al., 2021; RRID:SCR_016884).

### Organoid size measurements

Tissue culture plates were photographed on a lightbox with a metric ruler to standardize measurements between images. Diameter measurements were taken in pixels using Adobe Photoshop (RRID:SCR_014199), and the volume calculated assuming an ellipsoidal shape.

### Neuron differentiation

Stem cells were differentiated to excitatory neurons using NGN2 overexpression as described previously [Zhang et al., 2013]. Briefly, stem cell lines were stably transduced to express TetO-Ngn2-Puro and Ubq-rtTA. Cells were plated onto Matrigel-coated 6-well plates at a density of 4*10^5 to 8*10^5 cells/well in pre-differentiation media comprised of Knockout DMEM:F12 (ThermoFisher 12660012) with 1x N2 Supplement (ThermoFisher 17502048), 1x non-essential amino acids (NEAA, ThermoFisher 11140050), 10ng/ml BDNF, 10ng/ml NT3, 1ug/ml laminin, and 2ug/ml doxycycline. Media was changed daily. After two days in pre-differentiation media, cells were collected and plated onto laminin-coated coverslips or tissue culture wells at a density of 1*10^5-8*10^5 cells/well in maturation media comprised of 50% Neurobasal A, 50% DMEM:F12, 0.5x N2 supplement, 0.5x B27 supplement, 1x NEAA, 0.5x GlutaMAX, 10ng/ml BDNF, 10ng/ml NT3, 1ug/ml laminin, and 2ug/ml doxycycline. This was considered to be day 0 of differentiation. At day 3, half of the media was replaced with fresh maturation media. At day 7, half of the media was replaced again, now without doxycycline. At day 14, half of the media was removed and twice that volume of maturation media was added, for a volume 1.5x the original plating volume. This was repeated at day 21, resulting in a 2x final volume. A third of the media was subsequently replaced weekly until neurons were used for downstream applications.

### Immunofluorescence

#### Organoid sectioning

Organoids were fixed with 4% PFA for 24 hours and rinsed with PBS. Samples were then transferred to a 30% sucrose solution and incubated overnight at 4°C. Using an embedding mold, organoids were enrobed in 1% agarose and cut into 40um sections on a vibratome (Leica).

#### Immunofluorescence

Organoid sections were blocked in PBS with 5% FBS, 1% BSA, 0.3% Triton-X, and 0.1% sodium azide for 20 minutes at room temperature. Primary antibody was diluted in blocking buffer and the section was incubated overnight at 4C. The next day, the section was washed three times with PBS + 0.1% Tween-20. Secondary antibody was diluted in blocking buffer and the section was incubated in the dark at room temperature on a rocker. After three washes with PBS-T, the section was mounted on a slide using Fluormount with DAPI (Southern Biotech 0100-20). Antibodies used are listed in Table 3. Full immunofluorescence images are in Supplemental Figure 14.

#### Imaging and analysis

Slides were imaged on a Zeiss LSM 900. Whole slide images were captured using an Akoya Phenoimager (Akoya Biosciences). Images were quantified using QuPath (Bankhead et al., 2017; RRID:SCR_018257). To quantify cells, we first used QuPath’s cell detection tool to identify cells based on DAPI staining. Next, single channel classifiers were trained on each fluorophore, using a simple threshold for signal intensity. Classifiers were then run together to identify cells staining positive for multiple markers. The same classifiers were used for all slides in the same experiment.

### Western blotting

Cells or organoids were lysed in RIPA buffer containing Halt protease inhibitor (Thermo 87786) for 30 minutes on ice. Whole cell lysates were centrifuged for 15 minutes at 12,000 g at 4C and the supernatants transferred to new microtubes. Protein concentration was quantified using a Pierce BCA assay (Thermo 23227) following manufacturer’s instructions. Lysates were normalized to 10-50ug protein per sample and diluted with RIPA buffer and 5X denaturing loading buffer prior to heating at 95C for 5 minutes. Samples were loaded into Mini-Protean 4-20% TGX precast gel (BioRad 4561094) or 3-8% NuPAGE precast gel (ThermoFisher EA03752BOX) and run at 100V at 4C for 3 hours. Gels were transferred wet onto nitrocellulose membranes overnight at 25V at 4C for 16 hours. Membranes were blocked with 5% BSA in PBS-Tween for 1 hour at room temperature. Primary antibodies (Table 3) were diluted in blocking buffer and incubated with the membrane overnight at 4C. Blots were subsequently washed three times with PBS-T and incubated in HRP-conjugated secondary antibody diluted in blocking buffer at room temperature for 2 hours. After three additional washes, blots were developed using SuperSignal West Dura ECL (ThermoFisher 34075) and imaged using an Azure Biosystems C600 imager. Images were annotated using Adobe Photoshop and quantified using ImageJ. Full blot images are collected in Supplemental Figure 12.

### Flow cytometry

Organoids were dissociated using a modified protocol from the Worthington Papain Dissociation System (Worthington NC9067191). Briefly, 5-8 organoids were transferred into 2.5ml of freshly-reconstituted papain in EBSS. The organoids were incubated at 37C on a shaker at 100rpm, initially for 30 minutes, pausing every 15 minutes to manually triturate the sample. After no large clumps remained, the dissociated sample was centrifuged at 300g for 5 minutes. The supernatant was discarded and albumin ovomucoid inhibitor solution added per manufacturer’s instructions. Cells were then fixed using 4% paraformaldehyde for 15 minutes at room temperature. Cells were then washed with FACS buffer and counted. Samples were divided into approximately 1*10^6 cells/sample for flow cytometry. If permeabilization was necessary, cells were incubated in 0.1% saponin in HBSS for 15 minutes at RT prior to antibody incubation. Cells were incubated in primary antibody for 1 hour at 4C in the dark (Table 3). If a secondary antibody was necessary, cells were washed twice following primary incubation and incubated in secondary antibody for 1 hour at 4C in the dark. Cells were subsequently washed twice and Sytox added to the final cell suspension.

EdU staining were conducted per manufacturer’s instructions. Organoids are stained with EdU for 2 hours, then incubated for 48 hours prior to dissociation. Click-IT chemistry is conducted per manufacturer’s protocol. Samples were processed on a BD LSR II flow cytometer and data was analyzed using FlowJo (Treestar). Gating strategy is shown in Supplemental Figure 13.

### Electrophysiology

Neurons were differentiated on laminin-coated coverslips as described above. Coverslips were removed from culture wells and placed in an electrophysiology chamber filled with standard artificial cerebral spinal fluid (ACSF) containing (in mM): 130 NaCl, 3 KCl, 1.25 NaH_2_PO4, 26 NaHCO_3_, 2 MgCl_2_, 2 CaCl_2_, and 10 glucose) oxygenated with 95% O_2_-5% CO_2_ (pH 7.2-7.4, osmolality 290-310 mOsm/L). All recordings were performed at room temperature using an upright microscope (Olympus BX51WI) equipped with differential interference contrast optics (IR-DIC) and fluorescence imaging. Whole-cell patch clamp recordings in voltage- and current-clamp modes were obtained from cells using a MultiClamp 700B amplifier (Molecular Devices) and the Clampex 10.7 acquisition software. The patch pipette (3–5 MΩ resistance) contained a K-gluconate-based solution comprised of (in mM): 112.5 K-gluconate, 4 NaCl, 17.5 KCl, 0.5 CaCl_2_, 1 MgCl_2_, 5 ATP (potassium salt), 1 NaGTP, 5 EGTA, 10 HEPES, pH 7.2 (270–280 mOsm/L). Within 5 min of breaking through the cell membrane, passive membrane properties (capacitance and input resistance) were recorded while holding the membrane potential at -70 mV. Spontaneous postsynaptic currents (SPCs) were recorded in gap-free voltage-clamp mode at -70 mV. Resting membrane potential (RMP) and excitability measurements were recorded in current-clamp mode. Rheobase was determined using brief depolarizing current pulses (5 ms) applied incrementally until action potential firing was elicited. Input-output relationships as a test for cellular excitability were assessed using 1 sec depolarizing current injections in 25 pA increments. All recordings were filtered at 1-2 kHz and digitized at 100 μs. Electrode access resistances were maintained at <30 MΩ, and recordings exceeding this threshold were excluded from analysis.

Whole-cell patch clamp recordings were obtained from individual neurons across independent differentiation batches. Recordings were performed across 3-8 independent differentiation batches, with 1-6 cells recorded per batch. Each recorded cell was treated as an independent biological replicate for statistical analysis. Sample sizes for each condition are reported in the corresponding figure legends.

## Supporting information

All supplemental figures

## Data and Accessibility

Bulk RNA sequencing data is deposited in GEO (Accession #).

## Acknowledgements

The authors would like to thank the members of the Geschwind lab for their valuable feedback, Dr. Charlie Gersbach and Dr. Lexi Bounds for providing the dCas9-p300 construct and constitutively expressing stem cell lines. The SCN2A+/- patient-derived lines were selected from the Simons Foundation Autism Research Initiative (SFARI) iPS repository. We additionally would like to thank the UCLA Neuroscience Genomics Core and UCLA Broad Stem Cell Research Center Flow Cytometry Core for their expertise. This work used computational and storage services associated with the Hoffman2 Shared Cluster provided by UCLA Office of Advanced Research Computing’s Research Technology Group.

## Credits

GTC and DHG conceived the study, designed the experiments, and wrote the manuscript. CL and NES developed and characterized HUES66 isogenic CHD8^+/-^ and SCN2A^+/-^ cell lines. JM established the KOLF2.2J isogenic CHD8^+/-^ cell lines. GTC, GN, and AO characterized the mutant iPSC lines and established hCO cultures. GTC, GN, and AO performed immunofluorescence, microscopy, and image analysis of organoids. GTC and AO performed organoid dissociation and flow cytometry. GTC and KG performed Western blots. GTC and KG designed the guide RNA. GTC, GN, KG, JG, and YZ screened the gRNA in HEK293T and hNPCs. GN screened the gRNA in dCas9-p300 cortical organoids. GN, YZ, and AO treated mutant hCO with CRISPR-A. GTC, GN, AO, KG, and YZ performed qPCR analysis on CRISPR-A treated hCO. GTC, GN, AO, and YZ differentiated mutant lines into neurons. SH and CC conducted the patch clamp assays and analysis.

---

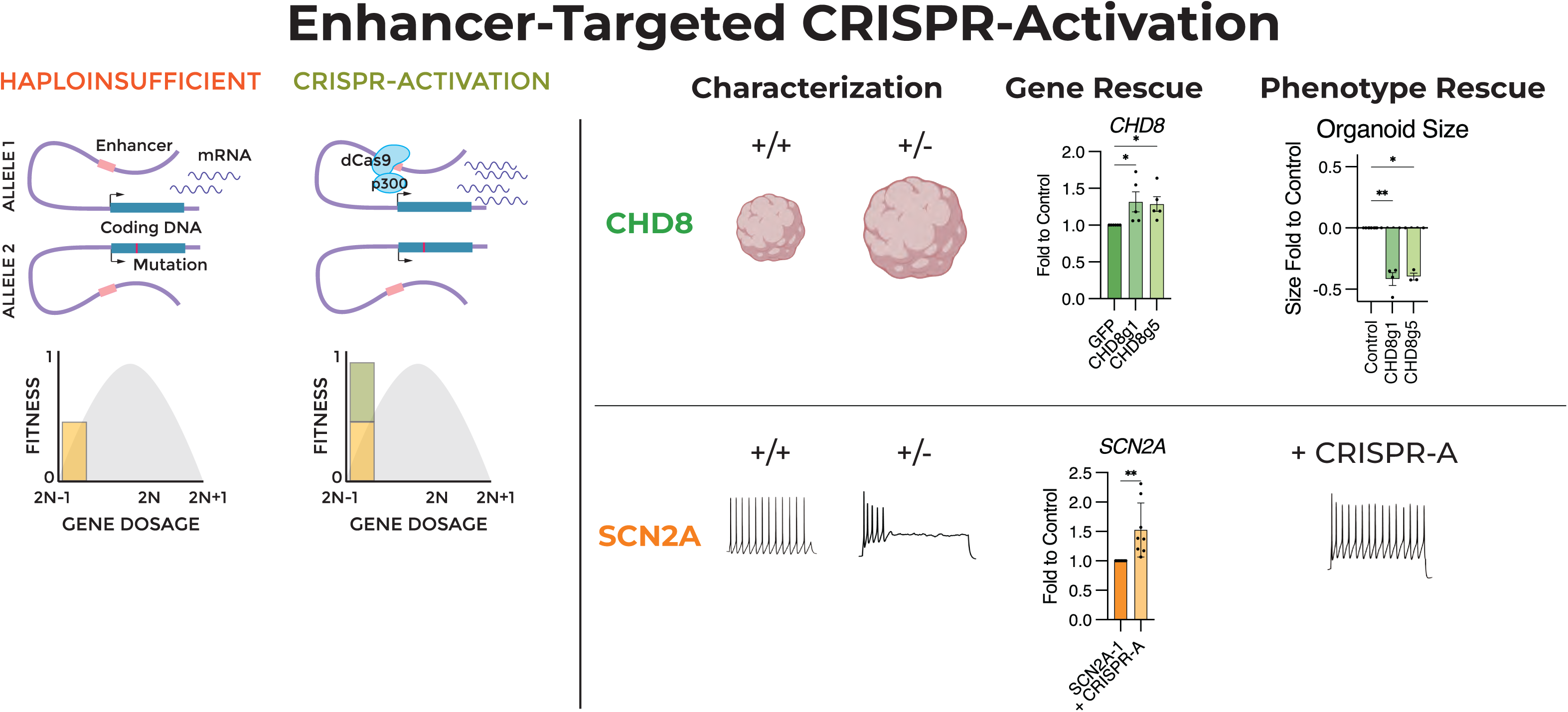

