## Supplementary material for "Enhancer-targeted CRISPR-A rescues haploinsufficiency and mutant phenotypes in organoid models of autism": All supplemental figures

**Supplemental Figure 1. Confirmation of mutations in CHD8 and SCN2A iPSC lines.** Sanger sequencing of CRISPR-edited regions of (A) CHD8 compared to HUES66 in CHD8-1A and CHD8-1B lines. (B) Sanger sequencing of KOLF2.2J CRISPR-edited lines CHD8-2A and CHD8-2B. (C) SCN2A edited in genome of HUES66 compared to SCN2A-1. (D) Sanger sequencing shows expected mutations in patient-derived lines SCN2A-2 and SCN2A-3 show expected deletions in mutant lines.

**Supplemental Figure 2. In vivo expression of CHD8 and SCN2A.** Expression of (A) *CHD8* and (B) *SCN2A* in the Allen Brainspan RNAseq dataset shows different time points of peak expression between the two genes of interest. (C) Expression of CHD8 and SCN2A in single cell fetal brain RNA sequencing from (Polioudakis et al. 2019) shows cell type specific expression of *SCN2A*.

**Supplemental Figure 3. Single cell RNA sequencing of CHD8 hCO reveal an increase of interneuron and interneuron progenitors compared to age-matched wildtype organoids.** Single cell data was downloaded from Paulsen et al. and re-annotated using Yoon et al cell type marker clusters. (A) tSNE plot of cell type clusters. (B) tSNE plot of wildtype versus mutant cells in single cell dataset. (C) Proportions of cell types show an increase in immature interneurons, interneuron progenitors, and cycling interneuron progenitors. N=3, SD shown.

**Supplemental Figure 4. Additional characterization of mutant hCO by qPCR and immunofluorescence.** (A) No difference in stem cell markers PAX6 and NES or neuron marker MAP2 observed between wildtype and mutant hCO. N = 3-6, SD shown. (B) Quantification of co-expression of CHD8 with cell type markers SOX2 and TBR2 shows significantly more CHD8+/TBR2+ cells in CHD8+/- hCO compared to wildtype, despite decreased overall number of CHD8+ cells (see figure 1E). N=4, SD shown, * $p < 0.05$ by Student's two-tailed t-test. (C) Gene Ontology of differentially expressed genes between $CHD8^{+/-}$ and HUES66 show an enrichment in neural progenitor cells and actively dividing cells in CHD8+/- hCO and an enrichment in neuron projection and oxidative phosphorylation, as a proxy for cell differentiation, in HUES66. (D) Gene Ontology of differentially expressed genes between SCN2A+/- and HUES66 show an enrichment of neural stem cell and neural progenitor cells in SCN2A hCO, calcium signaling and axoneme assembly in HUES66.

**Supplemental Figure 5. Putative enhancer sites for CHD8 and SCN2A.** Using the UCSC Genome Browser, enhancer annotations from the PsychENCODE Consortium (annotated enh##) and Human Fetal Brain Enhancers from de la Torre-Ubieta, et al. (annotated hfbe##) were mapped to hg19. Guide RNAs used in this study are marked on the left column next to the targeted enhancer sequence. The gene of interest is highlighted in red in the bottom.

**Supplemental Figure 6. Promoter-targeted CRISPR-A increases target gene expression in wildtype and haploinsufficient lines.** (A) GuideRNA targeting the *SCN2A* promoter significantly increases *SCN2A* expression in HEK293T cells. N = 3, SD shown, * $p < 0.05$. (B) CRISPR-A treatment of hCO targeting *CHD8* and *SCN2A* promoters in 25-day-old hCO increases target gene expression 1-month post-infection in both wildtype and haploinsufficient lines. N = 3-8, SEM shown, * $p < 0.05$. (C) Expression of target genes *CHD8* and *SCN2A* in iNGN2 neurons treated with promoter-targeted CRISPR-A. N=6, SEM shown, * p < 0.05. (D) Expression of *MAP2* is increased by promoter-targeted CRISPR-A for *CHD8* or *SCN2A* in both wildtype and haploinsufficient hCO. N = 3-8, SEM shown, * p < 0.05. (E) hCO size is reduced by CHD8 promoter-targeted CRISPR-A in both wildtype and $CHD8^{+/-}$ hCO. N = 5-10, SEM shown, * p < 0.05, ** p < 0.01, **** p < 0.0001.

**Supplemental Figure 7. Expression of CHD8 in CRISPR-A treated hCO normalized to wildtype lines.** Expression of *CHD8* in (A) CHD8-1A or (B) CHD8-1B hCO treated with CRISPR-A at 25-days old and measured 0-3 months post-infection is still less than the wildtype HUES66 line. (C) No significant difference was observed between wildtype lines HUES66 and KOLF2.2J in *CHD8* expression; addition of CRISPR-A to KOLF2.2J did not significantly change expression. Expression of *CHD8* in (D) CHD8-2A and (E) CHD8-2B treated with CRISPR-A compared to HUES66. For all, N = 8, SEM shown. (F) Expression of *CHD8* in the HUES66 wildtype line constitutively expressing dCas9-p300 shows no significant difference between control-treated and guideRNA treated hCO. (G) CHD8-1A constitutively expressing dCas9-p300 leads to rescue of *CHD8* expression to wildtype levels.

**Supplemental Figure 8. Expression of dCas9-p300 in CHD8-mutant neurons and organoids.** (A) Normalized expression of dCas9-p300 in iNGN2 neurons and cortical organoids shows strong expression of the construct at the earliest timepoint post-infection, decreasing across differentiation. Expression of dCas9-p300 in infected cortical organoids is very low, though significantly increased to vehicle-infected control. N=6-10, SD shown. * p < 0.05, ** p < 0.01, **** p < 0.0001. (B) Expression of GFP, as a proxy for gRNA expression, is not significantly changed during NGN2-driven differentiation or across organoid timepoints. N=6-10, SD shown. (C) Expression of Cas9 protein in hCO measured by immunofluorescence shows approximately 25% of cells expressed the dCas9-p300 construct. N=6-10, SD shown.

**Supplemental Figure 9. Extent of gene expression and electrophysiology changes in CRISPR-A treated $SCN2A^{+/-}$ hCO and neurons normalized to wildtype lines.** (A) Expression of SCN2A in SCN2A-1 hCO treated with CRISPR-A at 100 days old, normalized to HUES66 wildtype line showed increase in SCN2A expression to wildtype levels 2 months post-infection. N=3, SD shown. (B) Expression of neuron markers MAP2 and TUJ1 by immunoblot showed significant increases in both in CRISPR-A treated SCN2A+/- hCO; TUJ1 expression is increased to wildtype levels. N = 6, SEM shown. * p < 0.05, ** p < 0.01, **** p < 0.0001 by Student's Two-Tailed t-test. (C) Voltage clamp recordings of intrinsic currents in response to step voltage commands (10 mV steps from -80 to +10 mV). Depolarizing voltage commands induced large $Na^+$ currents of variable amplitudes (arrow). (D) Sample action potential responses evoked by 5 ms depolarizing current injections. Cells were held at -60 mV prior to current pulses . Rheobase was defined as the minimum current required to elicit a single action potential, determined using incremental 5 pA current pulses . (E) Summary plots of passive cell membrane properties (capacitance, input resistance), $Na^+$ current amplitudes, RMPs, rheobase, and action potential firing threshold across wildtype controls, $SCN2A^{+/-}$ and $SCN2A^{+/-}$ +CRISPR-A expressing cells. $SCN2A\text{-}1^{+/–}$ neurons were compared to the HUES66 wildtype line and analyzed independently, whereas patient-derived $SCN2A\text{-}2^{+/–}$ and $SCN2A\text{-}3^{+/–}$ neurons were compared to the KOLF2.2J wildtype line and pooled for analysis; corresponding CRISPR-A lines were combined similarly. Mean values ±SEM are shown;

N=11 (HUES66), N=14 (SCN2A-1$^{+/-}$), N=17 (SCN2A-1$^{+/-}$ +CRISPR-A), N=13 (KOLF2.2J), N=29 (SCN2A-2,3$^{+/-}$), and N=17 (SCN2A-2,3$^{+/-}$ +CRISPR-A). Statistical significance was determined by one-way ANOVA followed by Tukey's multiple-comparisons test, with mutant and CRISPR-A groups compared to their matched wildtype controls; * $p < 0.05$, ** $p < 0.01$, **** $p < 0.0001$. (E) Expression of sodium channel subunits in wildtype and mutant lines shows that patient-derived SCN2A-mutant iPSC lines SCN2A-2 and SCN2A-3 show increased expression of SCN1A, SCN3A, and SCN5A compared to HUES66-derived SCN2A-1. N = 6, SEM shown, * $p < 0.05$ by Student's Two-tailed t-test.

**Supplemental Figure 10. Applying CRISPR-A at alternative timepoints for CHD8 and SCN2A haploinsufficient organoids does not increase target gene expression.** (A) Treatment of HUES66 hCO with CHD8- or SCN2A-targeted CRISPR-A in 50-day old hCO (late for CHD8, early for SCN2A) does not significantly change target gene expression 1-month post-infection. N = 4, SEM shown, p-value by One-Way ANOVA. (B) Expression of CHD8 in 50- or 100-day old CHD8-1A hCO transduced with CRISPR-A and collected 1-month post-infection shows no significant increase. N = 4-7, SEM shown, p-values by One-Way ANOVA. (C) Expression of SCN2A in 50-day old SCN2A-1 hCO treated with CRISPR-A and collected 1-month post infection shows no significant increase. (D) Expression of CHD8 in 300-day old HUES66 and CHD8-1A hCO treated with CRISPR-A and collected 1-month post infection. N = 3, SD shown.

**Supplemental Figure 11. Guide RNA optimization for CRISPR editing does not optimize for gene activation.** (A) UCSC Genome Browser window mapping CHD8 enhancers, guides, and histone accessibility data from PsychENCODE. (B) UCSC Genome Brower window mapping SCN2A enhancers, guides, and histone accessibility data from PsychENCODE.

**Supplemental Figure 12.** Full Western blot images.

**Supplemental Figure 13.** Flow cytometry gating strategy

**Supplemental Figure 14.** Full immunofluorescence images.

**Supplemental Table 1. Guide RNA optimization for CRISPR editing.** Summary of guide RNA efficiency optimizations reviewed in Konstantakos et al. on tested guides.

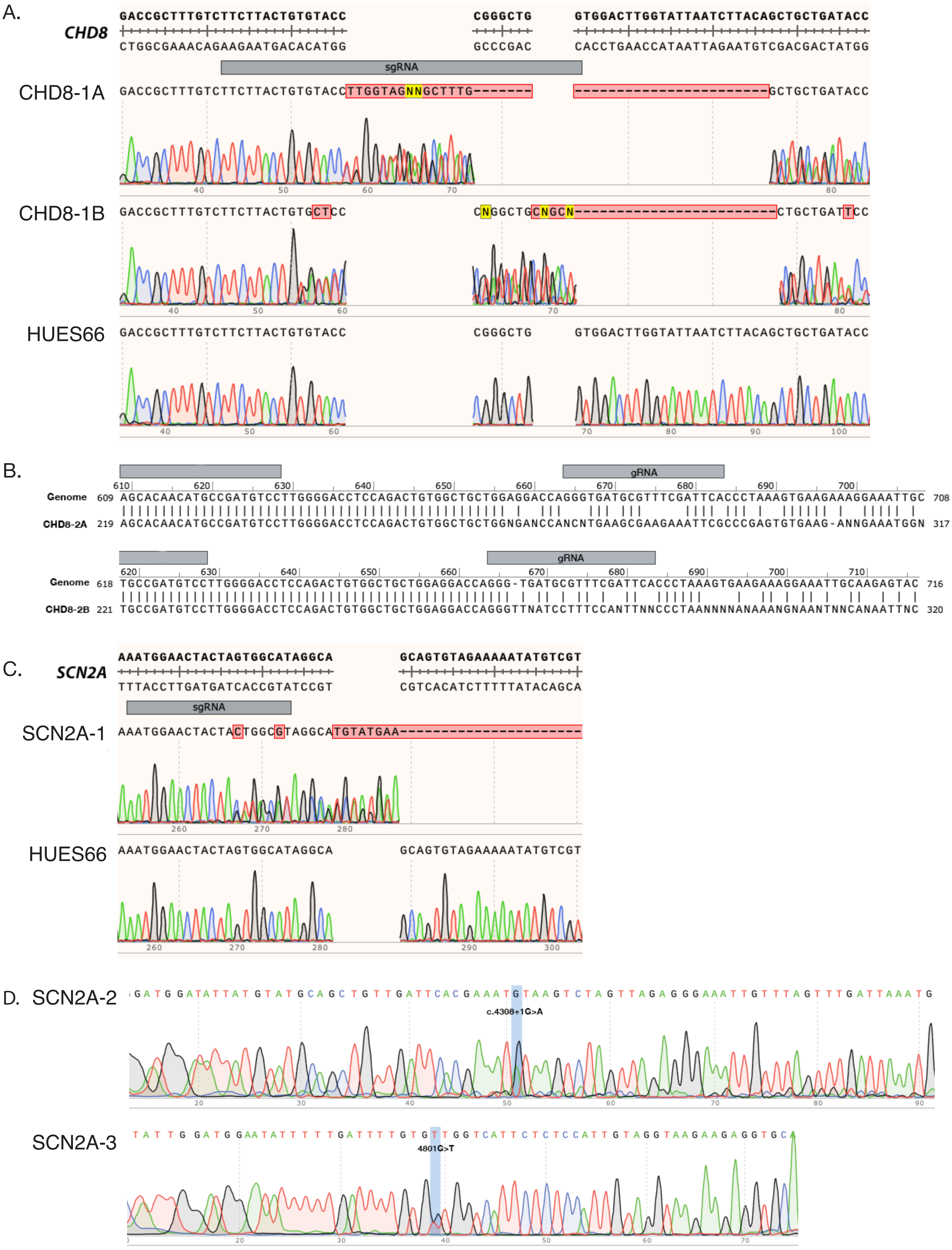

A.
CHD8
GACCGCTTTGTCTTCTTACTGTGTACC
CGGGCTG
GTGGACTTGGTATTAATCTTACAGCTGCTGATACC
CTGGCGAAACAGAAGAATGACACATGG
GCCCGAC
CACCTGAACCATAATTAGAATGTCGACGACTATGG
sgRNA
CHD8-1A
GACCGCTTTGTCTTCTTACTGTGTACCTTGGTAGNNGCTTTG-------
------------------------GCTGCTGATACC
CHD8-1B
GACCGCTTTGTCTTCTTACTGTGCTCC
CNGGCTGCNGCN-------------------------CTGCTGATTCC
HUES66
GACCGCTTTGTCTTCTTACTGTGTACC
CGGGCTG
GTGGACTTGGTATTAATCTTACAGCTGCTGATACC
B.
gRNA
Genome 609 AGCACAACATGCCGATGTCCTTGGGGACCTCCAGACTGTGGCTGCTGGAGGACCAGGGTGATGCGTTTCGATTCACCCTAAAGTGAAGAAAGGAAATTGC 708
CHD8-2A 219 AGCACAACATGCCGATGTCCTTGGGGACCTCCAGACTGTGGCTGCTGGNGANCCANCNTGAAGCGAAGAAATTCGCCCGAGTGTGAAG-ANNGAAATGGN 317
gRNA
Genome 618 TGCCGATGTCCTTGGGGACCTCCAGACTGTGGCTGCTGGAGGACCAGGG-TGATGCGTTTCGATTCACCCTAAAGTGAAGAAAGGAAATTGCAAGAGTAC 716
CHD8-2B 221 TGCCGATGTCCTTGGGGACCTCCAGACTGTGGCTGCTGGAGGACCAGGGTTNATCCTTTCCANTTNNCCCTAANNNNANAAANGNAANTNNCANAATTNC 320
C.
SCN2A
AAATGGAACTACTAGTGGCATAGGCA
GCAGTGTAGAAAAATATGTCGT
TTTACCTTGATGATCACCGTATCCGT
CGTCACATCTTTTTATACAGCA
sgRNA
SCN2A-1
AAATGGAACTACTACTGGCGTAGGCATGTATGAA---------------------
HUES66
AAATGGAACTACTAGTGGCATAGGCA
GCAGTGTAGAAAAATATGTCGT
D.
SCN2A-2
GATGGATATTATGTATGCAGCTGTTGATTCACGAAATGTAAGTCTAGTTAGAGGGAAATTGTTTAGTTTGATTAAATG
c.4308+1G>A
SCN2A-3
TATTGGATGGAATATTTTTGATTTTGTGTTGGTCATTCTCTCCATTGTAGGTAAGAAGAGGTGCA
4801G>T

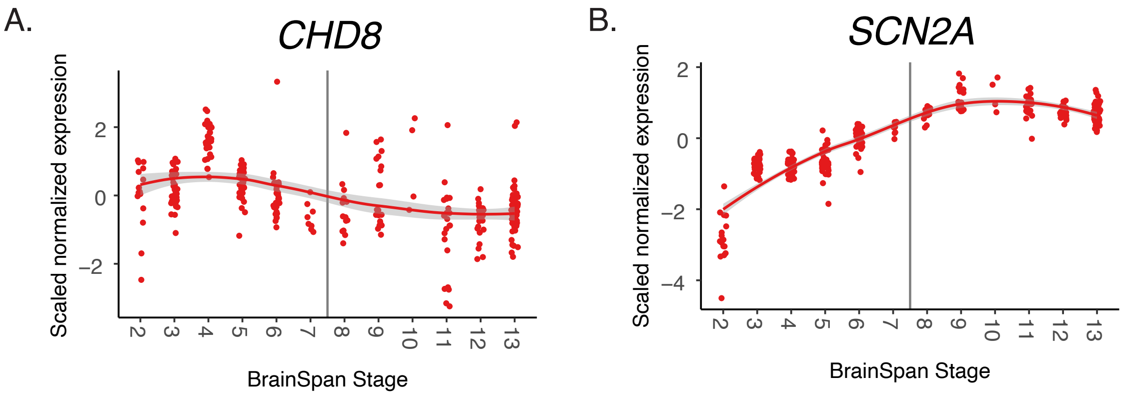


| Stage | Age |
|---|---|
| 1 | 4-7 pcw |
| 2 | 8-12 pcw |
| 3 | 10-12 pcw |
| 4 | 13-15 pcw |
| 5 | 16-18 pcw |
| 6 | 19-24 pcw |
| 7 | 25-38 pcw |
| 8 | Birth-5 months |
| 9 | 6-18 months |
| 10 | 19 months-5 yrs |
| 11 | 6-11 yrs |
| 12 | 12-19 yrs |
| 13 | 20-60+ yrs |

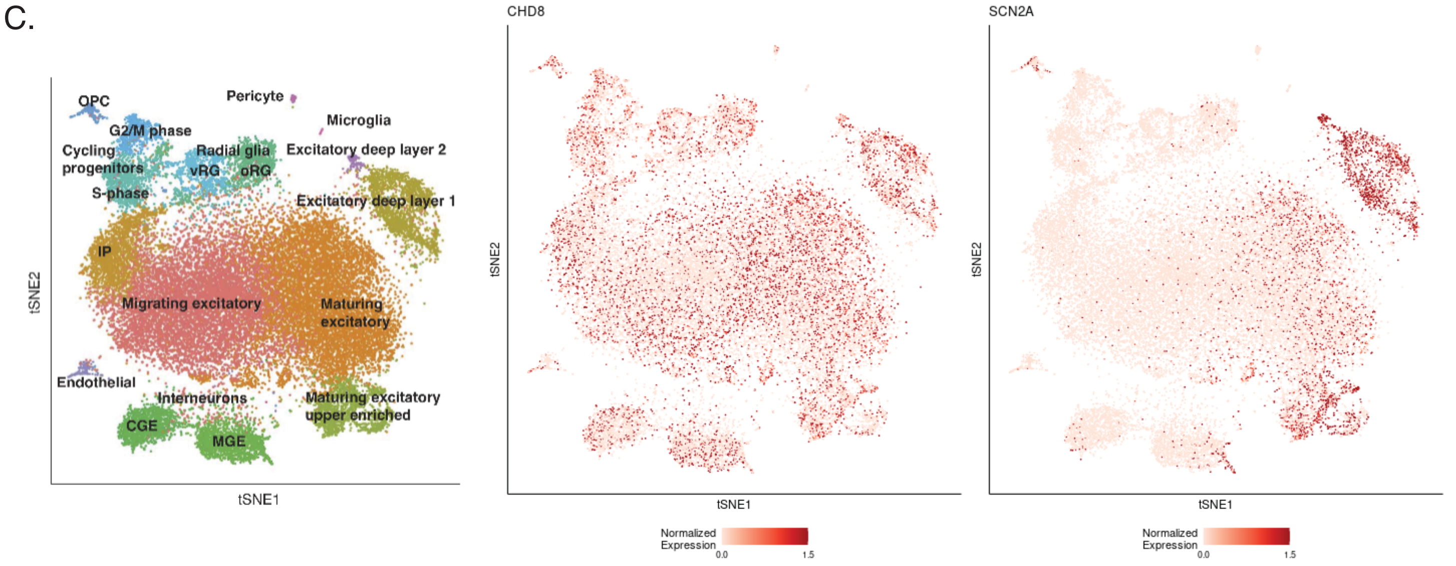

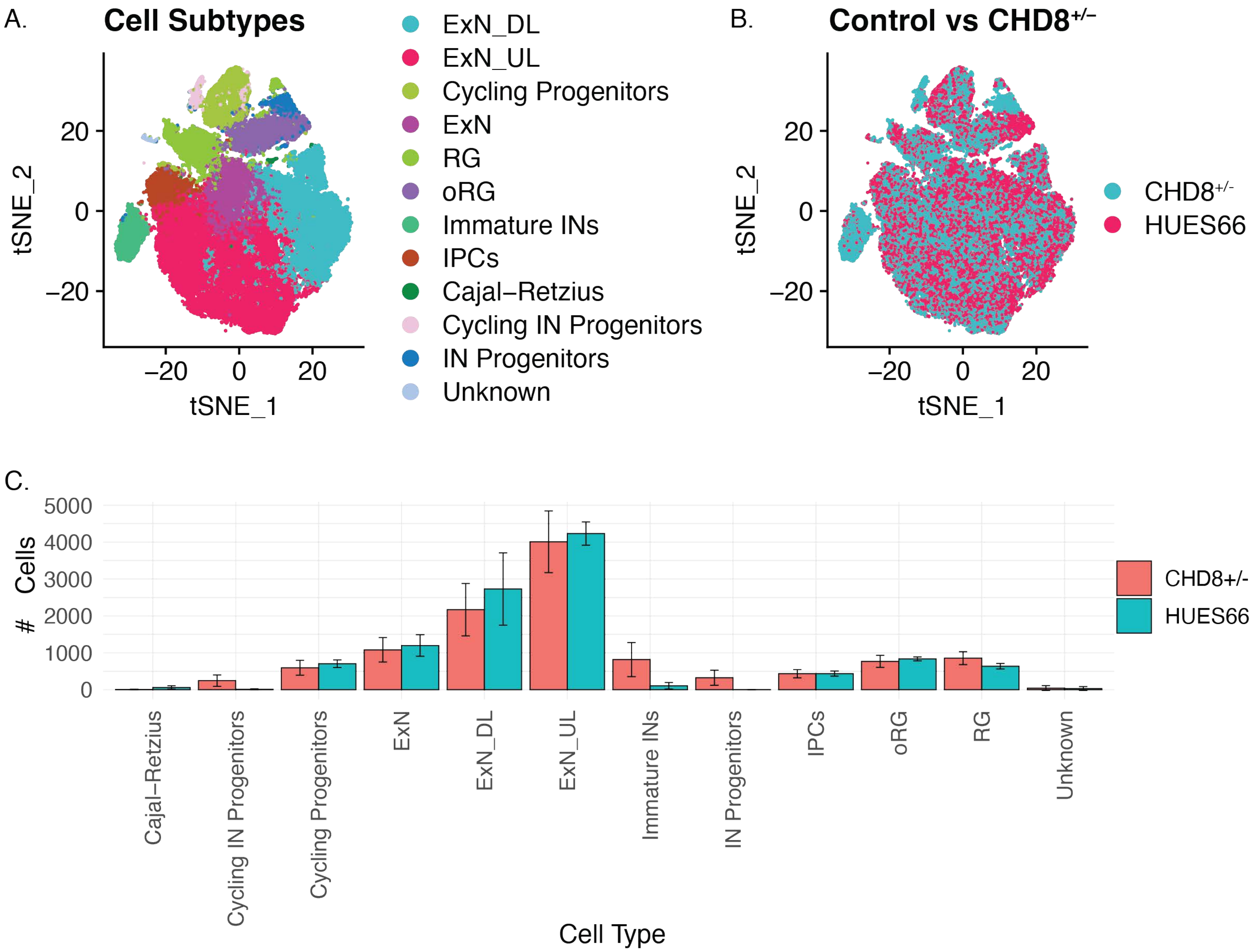

A.
Cell Subtypes
ExN_DL
ExN_UL
Cycling Progenitors
ExN
RG
oRG
Immature INs
IPCs
Cajal–Retzius
Cycling IN Progenitors
IN Progenitors
Unknown
tSNE_2
tSNE_1
B.
Control vs CHD8+/–
CHD8+/-
HUES66
C.
### Cells
5000
4000
3000
2000
1000
0
CHD8+/-
HUES66
Cajal–Retzius
Cycling IN Progenitors
Cycling Progenitors
ExN
ExN_DL
ExN_UL
Immature INs
IN Progenitors
IPCs
oRG
RG
Unknown
Cell Type

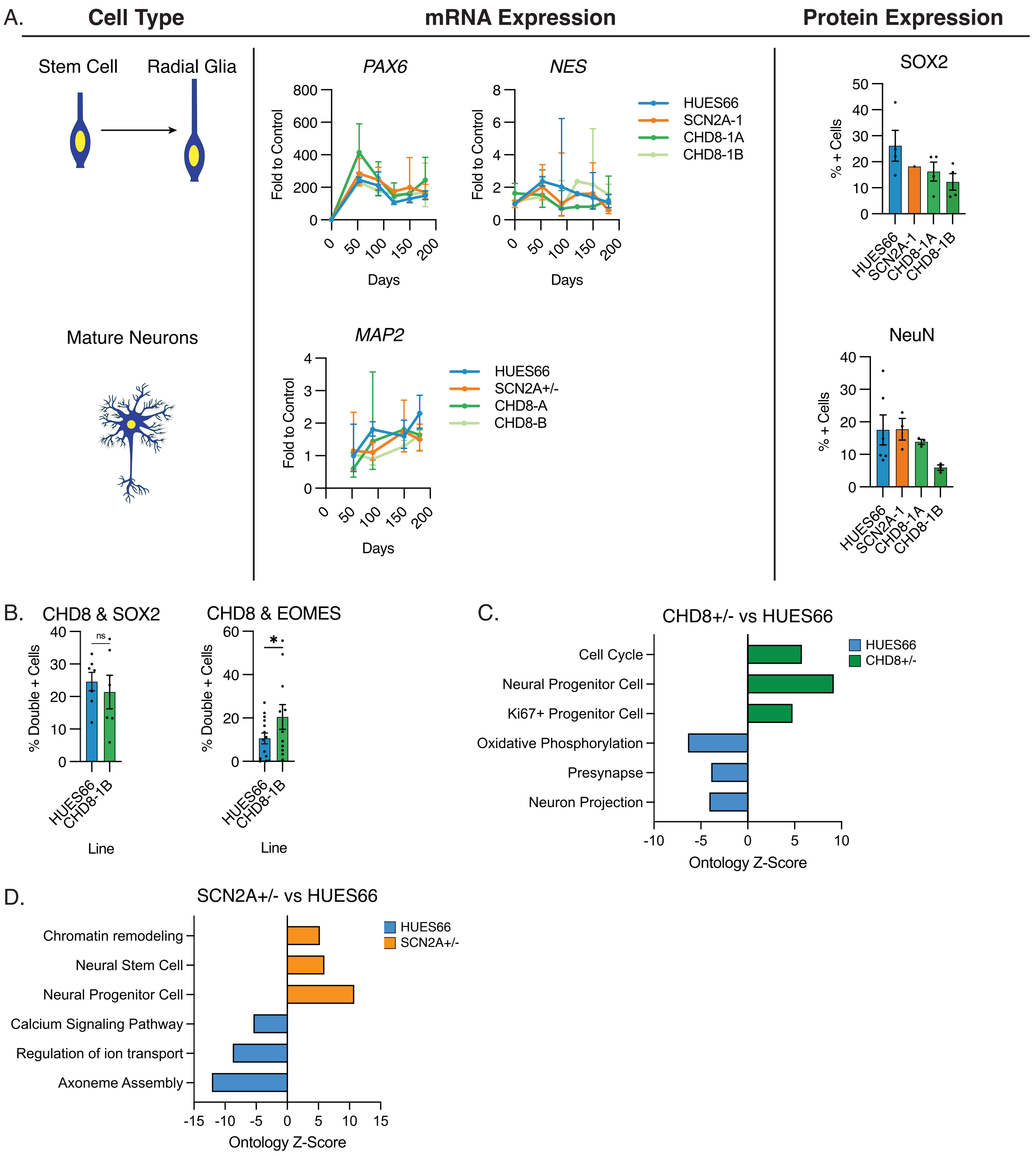
A.
Cell Type
mRNA Expression
Protein Expression
Stem Cell
Radial Glia
Mature Neurons
PAX6
NES
MAP2
Fold to Control
Days
HUES66
SCN2A-1
CHD8-1A
CHD8-1B
SCN2A+/-
CHD8-A
CHD8-B
SOX2
NeuN
% + Cells
B.
CHD8 & SOX2
CHD8 & EOMES
% Double + Cells
ns
*
Line
C.
CHD8+/- vs HUES66
Cell Cycle
Neural Progenitor Cell
Ki67+ Progenitor Cell
Oxidative Phosphorylation
Presynapse
Neuron Projection
CHD8+/-
Ontology Z-Score
D.
SCN2A+/- vs HUES66
Chromatin remodeling
Neural Stem Cell
Neural Progenitor Cell
Calcium Signaling Pathway
Regulation of ion transport
Axoneme Assembly
SCN2A+/-
Ontology Z-Score

A. CHD8

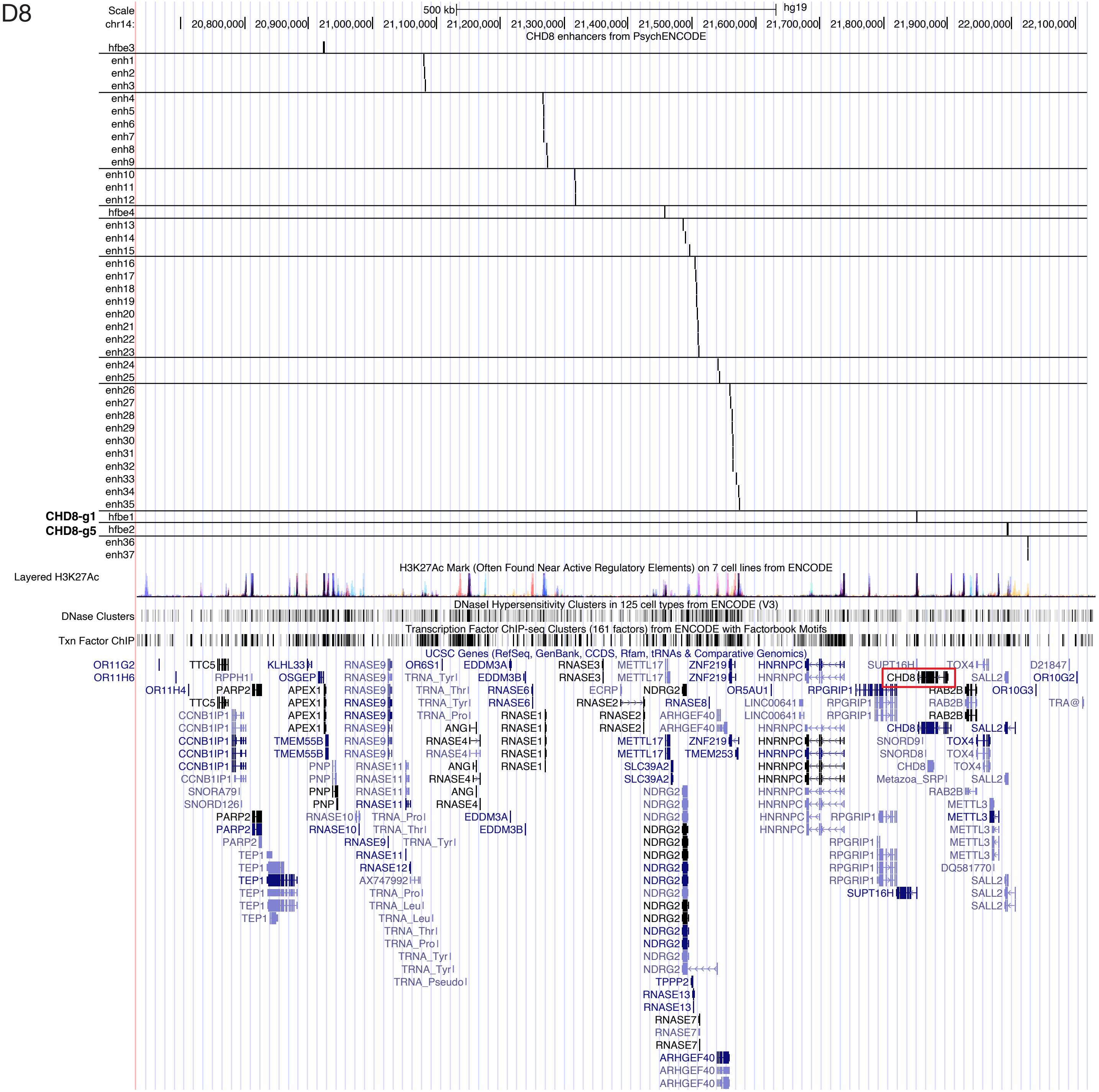


B. SCN2A

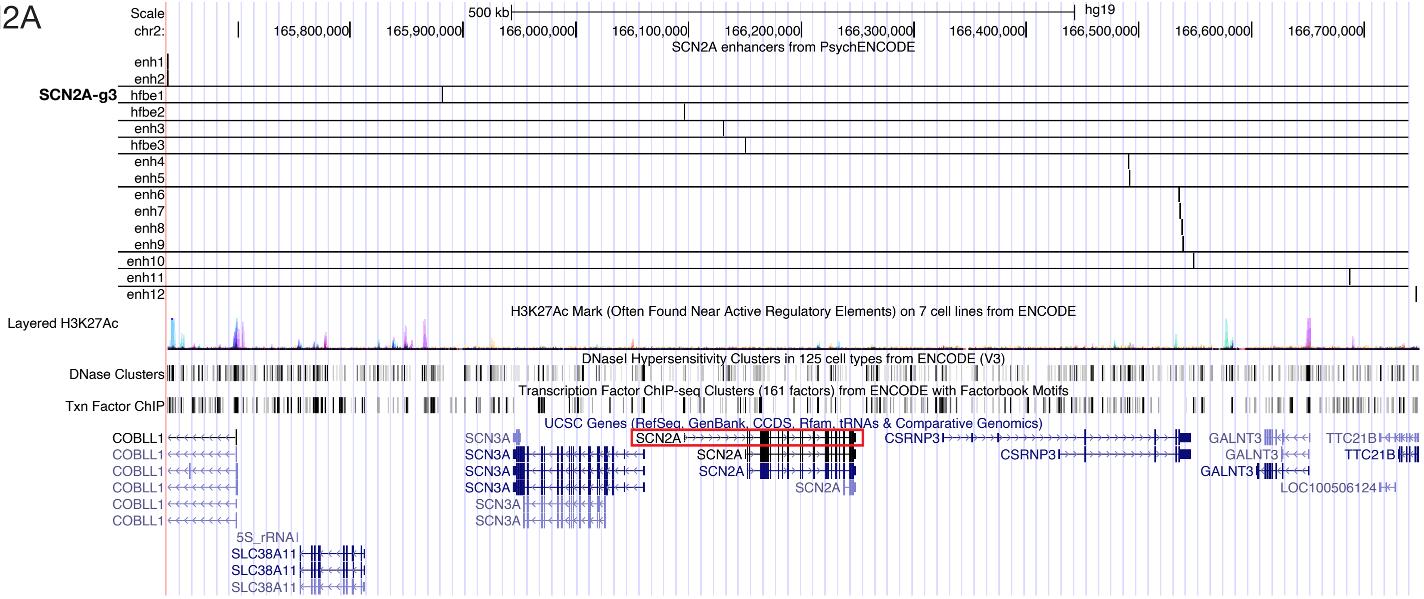

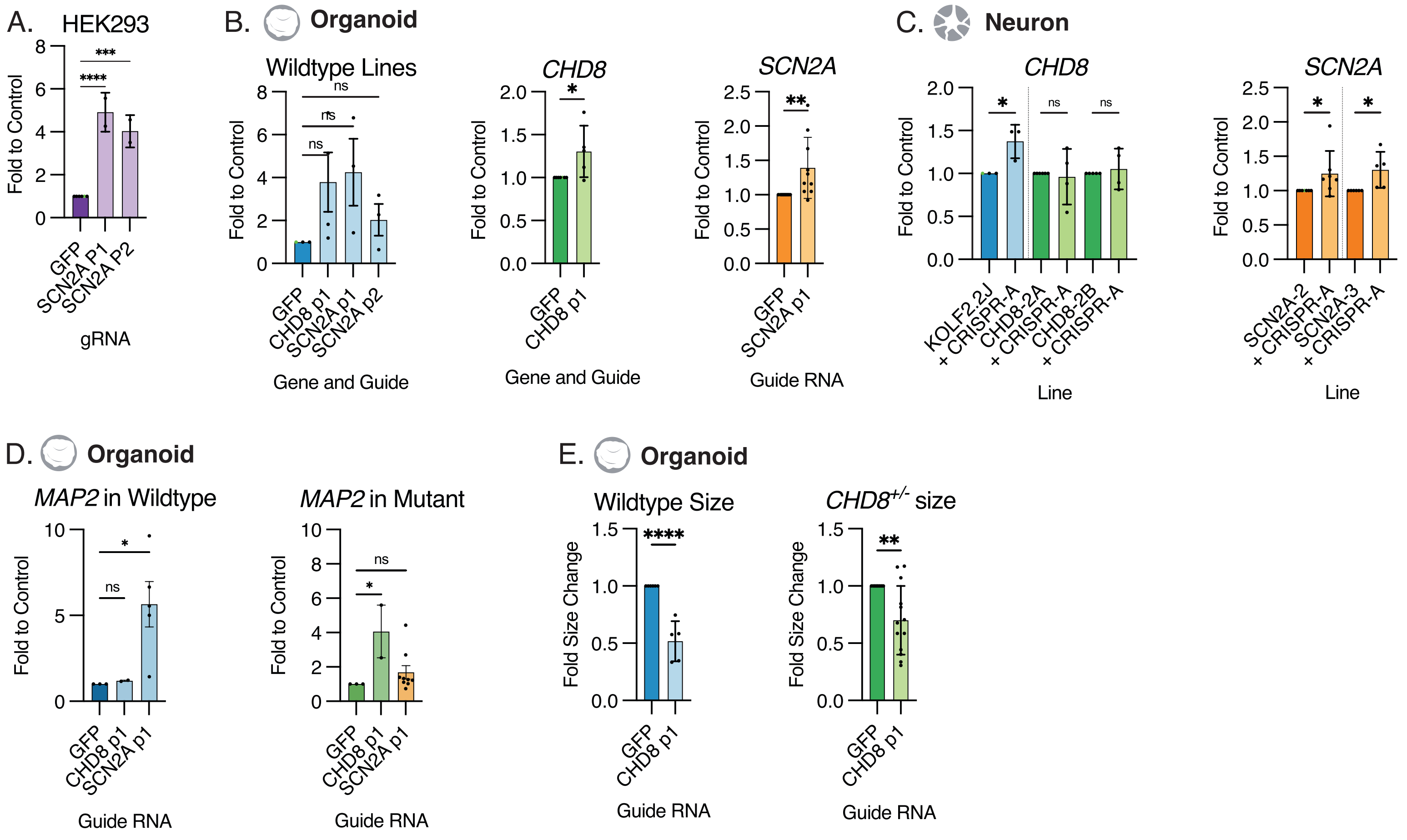
A. HEK293
Fold to Control
GFP
SCN2A P1
SCN2A P2
gRNA
****
***
B. Organoid
Wildtype Lines
Fold to Control
GFP
CHD8 p1
SCN2A p1
SCN2A p2
Gene and Guide
ns
CHD8
GFP
CHD8 p1
Gene and Guide
*
SCN2A
GFP
SCN2A p1
Guide RNA
**
C. Neuron
CHD8
Fold to Control
KOLF2.2J
× CRISPR-A
CHD8-2A
× CRISPR-A
CHD8-2B
× CRISPR-A
Line
SCN2A
SCN2A-2
× CRISPR-A
SCN2A-3
× CRISPR-A
Line
D. Organoid
MAP2 in Wildtype
Fold to Control
GFP
CHD8 p1
SCN2A p1
Guide RNA
MAP2 in Mutant
Guide RNA
E. Organoid
Wildtype Size
Fold Size Change
GFP
CHD8 p1
Guide RNA
****
CHD8+/- size
Guide RNA

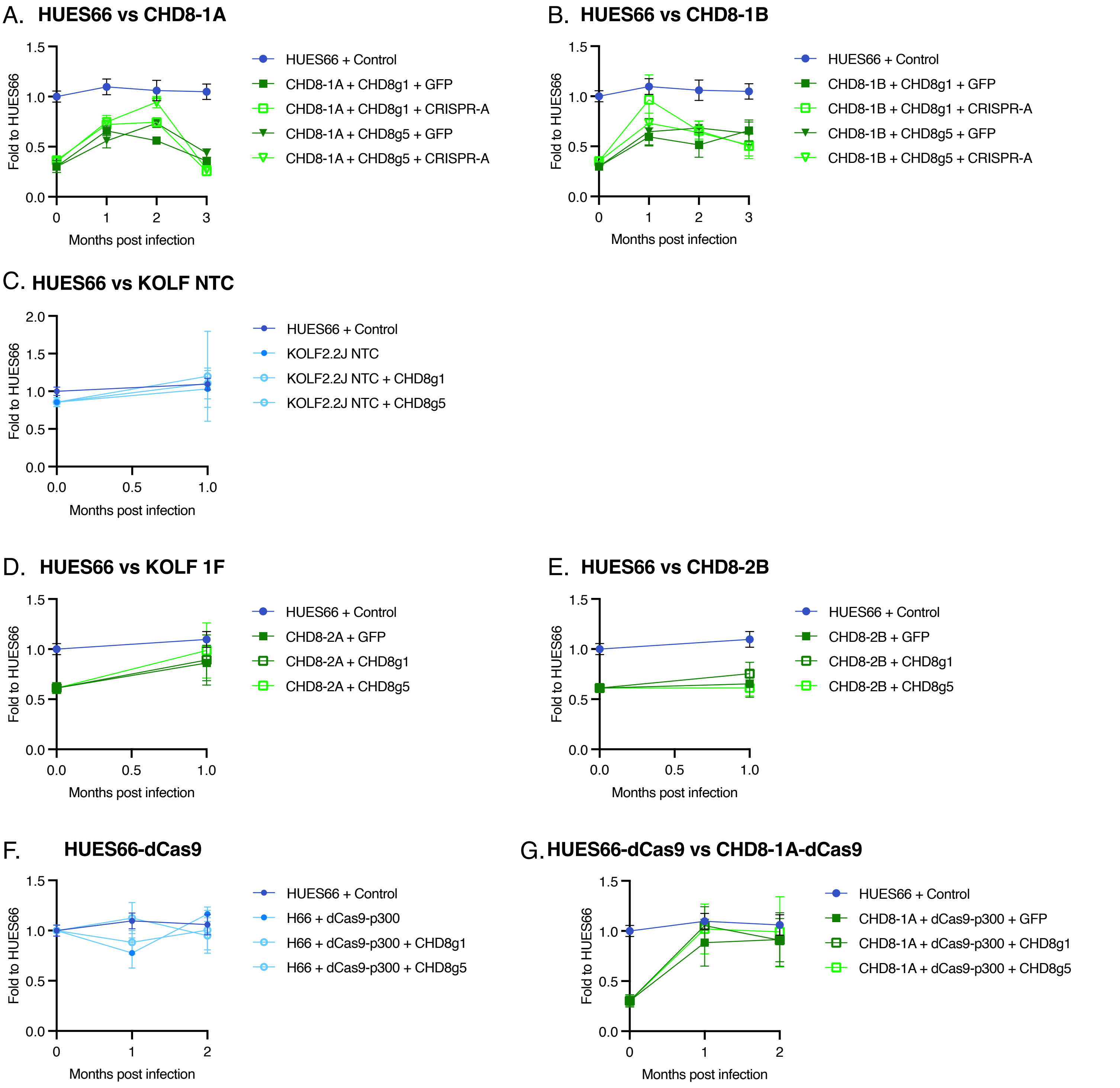

A. HUES66 vs CHD8-1A
Fold to HUES66
Months post infection
HUES66 + Control
CHD8-1A + CHD8g1 + GFP
CHD8-1A + CHD8g1 + CRISPR-A
CHD8-1A + CHD8g5 + GFP
CHD8-1A + CHD8g5 + CRISPR-A
B. HUES66 vs CHD8-1B
Fold to HUES66
Months post infection
HUES66 + Control
CHD8-1B + CHD8g1 + GFP
CHD8-1B + CHD8g1 + CRISPR-A
CHD8-1B + CHD8g5 + GFP
CHD8-1B + CHD8g5 + CRISPR-A
C. HUES66 vs KOLF NTC
Fold to HUES66
Months post infection
HUES66 + Control
KOLF2.2J NTC
KOLF2.2J NTC + CHD8g1
KOLF2.2J NTC + CHD8g5
D. HUES66 vs KOLF 1F
Fold to HUES66
Months post infection
HUES66 + Control
CHD8-2A + GFP
CHD8-2A + CHD8g1
CHD8-2A + CHD8g5
E. HUES66 vs CHD8-2B
Fold to HUES66
Months post infection
HUES66 + Control
CHD8-2B + GFP
CHD8-2B + CHD8g1
CHD8-2B + CHD8g5
F. HUES66-dCas9
Fold to HUES66
Months post infection
HUES66 + Control
H66 + dCas9-p300
H66 + dCas9-p300 + CHD8g1
H66 + dCas9-p300 + CHD8g5
G. HUES66-dCas9 vs CHD8-1A-dCas9
Fold to HUES66
Months post infection
HUES66 + Control
CHD8-1A + dCas9-p300 + GFP
CHD8-1A + dCas9-p300 + CHD8g1
CHD8-1A + dCas9-p300 + CHD8g5

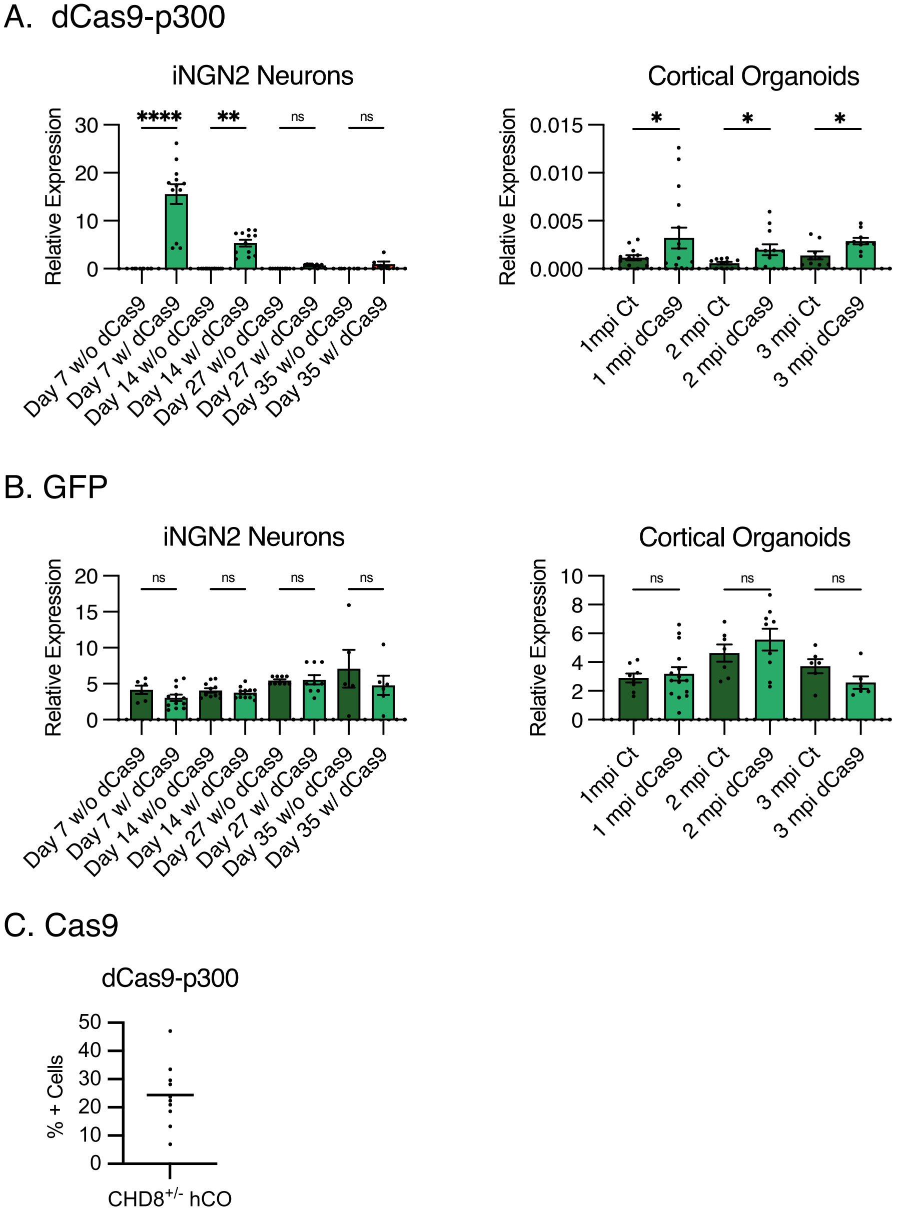
A. dCas9-p300
iNGN2 Neurons
Relative Expression
0
10
20
30
****
**
ns
ns
Day 7 w/o dCas9
Day 7 w/ dCas9
Day 14 w/o dCas9
Day 14 w/ dCas9
Day 27 w/o dCas9
Day 27 w/ dCas9
Day 35 w/o dCas9
Day 35 w/ dCas9
Cortical Organoids
Relative Expression
0.000
0.005
0.010
0.015
*
*
*
1mpi Ct
1 mpi dCas9
2 mpi Ct
2 mpi dCas9
3 mpi Ct
3 mpi dCas9
B. GFP
iNGN2 Neurons
Relative Expression
0
5
10
15
20
ns
ns
ns
ns
Day 7 w/o dCas9
Day 7 w/ dCas9
Day 14 w/o dCas9
Day 14 w/ dCas9
Day 27 w/o dCas9
Day 27 w/ dCas9
Day 35 w/o dCas9
Day 35 w/ dCas9
Cortical Organoids
Relative Expression
0
2
4
6
8
10
ns
ns
ns
1mpi Ct
1 mpi dCas9
2 mpi Ct
2 mpi dCas9
3 mpi Ct
3 mpi dCas9
C. Cas9
dCas9-p300
% + Cells
0
10
20
30
40
50
CHD8+/- hCO

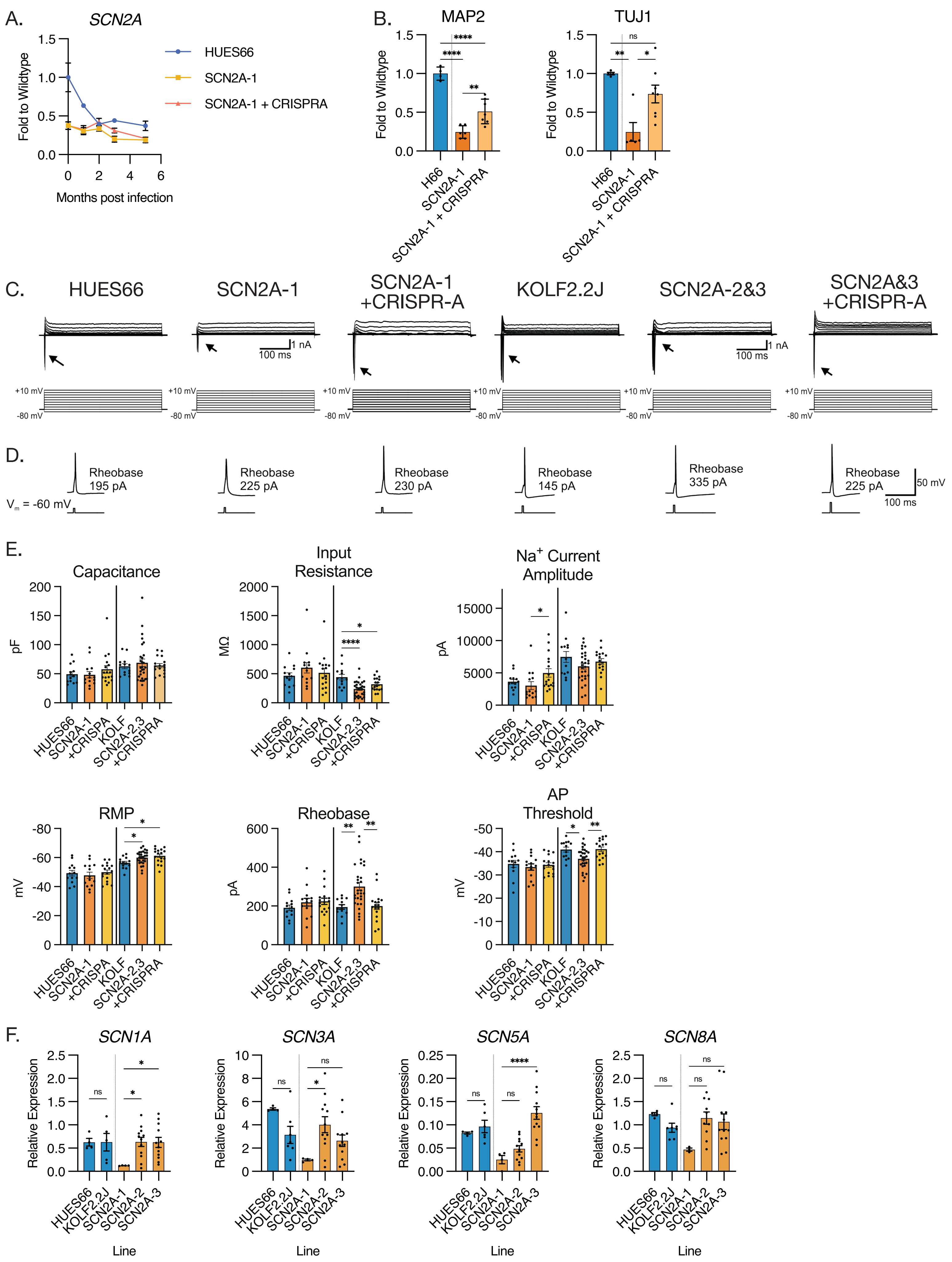

A.
SCN2A
Fold to Wildtype
Months post infection
HUES66
SCN2A-1
SCN2A-1 + CRISPRA
B.
MAP2
TUJ1
Fold to Wildtype
H66
SCN2A-1
SCN2A-1 + CRISPRA
C.
HUES66
SCN2A-1
SCN2A-1 +CRISPR-A
KOLF2.2J
SCN2A-2&3
SCN2A&3 +CRISPR-A
1 nA
100 ms
+10 mV
-80 mV
D.
Vm = -60 mV
Rheobase 195 pA
Rheobase 225 pA
Rheobase 230 pA
Rheobase 145 pA
Rheobase 335 pA
Rheobase 225 pA
50 mV
100 ms
E.
Capacitance
Input Resistance
Na+ Current Amplitude
RMP
Rheobase
AP Threshold
pF
MΩ
pA
mV
HUES66
SCN2A-1
+CRISPA
KOLF
SCN2A-2,3
+CRISPRA
F.
SCN1A
SCN3A
SCN5A
SCN8A
Relative Expression
HUES66
KOLF2.2J
SCN2A-1
SCN2A-2
SCN2A-3
Line

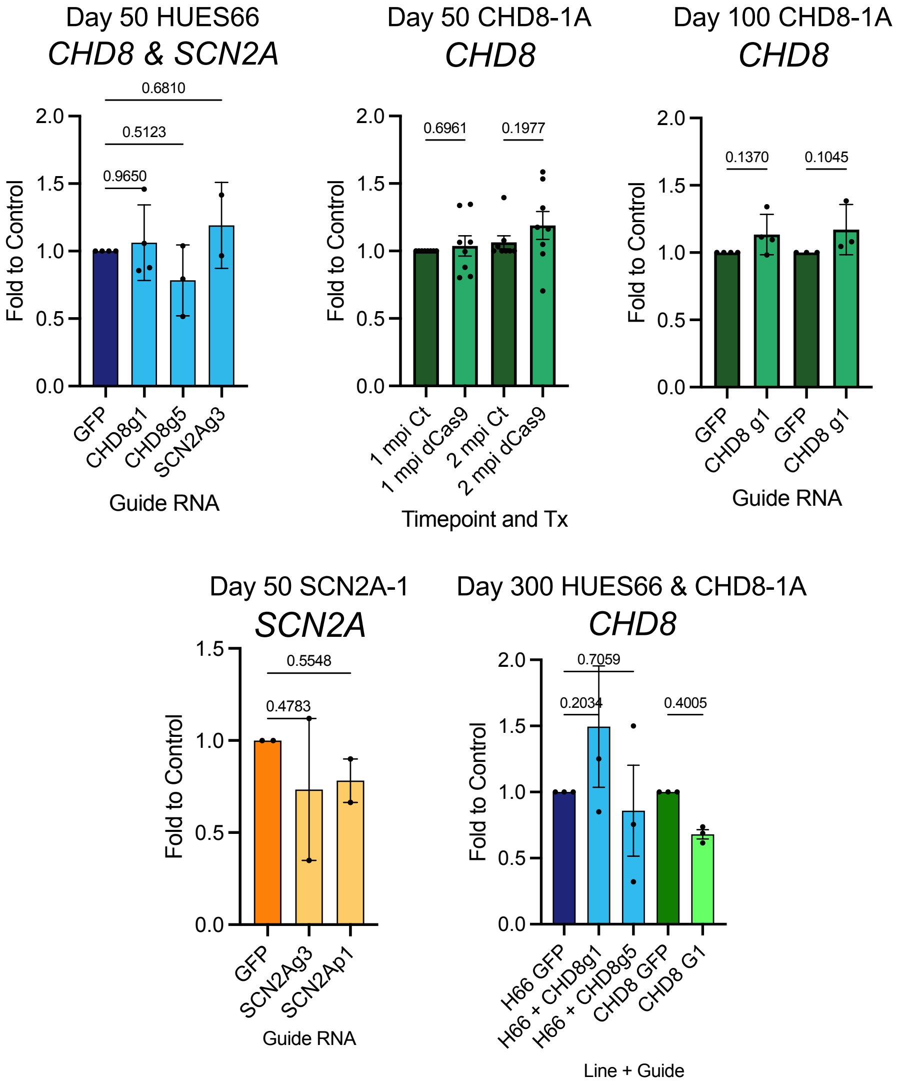

Day 50 HUES66
*CHD8 & SCN2A*
0.6810
0.5123
0.9650
Fold to Control
2.0
1.5
1.0
0.5
0.0
GFP
CHD8g1
CHD8g5
SCN2Ag3
Guide RNA
Day 50 CHD8-1A
*CHD8*
0.6961
0.1977
Fold to Control
1 mpi Ct
1 mpi dCas9
2 mpi Ct
2 mpi dCas9
Timepoint and Tx
Day 100 CHD8-1A
*CHD8*
0.1370
0.1045
Fold to Control
GFP
CHD8 g1
GFP
CHD8 g1
Guide RNA
Day 50 SCN2A-1
*SCN2A*
0.5548
0.4783
Fold to Control
GFP
SCN2Ag3
SCN2Ap1
Guide RNA
Day 300 HUES66 & CHD8-1A
*CHD8*
0.7059
0.2034
0.4005
Fold to Control
H66 GFP
H66 + CHD8g1
H66 + CHD8g5
CHD8 GFP
CHD8 G1
Line + Guide

#### **A.** Guide RNA efficiency optimization

| Guide | PAM | % GC | G/A in position 20 | Consecutive G's | %GC in positions 4-8 | % GC in positions 15-20 | G/A in position 19 | C in position 16 or 18 |
|---|---|---|---|---|---|---|---|---|
| Optimal | | 40-60% | Yes | Minimal | High | High | Yes | Yes |
| CHD8-g1 | TGG | 47.8 | Y | 3 | 40 | 0 | N | N |
| CHD8-g2 | AGG | 47.8 | N | 2 | 60 | 20 | Y | N |
| CHD8-g3 | TGG | 60.9 | N | 2 | 80 | 40 | Y | N |
| CHD8-g4 | TGG | 60.9 | N | 2 | 80 | 40 | Y | Y |
| CHD8-g5 | AGG | 56.5 | Y | 3 | 60 | 60 | N | Y |
| CHD8-g6 | GGG | 65.2 | Y | 4 | 80 | 80 | Y | N |
| SCN2A-g1 | TGG | 39.1 | Y | 1 | 40 | 20 | N | N |
| SCN2A-g2 | AGG | 43.5 | N | 1 | 40 | 60 | N | Y |
| SCN2A-g3 | TGG | 39.1 | N | 1 | 20 | 40 | N | N |

**B.**

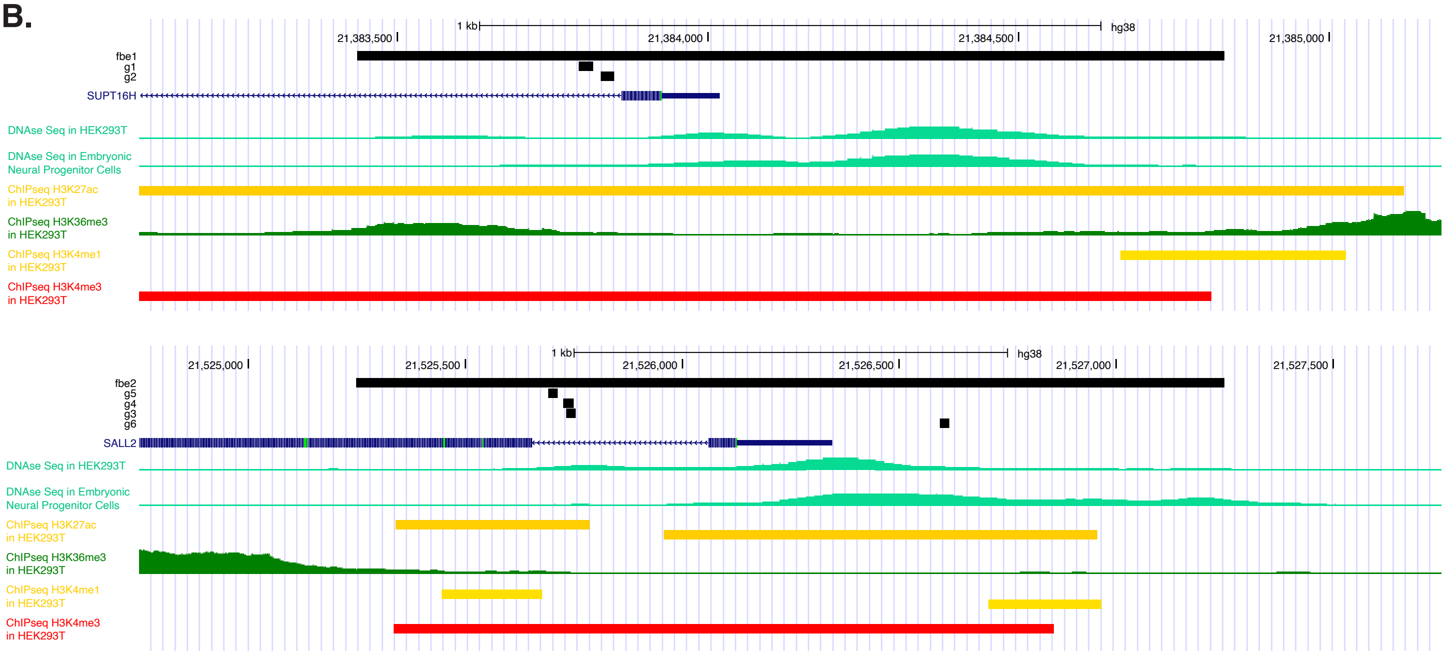


**C.**

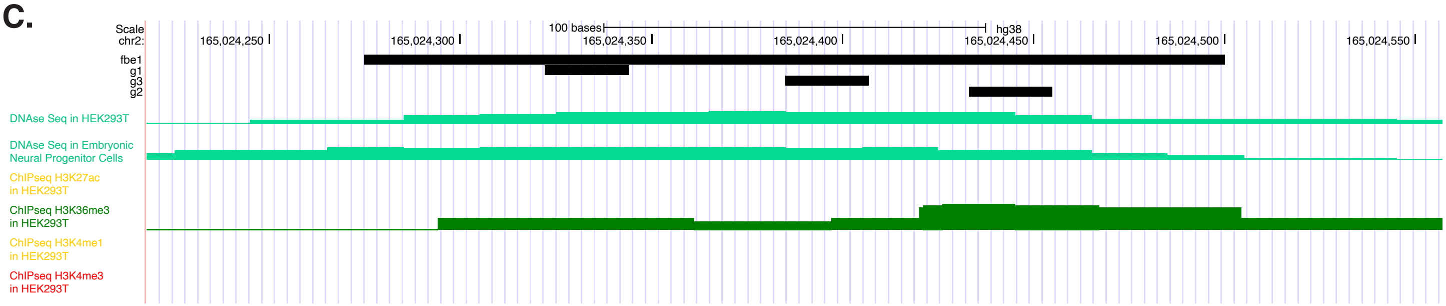

Figure 1. CHD8

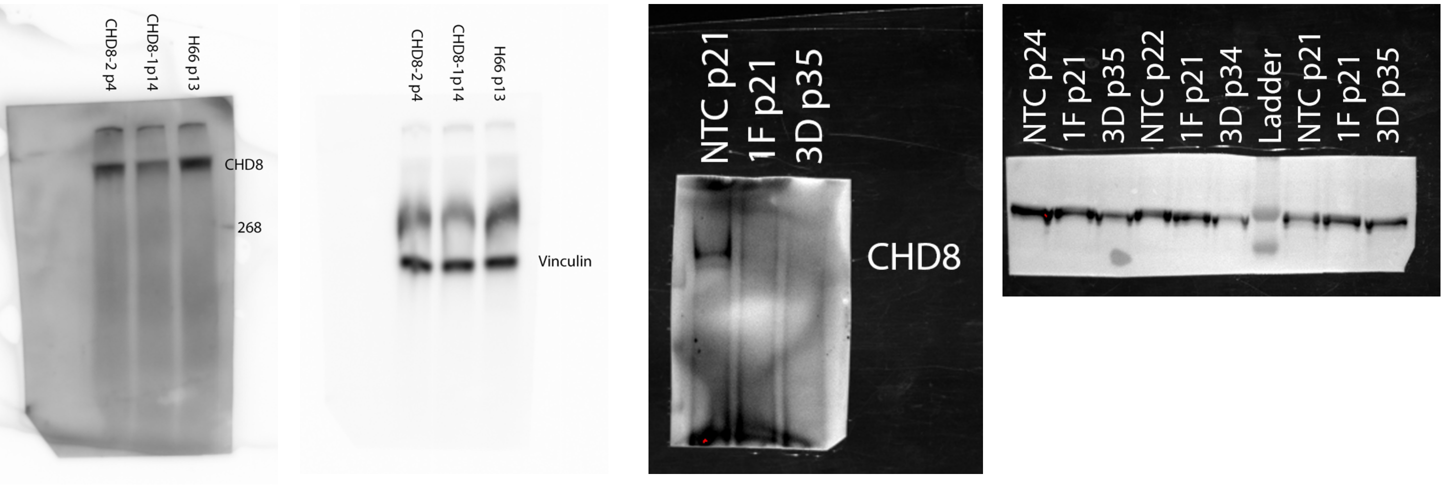


Figure 2. SCN2A

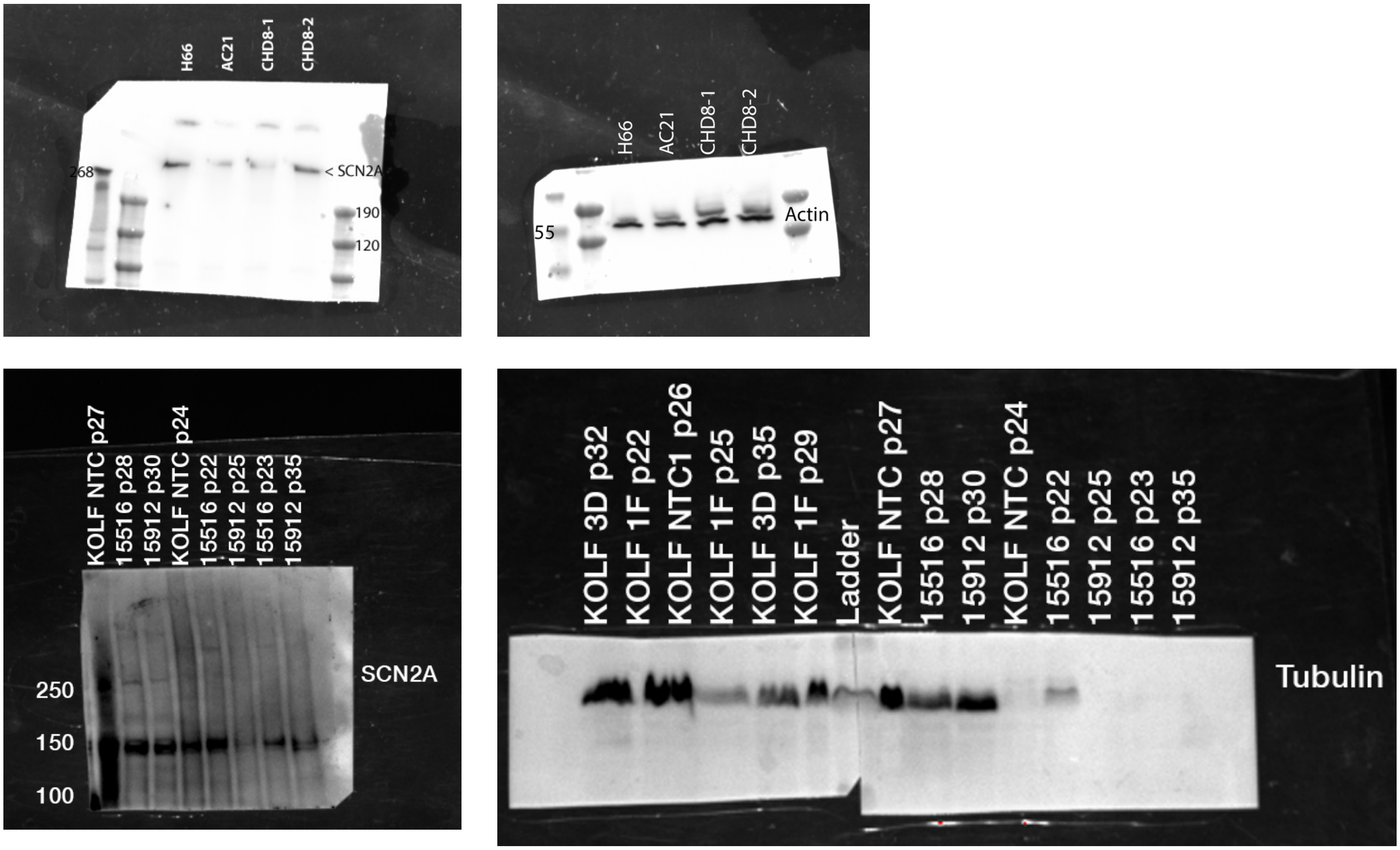

Figure 4. SCN2A

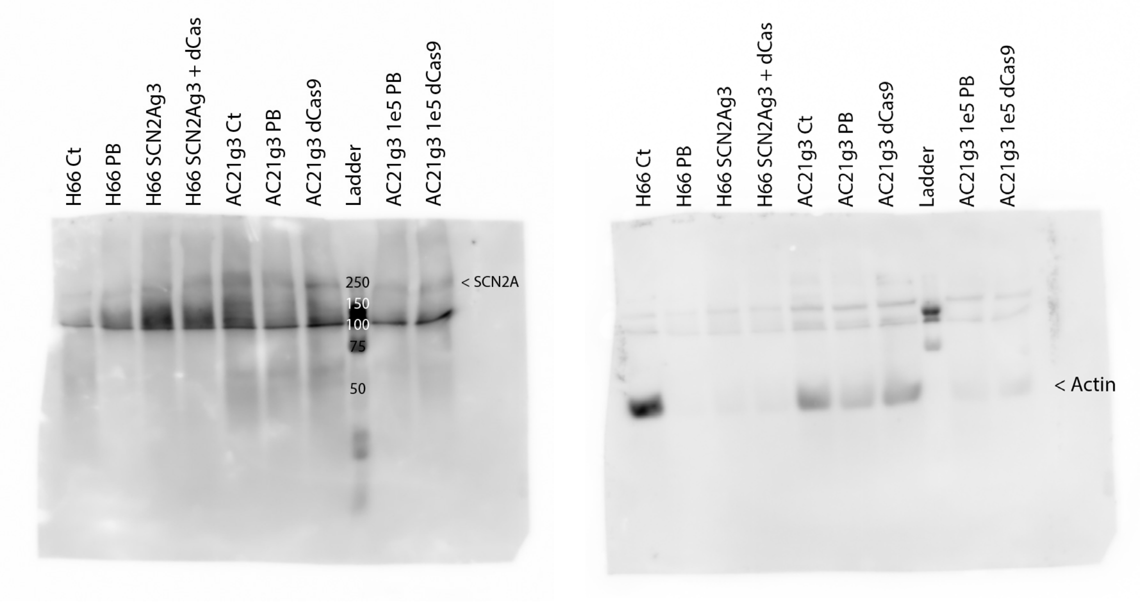

Figure 6. MAP2

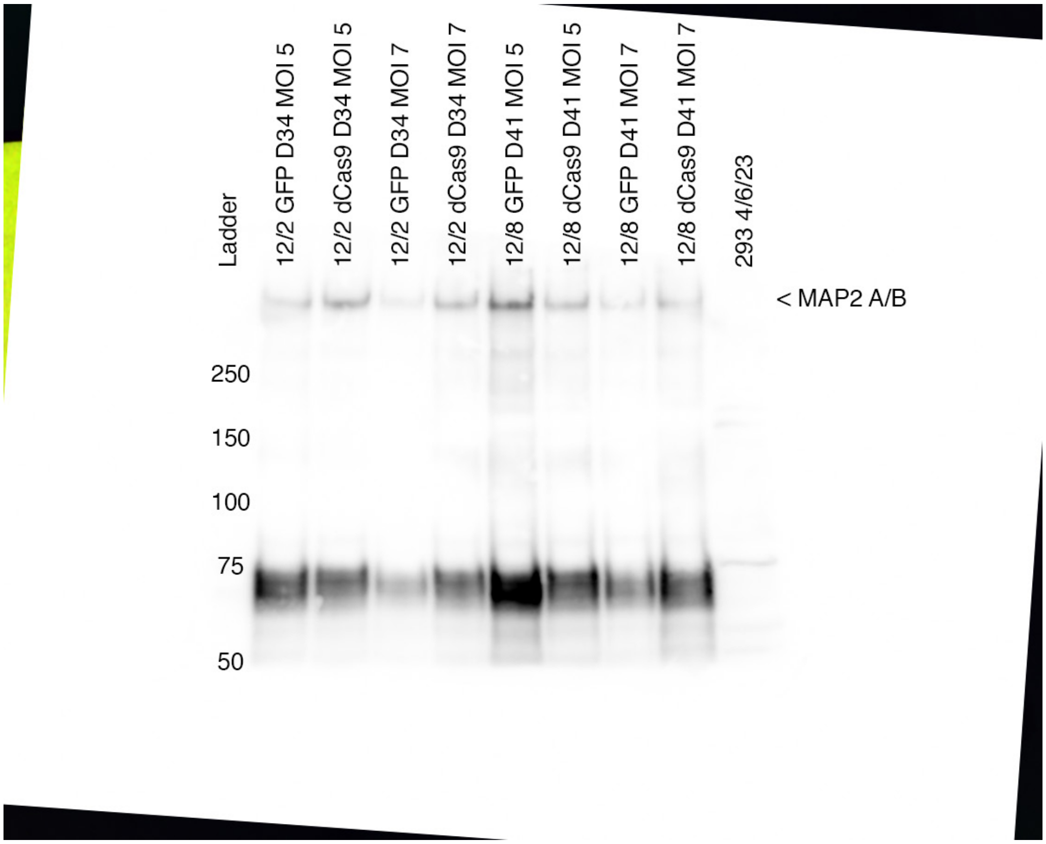


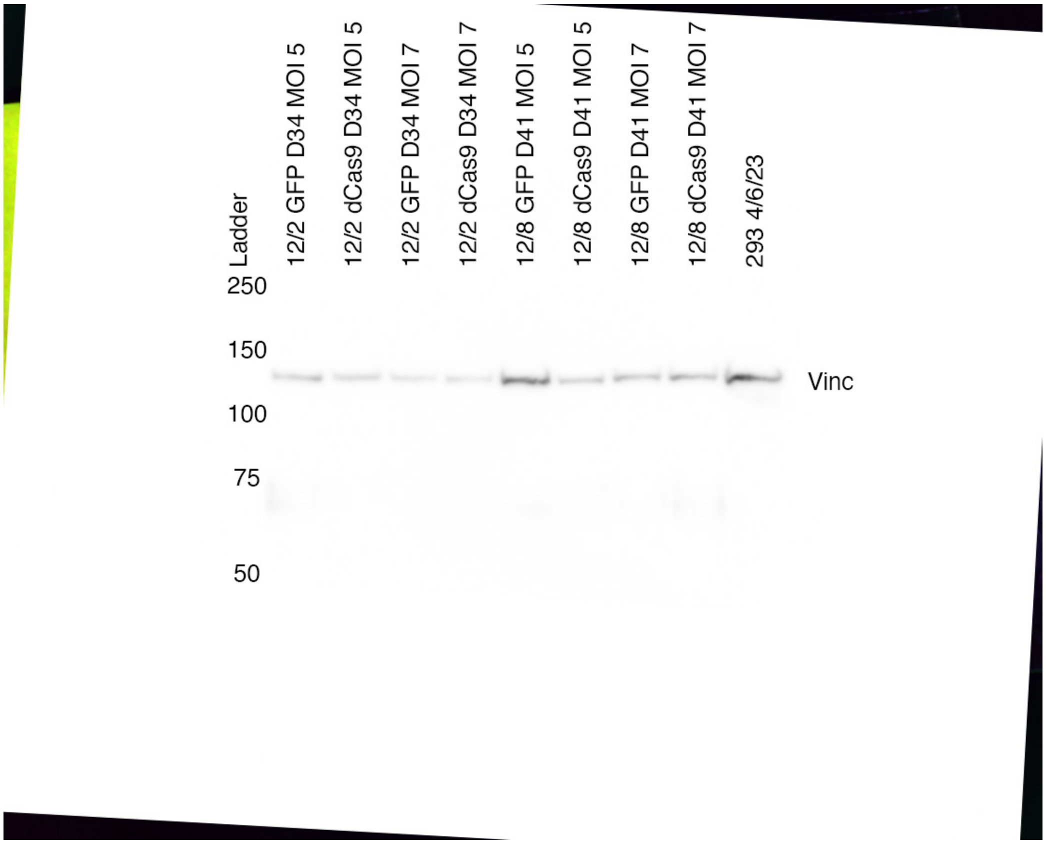

Figure 6. TUBB3

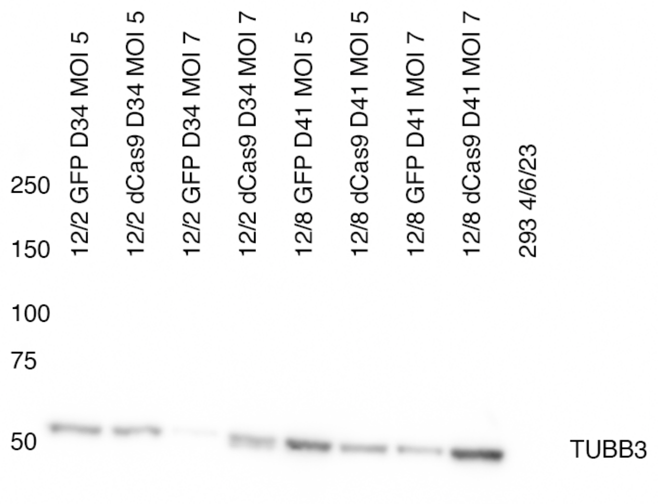


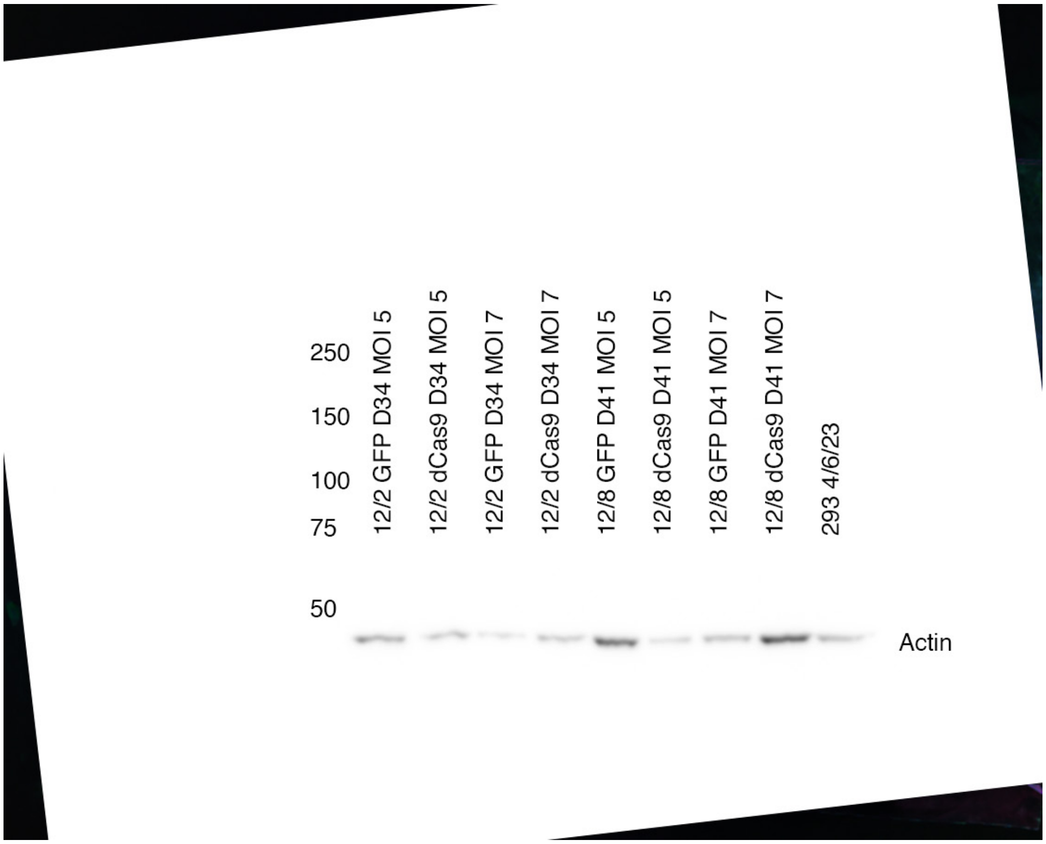

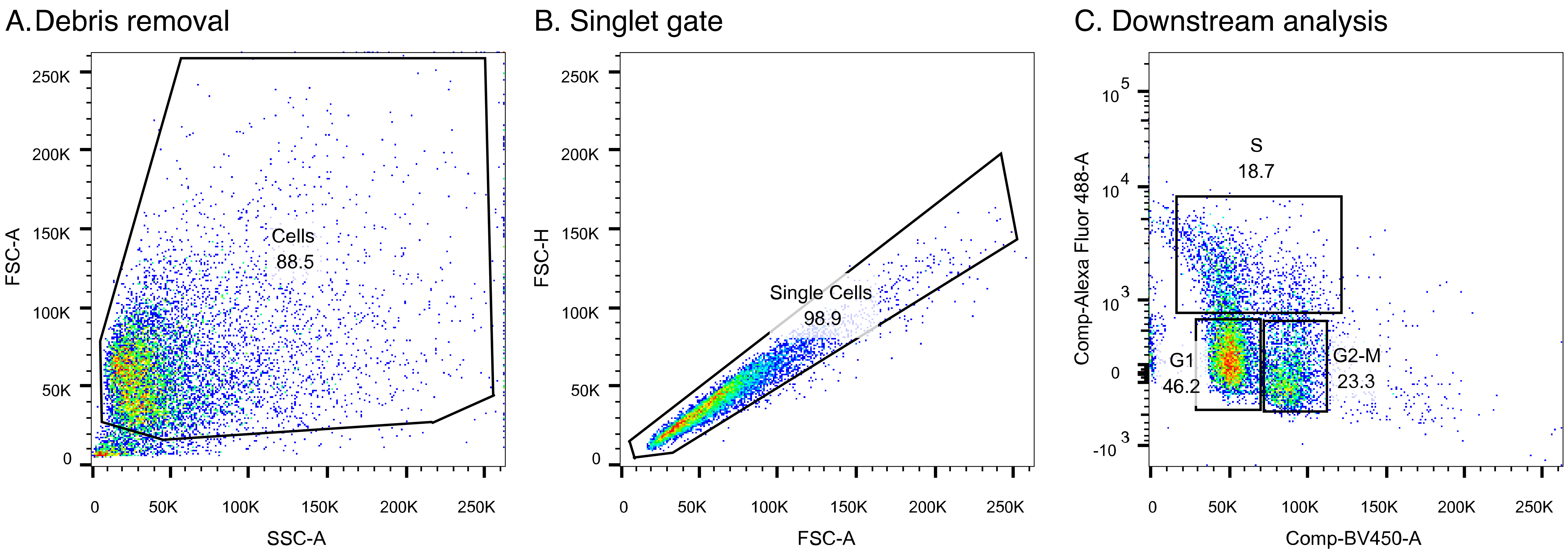
A.Debris removal
Cells
88.5
FSC-A
SSC-A
B. Singlet gate
Single Cells
98.9
FSC-H
FSC-A
C. Downstream analysis
S
18.7
G1
46.2
G2-M
23.3
Comp-Alexa Fluor 488-A
Comp-BV450-A

Figure 1F. CHD8 in HUES66

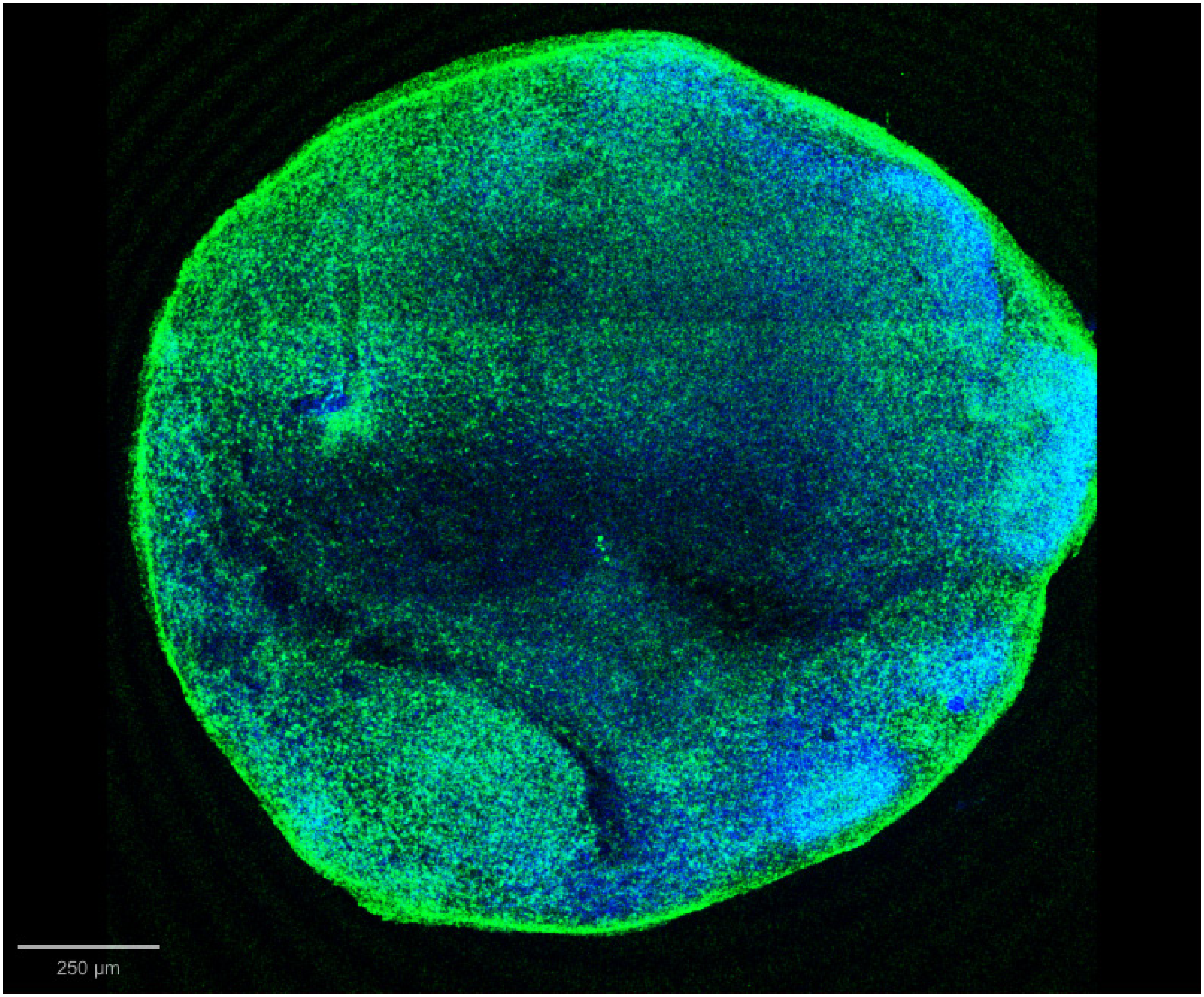

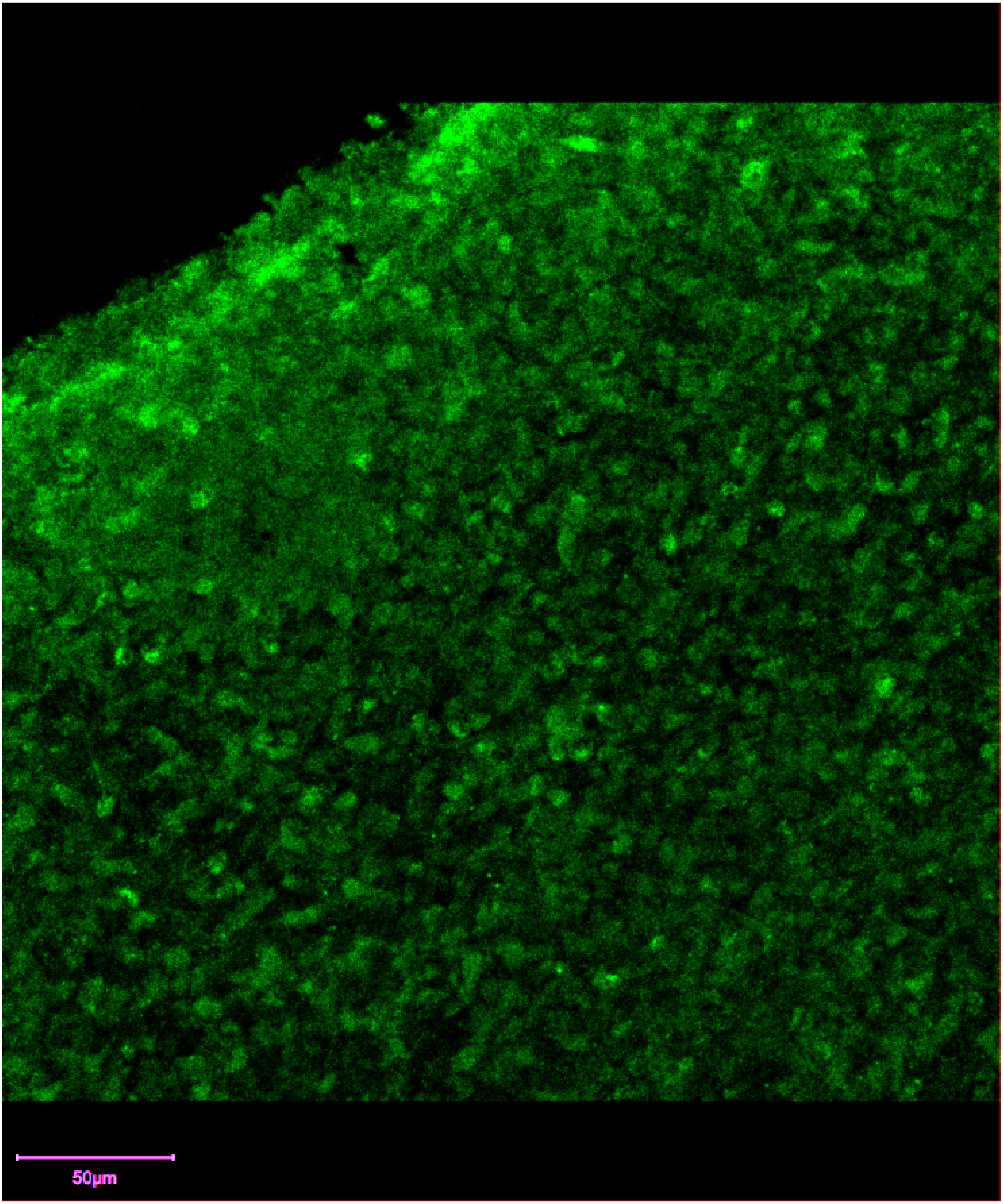
50µm

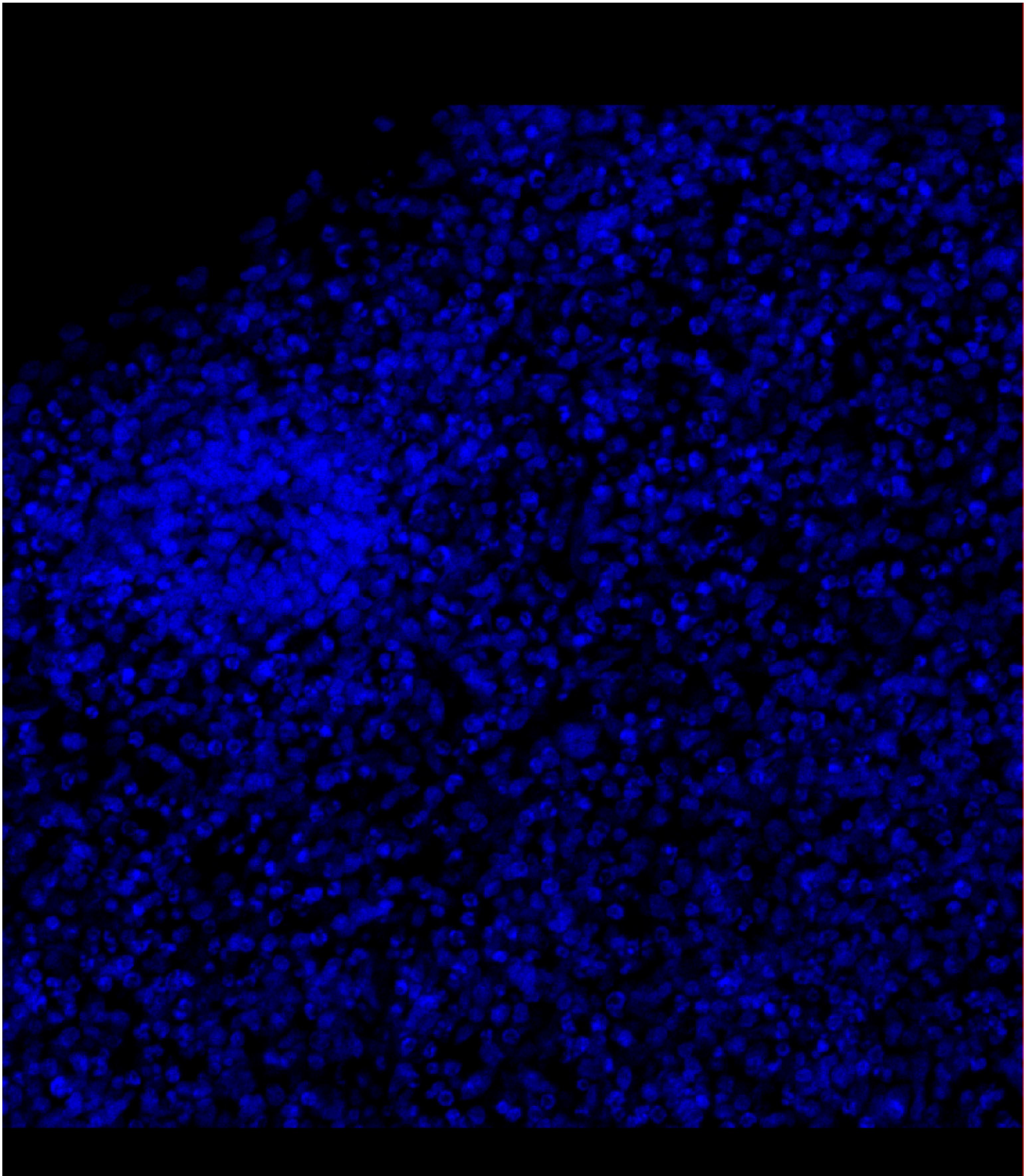
50µm

Figure 1F. CHD8 in CHD8-1A

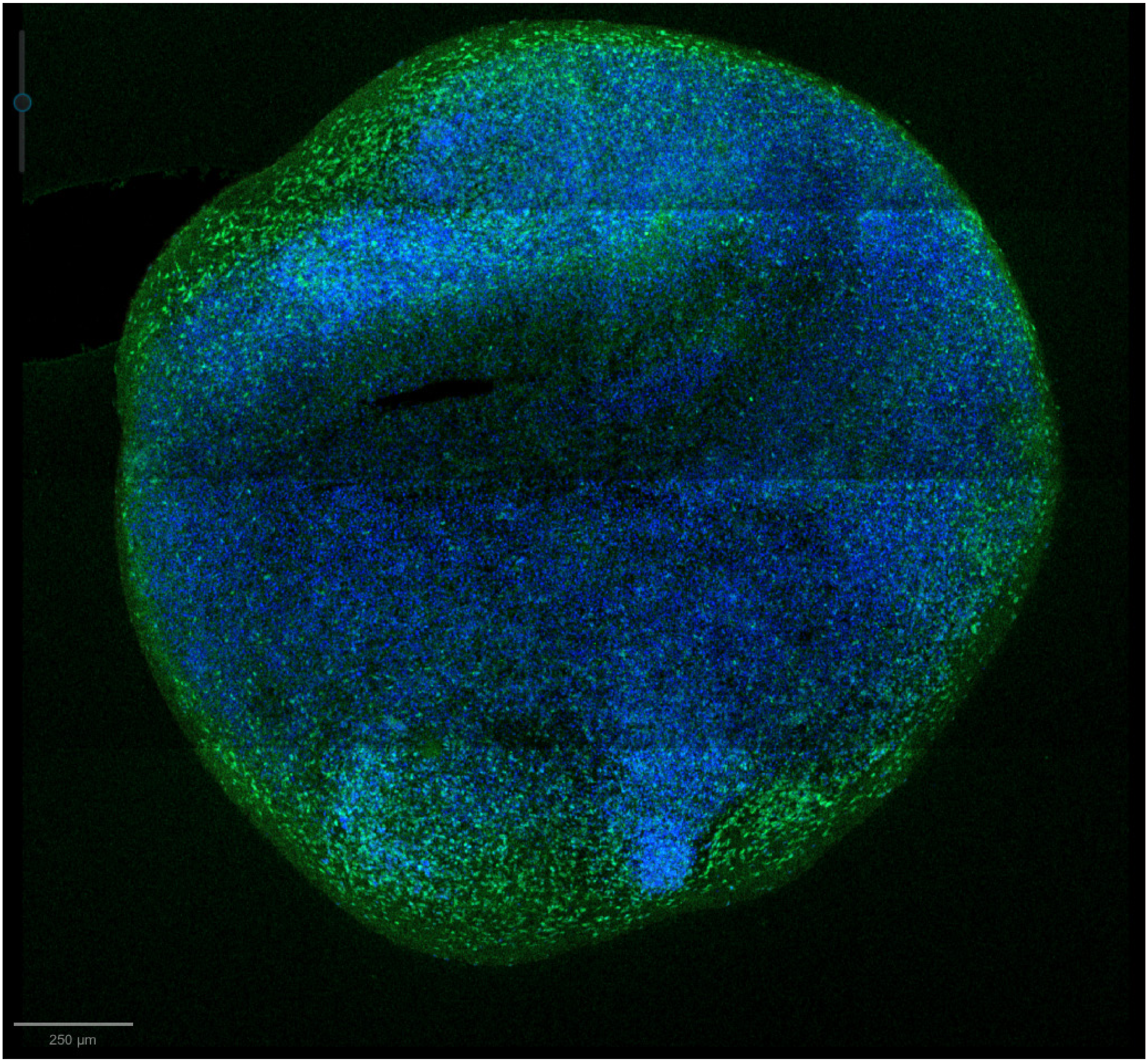

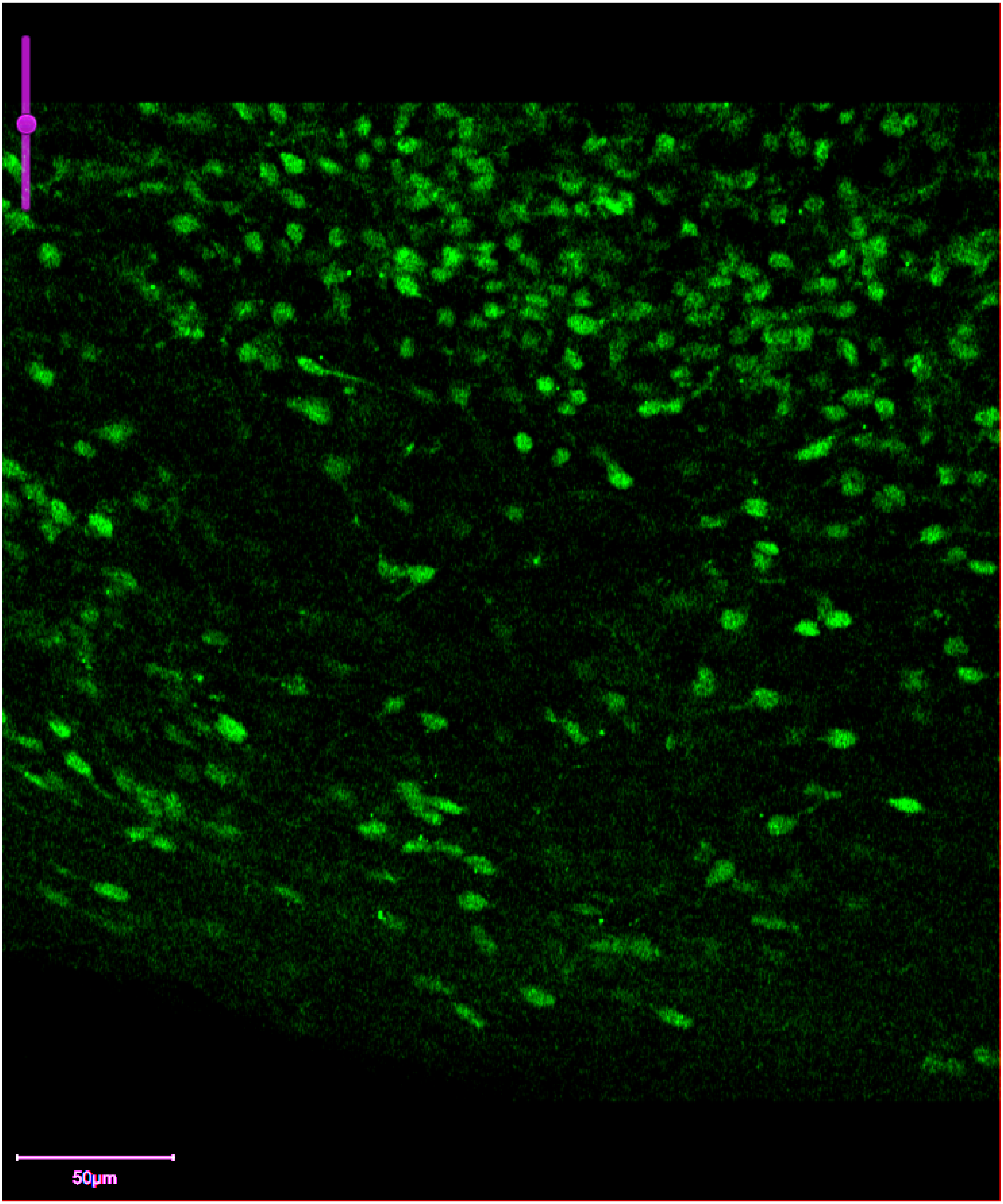
50μm

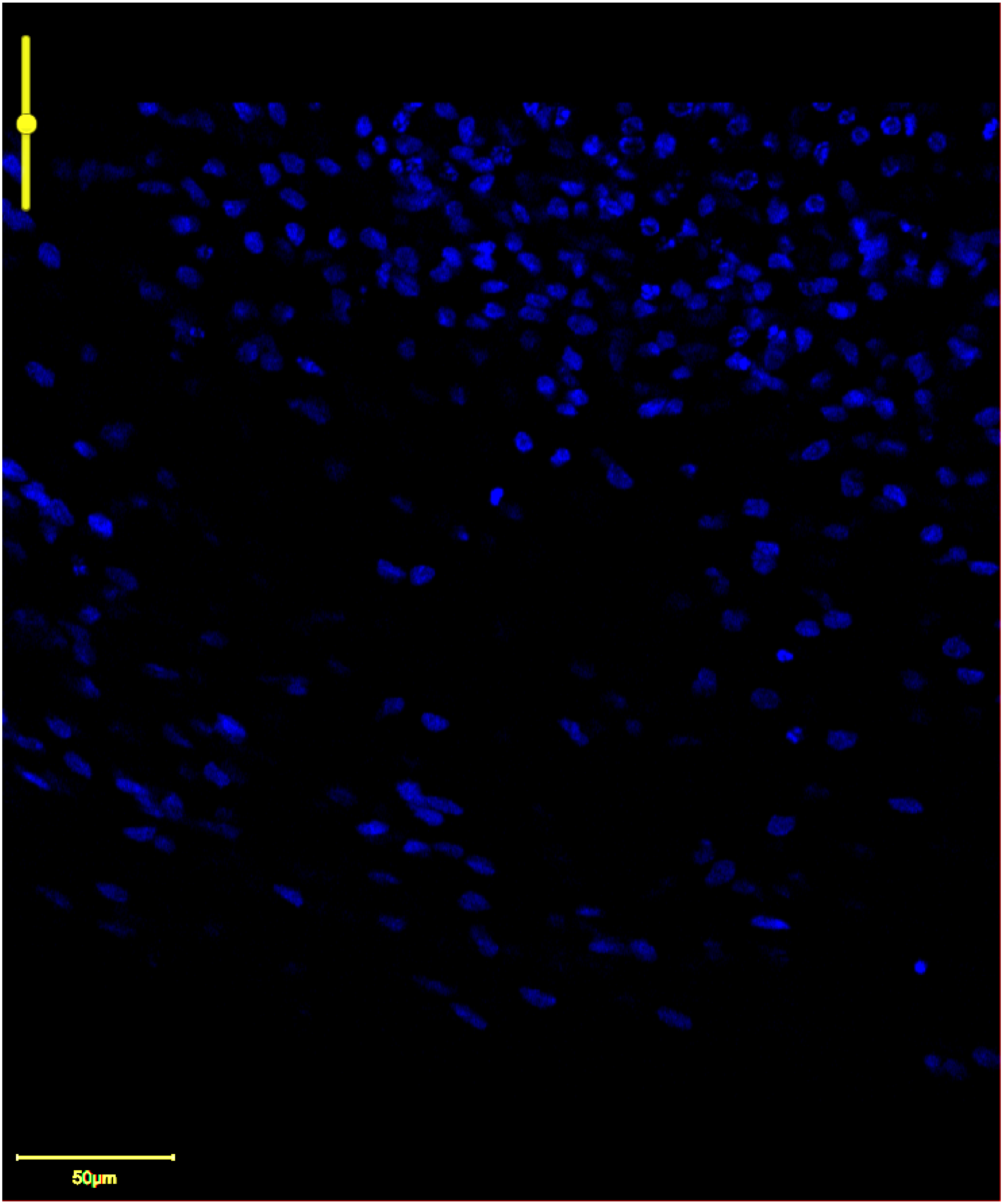
50μm

Figure 1F. CHD8 in CHD8-1B

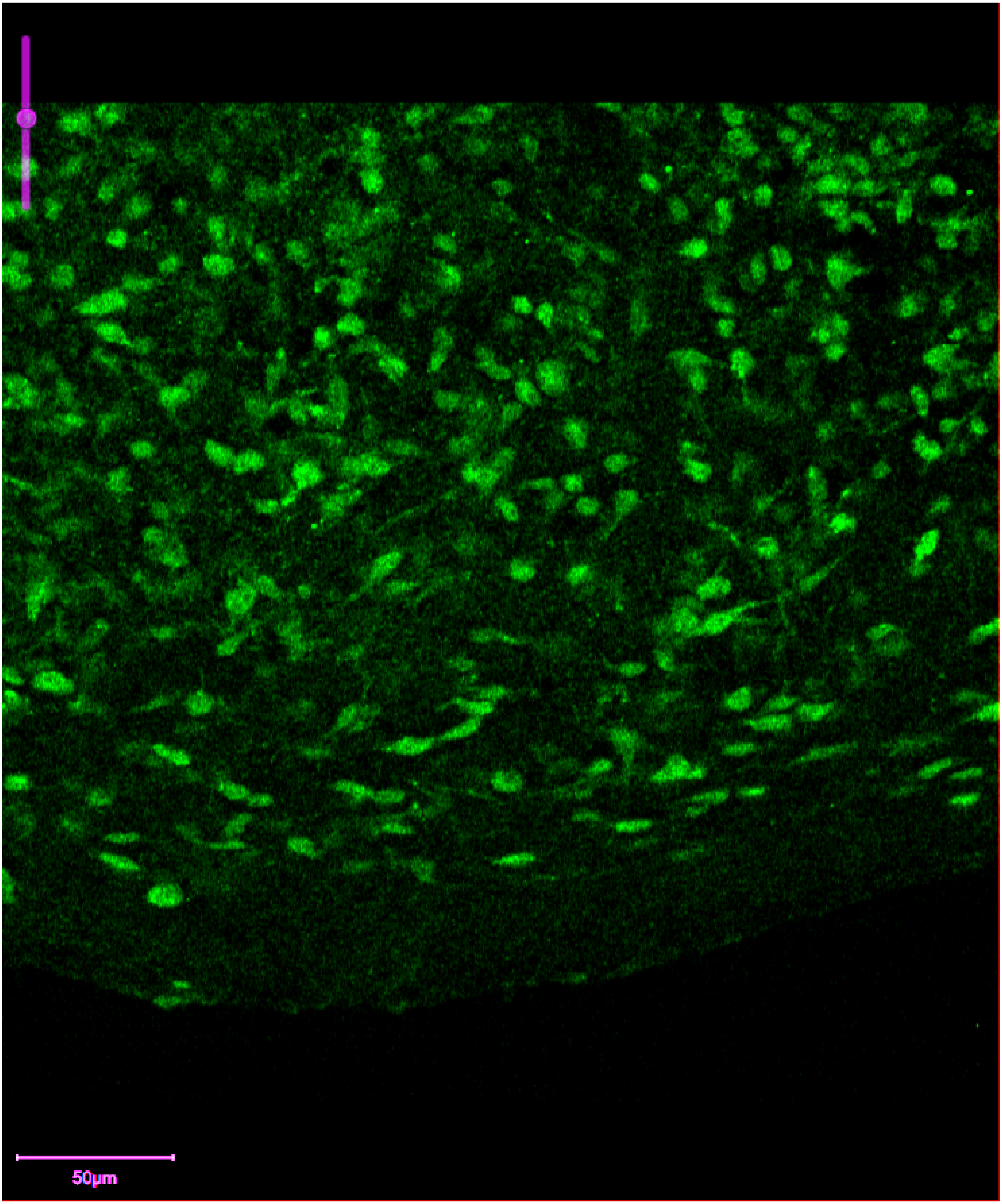

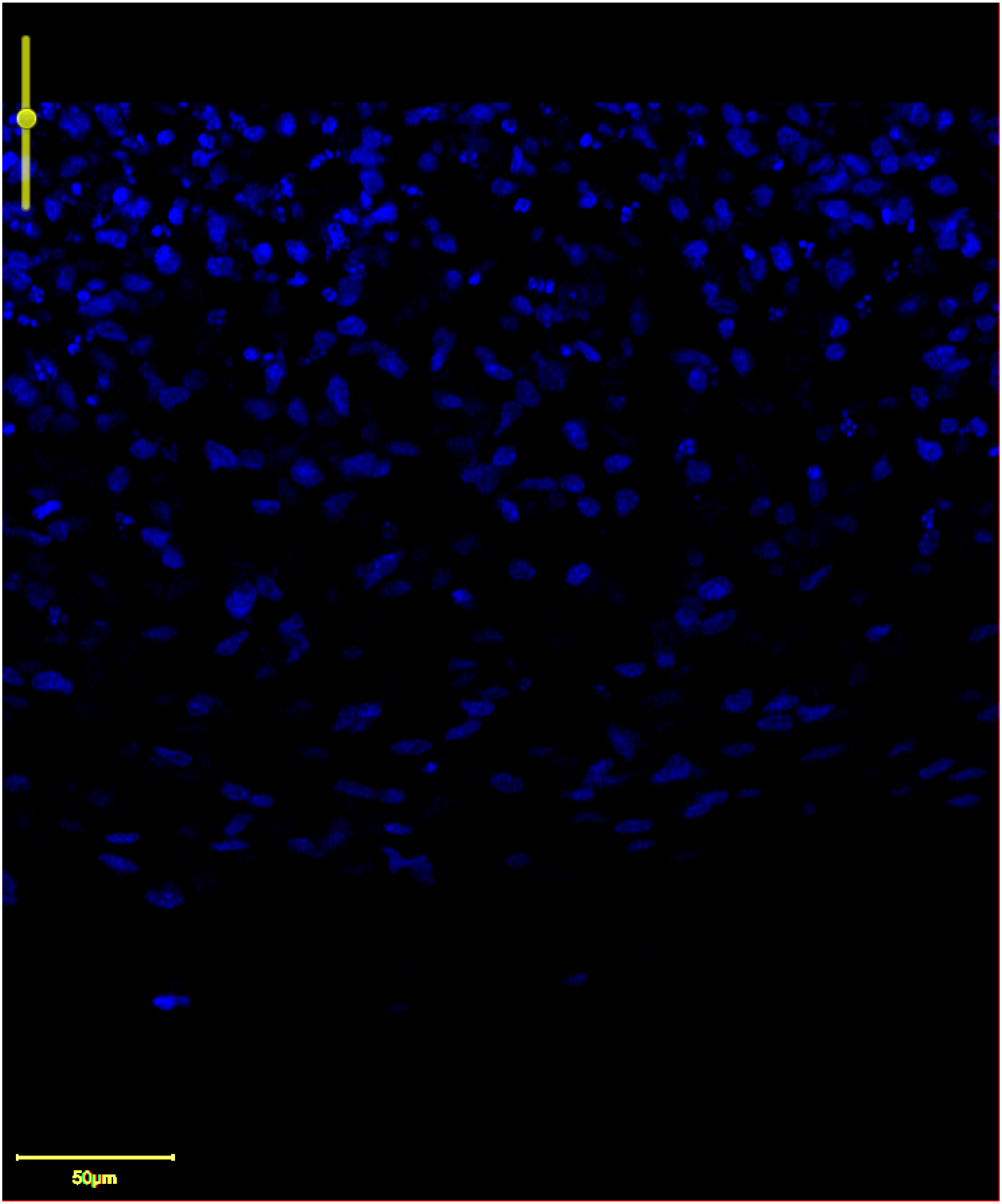
50μm

Figure 2E. SCN2A in HUES66

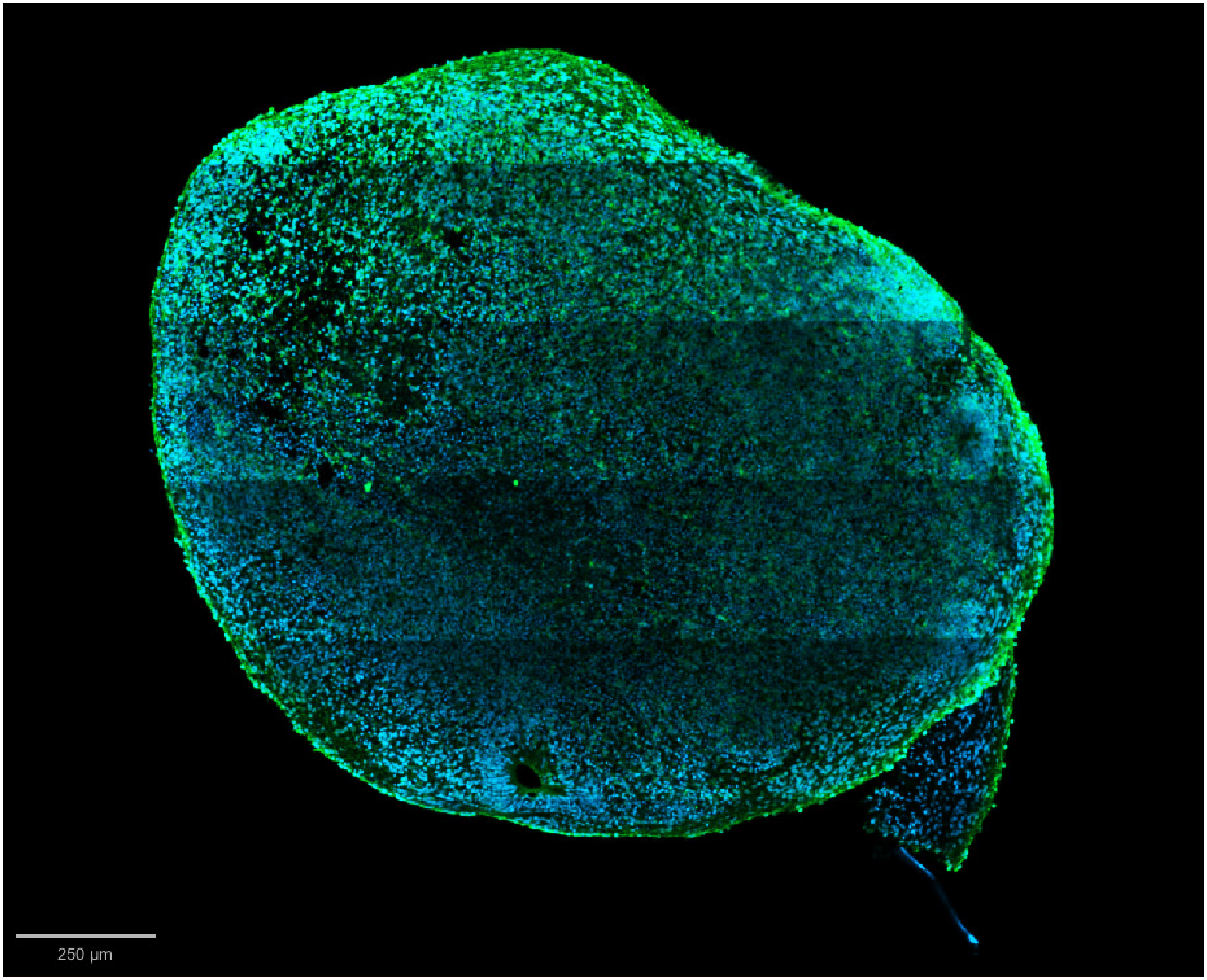

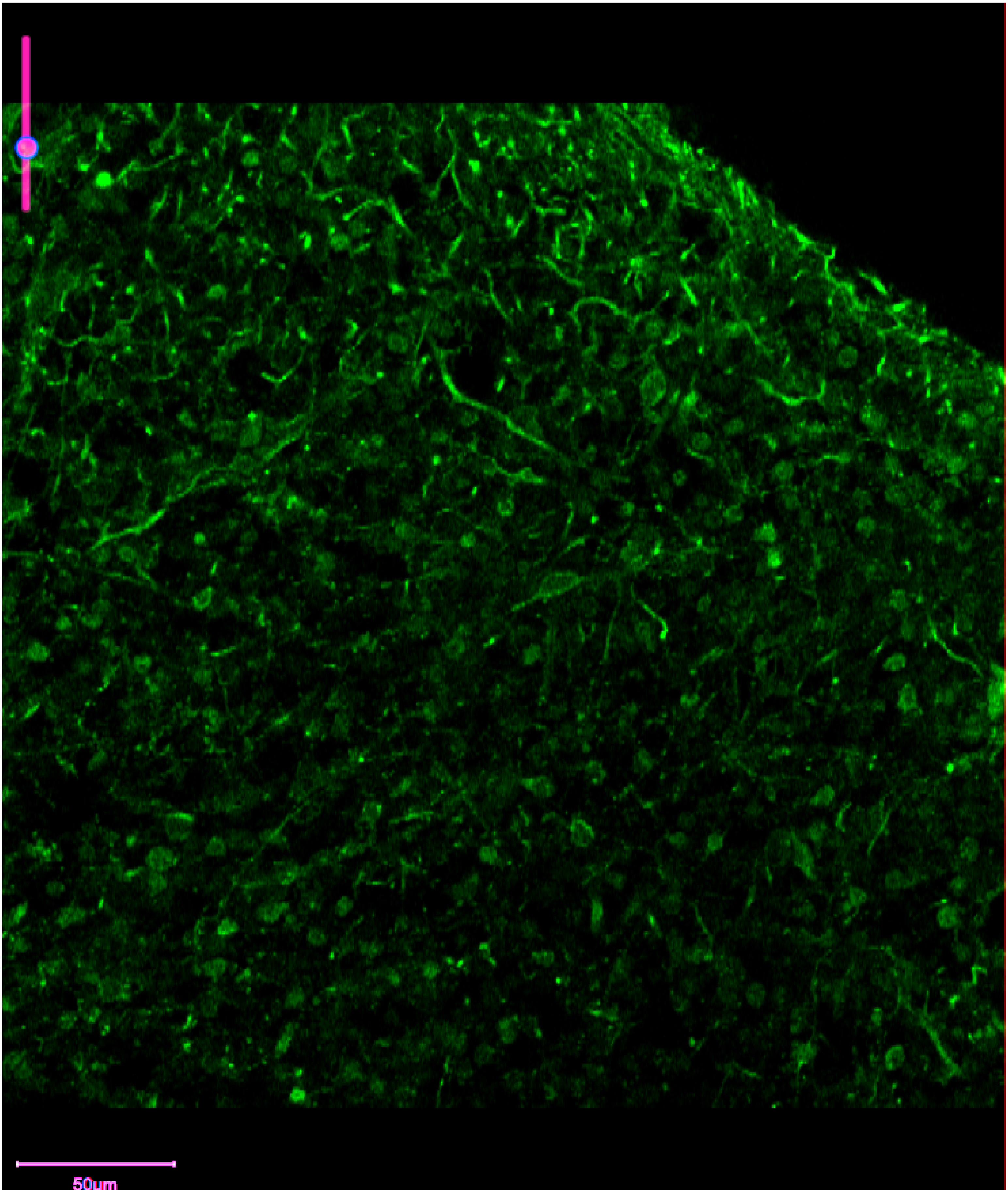
50µm

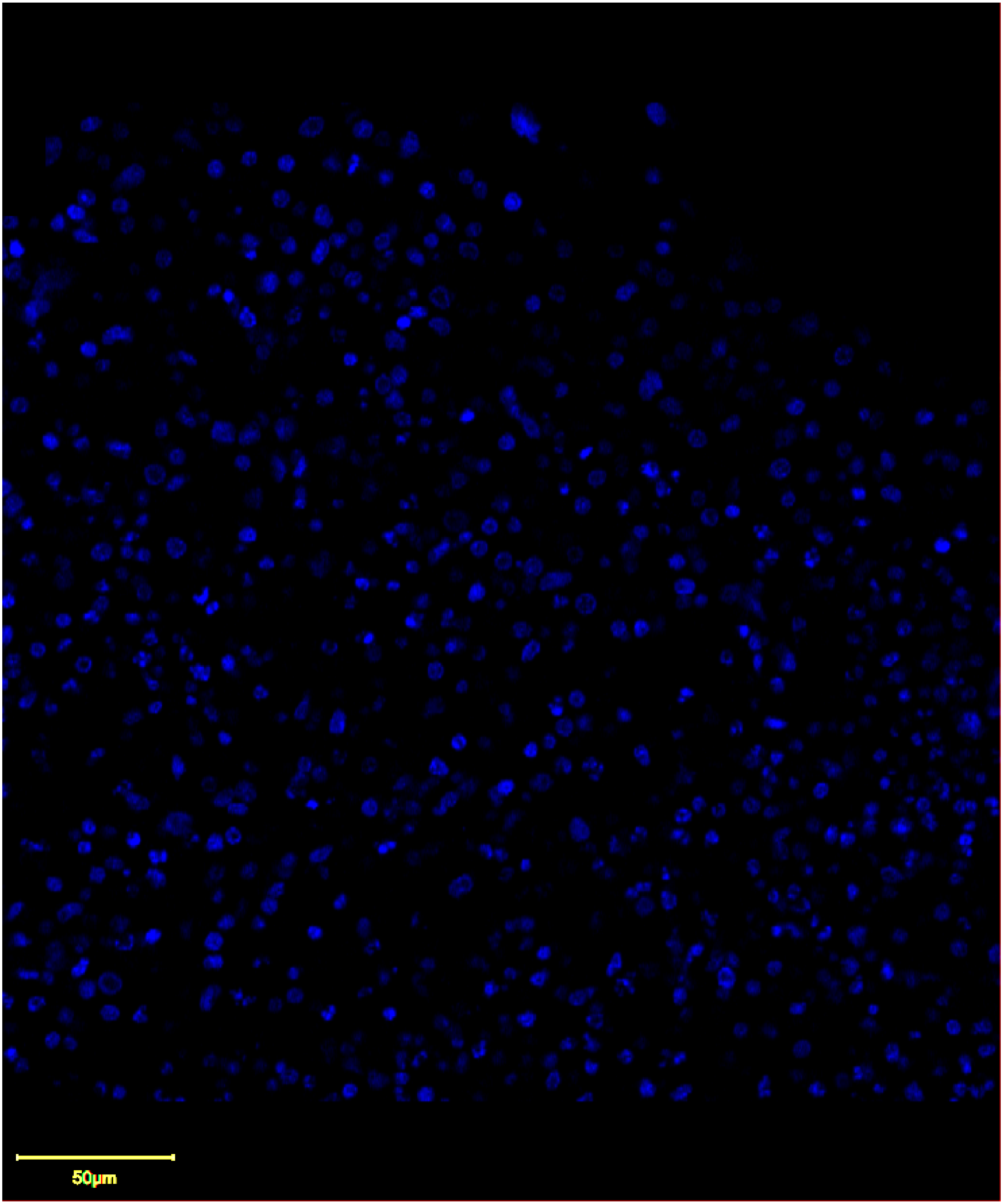
50μm

Figure 2E. SCN2A in SCN2A-1

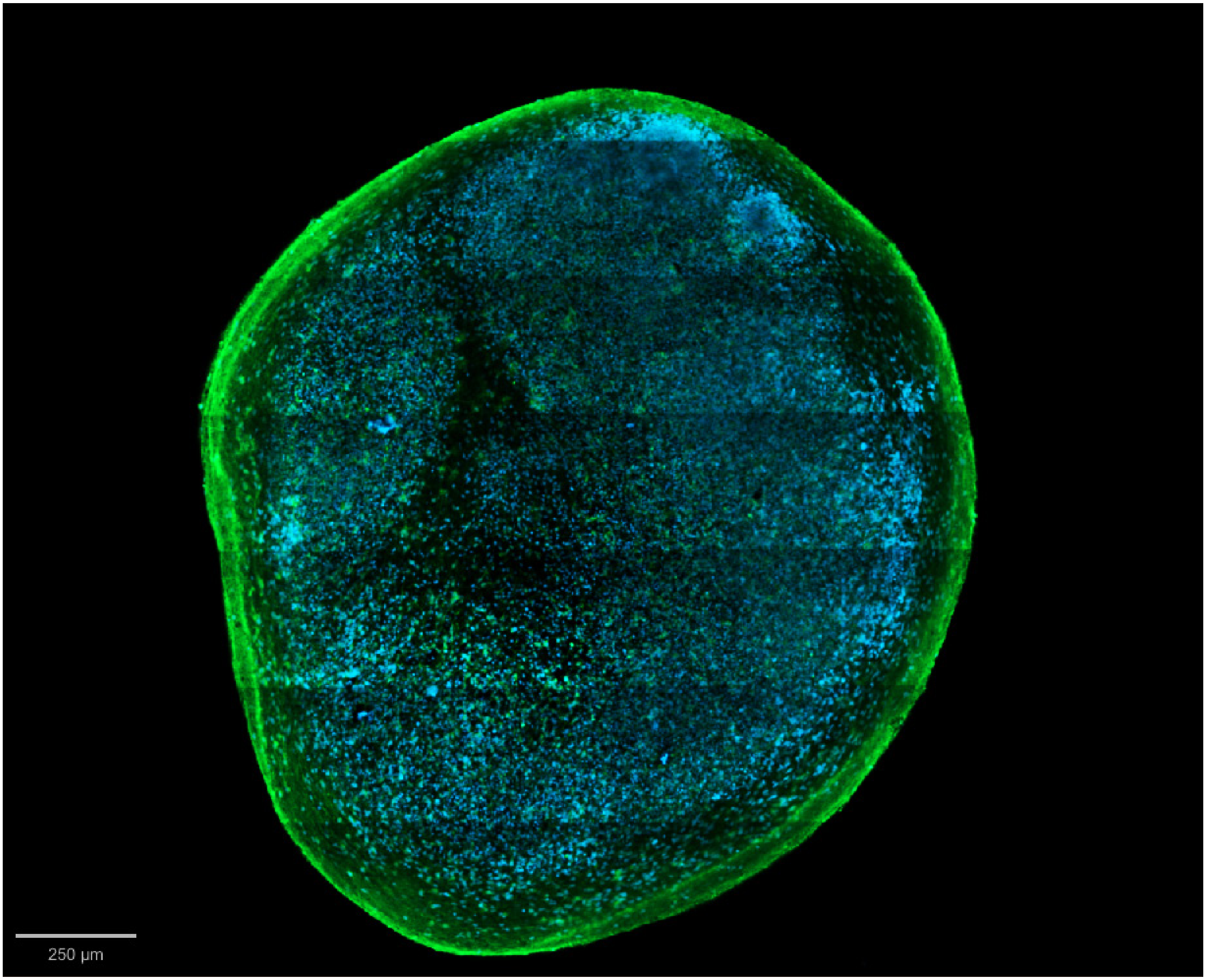

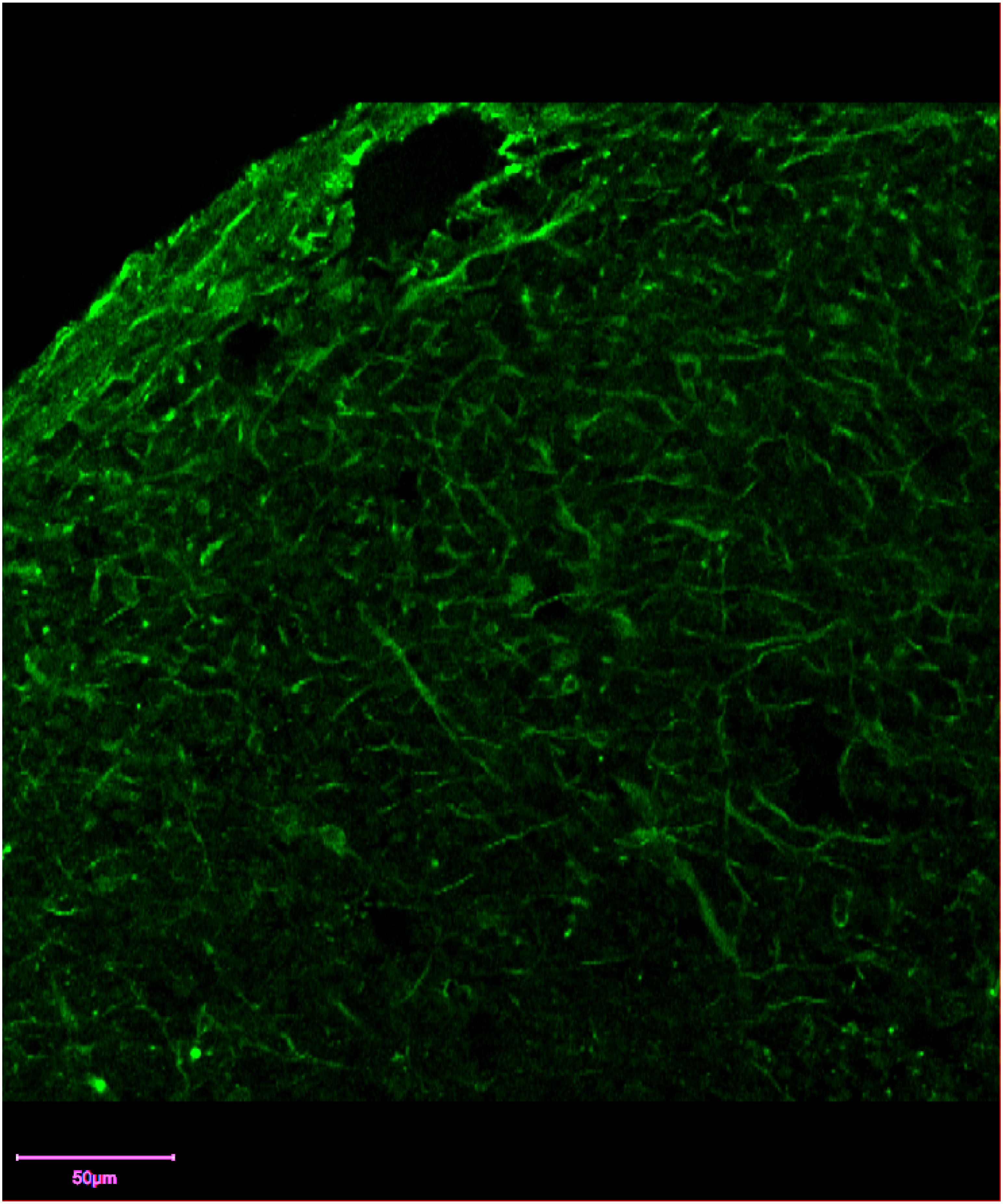
50µm

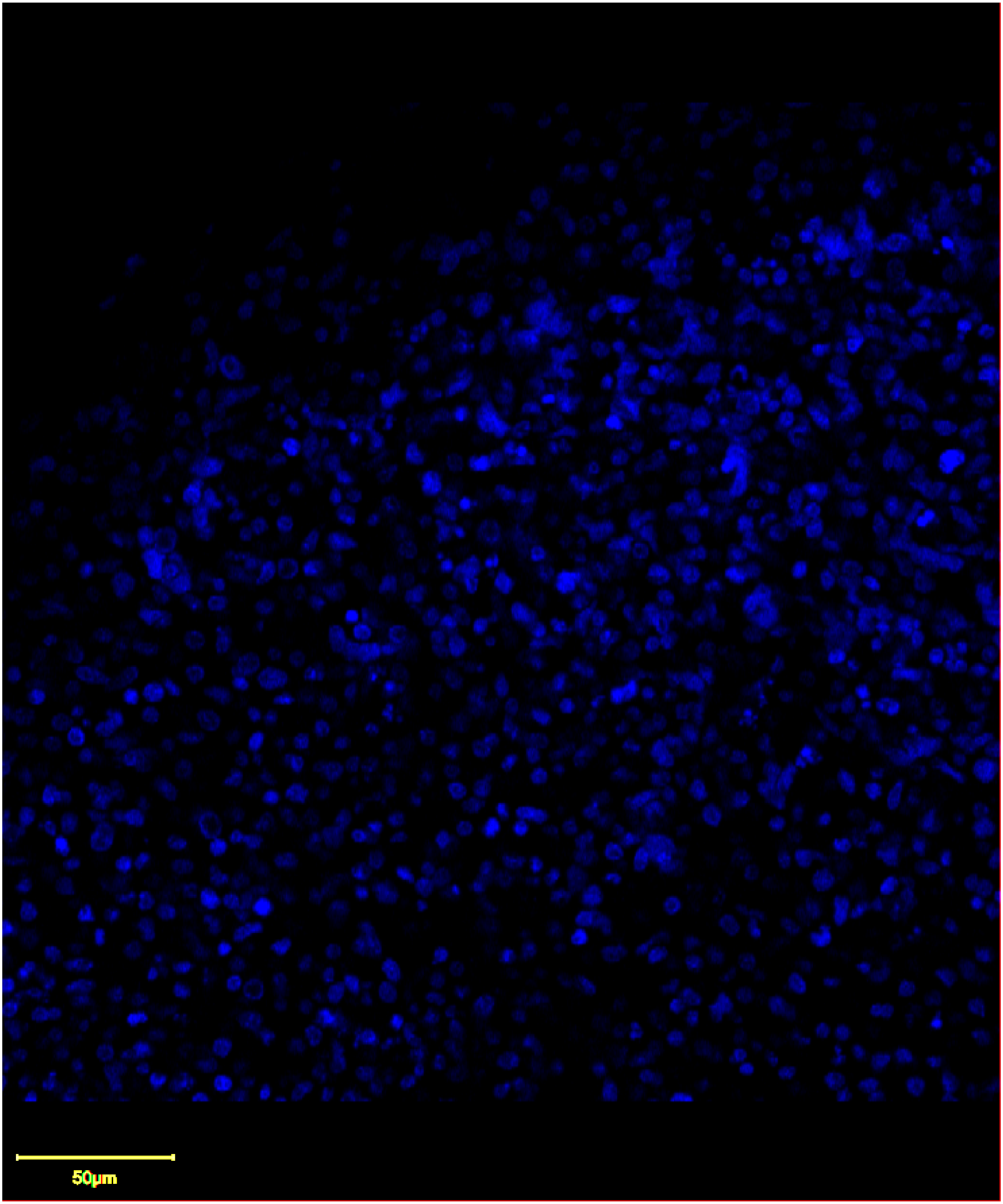
50μm

Figure 5A

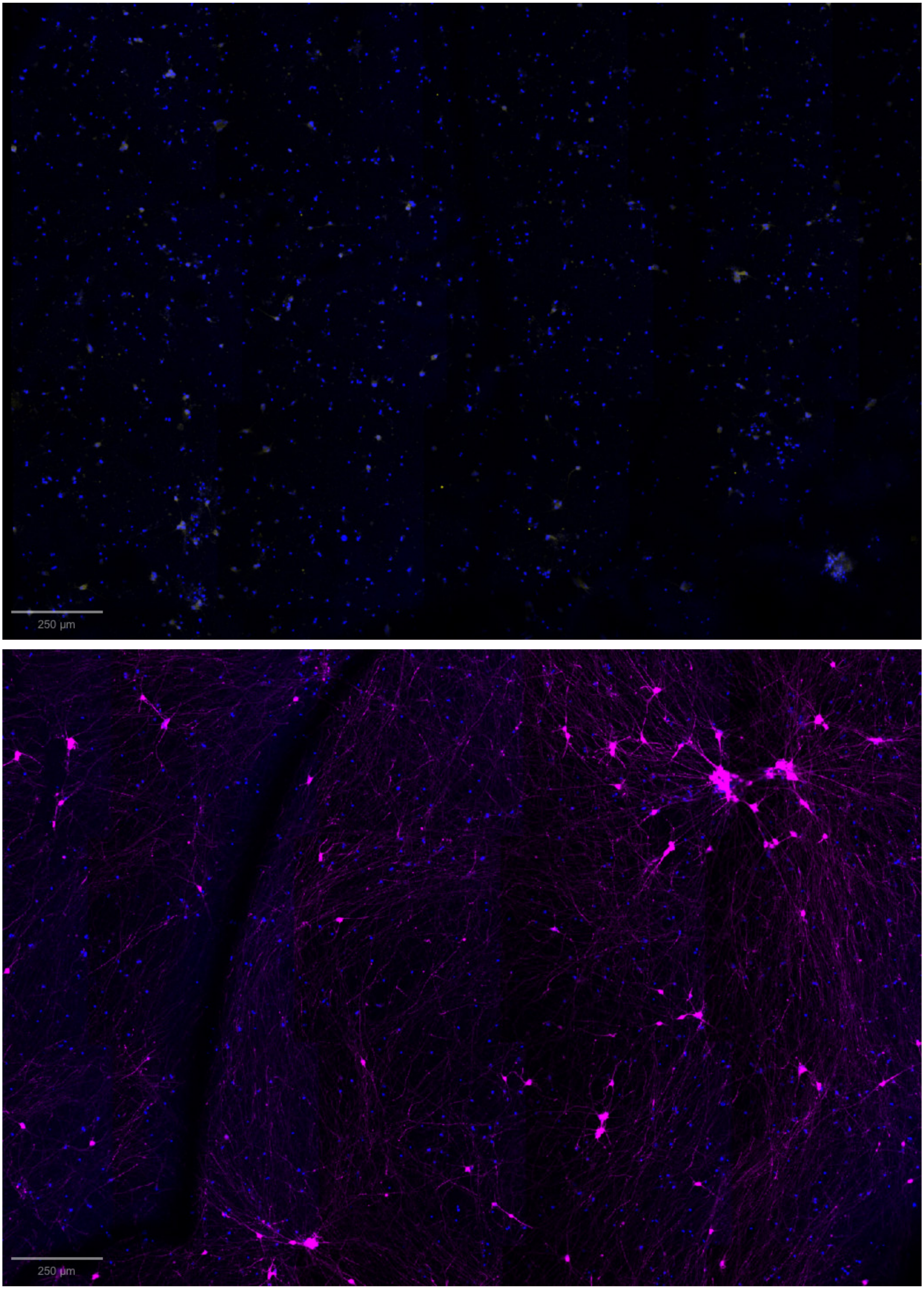

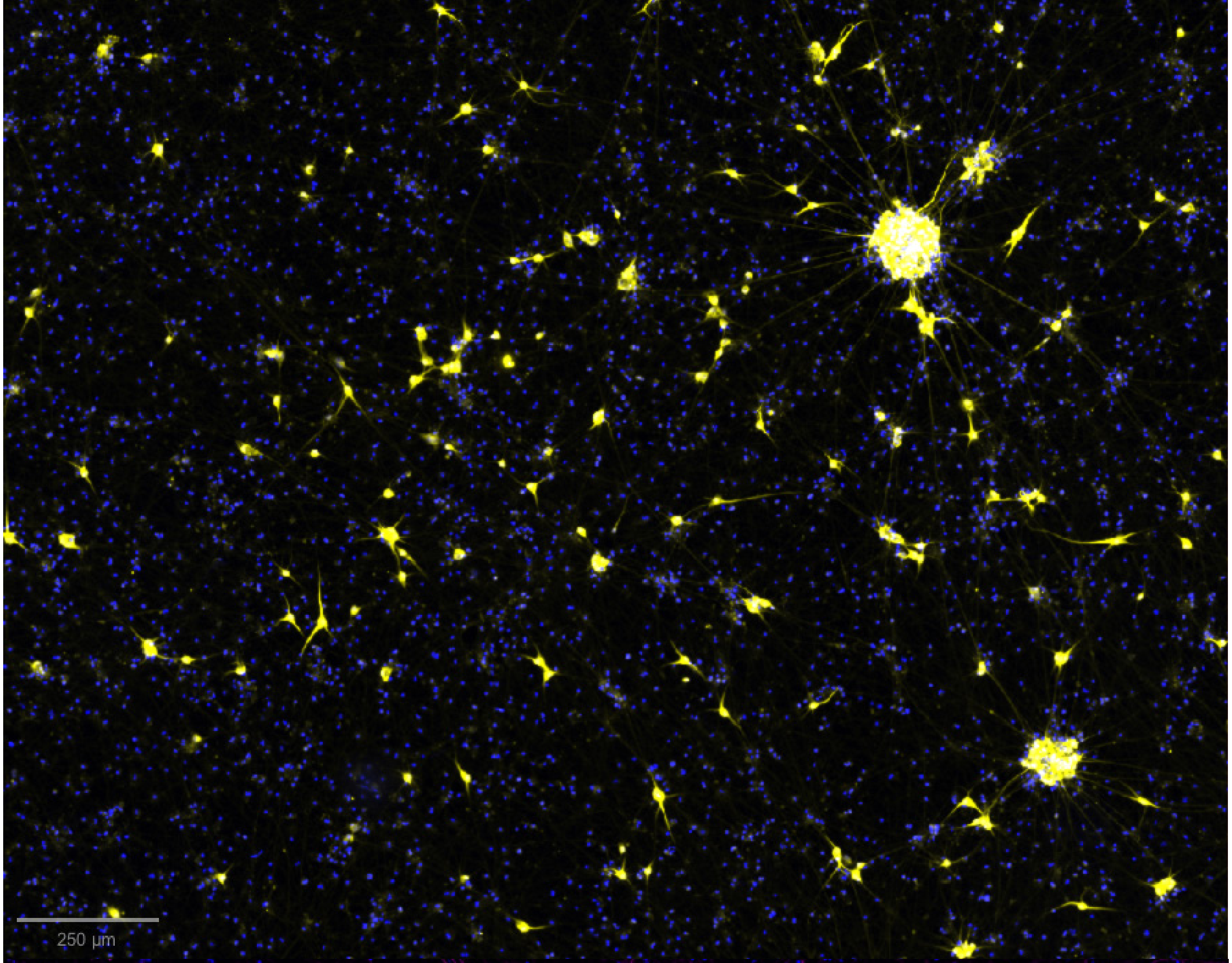
250 μm

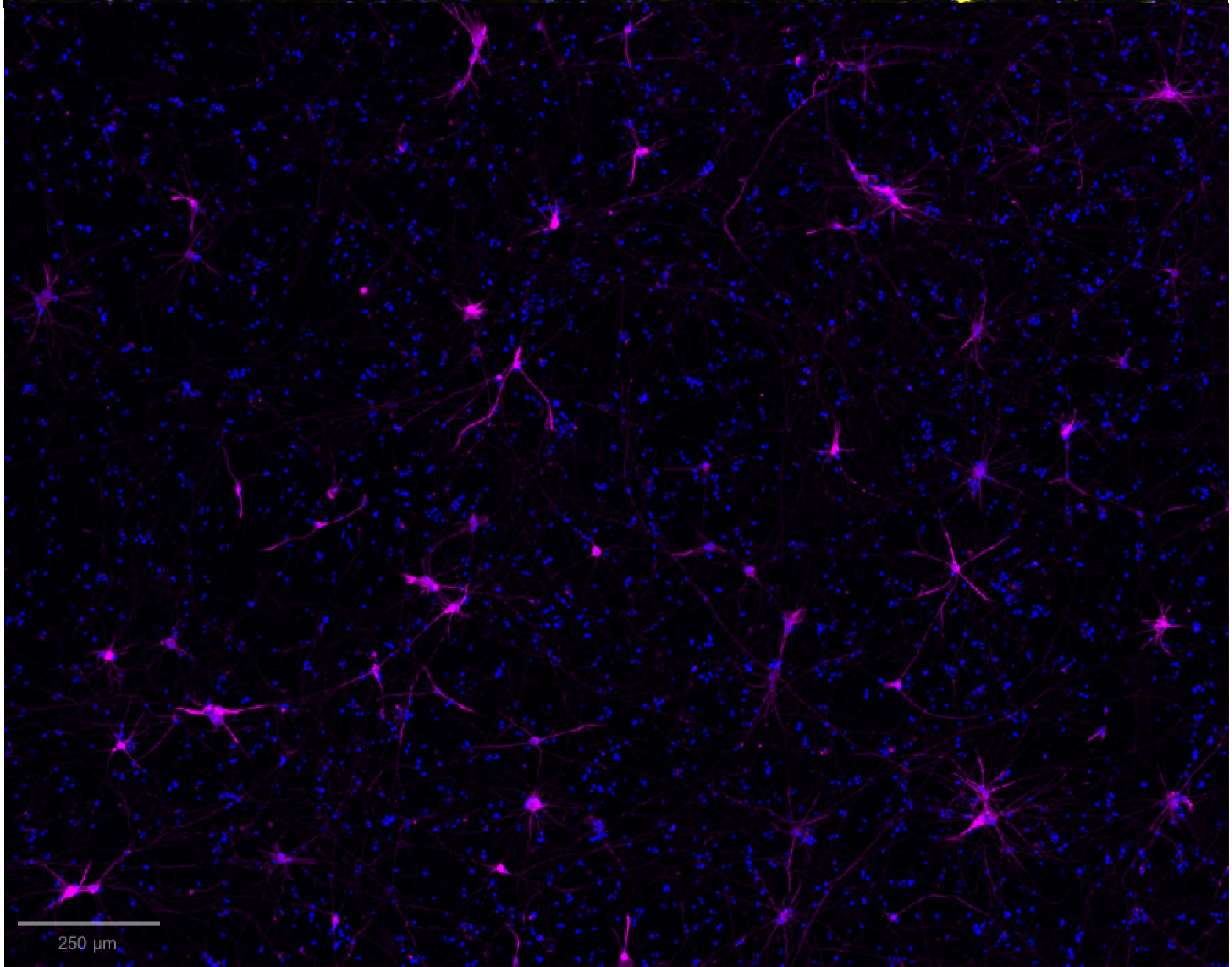
250 μm

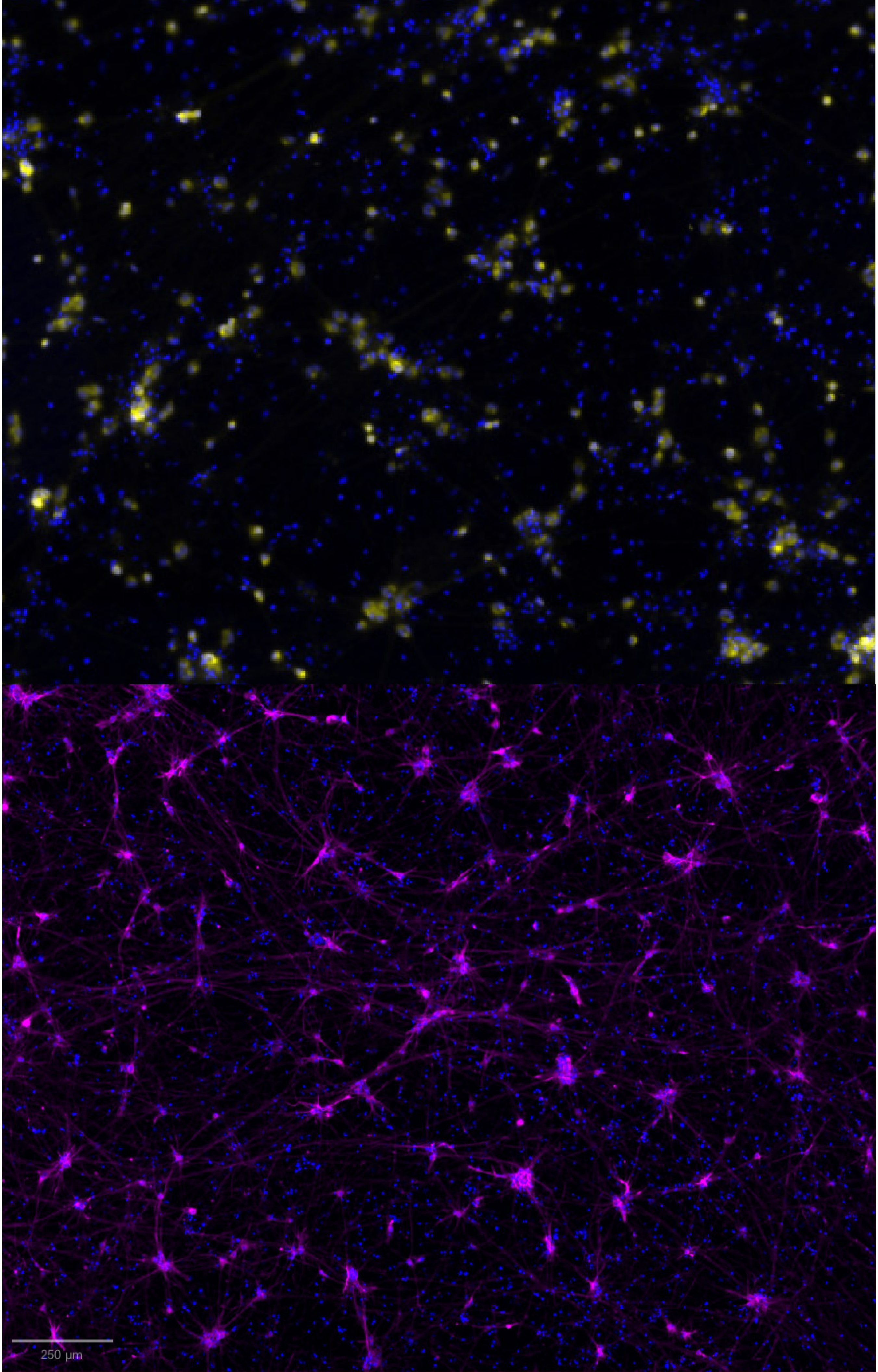
250 µm
